# Ultra-rare constrained missense variants in the epilepsies: Shared and specific enrichment patterns in neuronal gene-sets

**DOI:** 10.1101/2021.04.18.440264

**Authors:** Mahmoud Koko, Roland Krause, Thomas Sander, Dheeraj Reddy Bobbili, Michael Nothnagel, Patrick May, Holger Lerche, Epi25 Collaborative

**Affiliations:** Department of Neurology and Epileptology, Hertie Institute for Clinical Brain Research, University of Tübingen, Otfried-Müller Str. 25, 72076 Tübingen, Germany; Luxembourg Centre for Systems Biomedicine, University of Luxembourg, 6 Avenue du Swing, 4367 Belvaux, Luxembourg; University of Cologne, Faculty of Medicine and Cologne University Hospital, Cologne Center for Genomics, Weyertal 115b, 50931 Cologne, Germany; University of Cologne, Faculty of Mathematics and Natural Sciences and Cologne University Hospital, Cologne Center for Genomics, Weyertal 115b, 50931 Cologne, Germany

**Author notes:** See contributing authors in the appendix. Contributed equally as PIs (Patrick May, Holger Lerche). Correspondence to: Prof. Holger Lerche. Address: Department of Neurology and Epileptology, Hertie Institute for Clinical Brain Research, University and University Clinic Tübingen, Hoppe-Seyler-Str. 3, 72076 Tübingen, Germany. E-Mail or Dr. Patrick May. Address: Luxembourg Centre for Systems Biomedicine, University of Luxembourg, 6 Avenue du Swing, 4367 Belvaux, Luxembourg.

**Keywords:** burden analysis, ultra-rare variants, gene-sets, epilepsy, exome sequencing

## Abstract

**Background:** Burden analysis in epilepsy has shown an excess of deleterious ultra-rare variants (URVs) in few gene-sets, such as known epilepsy genes, constrained genes, ion channel or GABA_A_ receptor genes. We set out to investigate the burden of URVs in a comprehensive range of gene-sets presumed to be implicated in epileptogenesis.

**Methods:** We investigated several constraint and conservation-based strategies to study whole exome sequencing data from European individuals with developmental and epileptic encephalopathies (DEE, *n* = 1,003), genetic generalized epilepsy (GGE, *n* = 3,064), and non-acquired focal epilepsy (NAFE, *n* = 3,522), collected by the Epi25 Collaborative, compared to 3,962 ancestry-matched controls. The burden of 12 non-synonymous URVs types in 92 gene-sets was compared between epilepsy cases (DDE, GGE, NAFE) and controls using logistic regression analysis.

**Results:** Burden analysis of brain-expressed genes revealed an excess of different URVs types in all three epilepsy categories which was largest for missense variants in highly constrained sites (constrained missense variants). The URVs burden was prominent in neuron-specific, synaptic and developmental genes as well as genes encoding ion channels and receptors, and it was generally higher for DEE and GGE compared to NAFE. The patterns of URVs burden in gene-sets expressed in inhibitory vs. excitatory neurons or receptors suggested a high burden in both groups in DEE but a differential involvement of inhibitory genes in GGE, while excitatory genes were predominantly affected in NAFE. Top ranking susceptibility genes from a recent genome-wide association study (GWAS) of generalized and focal epilepsies displayed a higher URVs burden in constrained coding regions in GGE and NAFE, respectively.

**Conclusions:** Using exome-based gene-set burden analysis, we demonstrate that missense URVs affecting mainly constrained sites are enriched in neuronal genes in both common and rare severe epilepsy syndromes. Our results indicate a differential impact of these URVs in genes expressed in inhibitory vs. excitatory neurons and receptors in generalized vs. focal epilepsies. The excess of URVs in top-ranking GWAS risk-genes suggests a convergence of rare deleterious and common risk-variants in the pathogenesis of generalized and focal epilepsies.

## Background

Dismantling the genetic architecture behind epilepsy is yet to be within reach in many individuals. The role of genetic causality is apparent in the developmental and epileptic encephalopathies (DEE) (1–3), sometimes with consequences on precision treatments (4–7). In contrast, only few individuals with familial or sporadic genetic generalized epilepsies (GGE) or non-acquired focal epilepsies (NAFE) harbor monogenic causative variations (8–10). Therefore, statistical methods investigating the mutational burden of neurobiologically meaningful gene-sets improve the prospects to dissect the joint effects of multiple genetic factors underlying the complex genetic architecture of these common epilepsy syndromes. Such ‘gene-set’ analysis approaches are likely to provide valuable insights into the role of certain gene-sets and pathways in epilepsy. Recent gene-set burden analyses have shown an enrichment in ultra-rare deleterious variants both in common and rare epilepsies in genes associated with dominant epilepsy syndromes, developmental and epileptic encephalopathy genes, and neuro-developmental disorders (NDD) with epilepsy genes, emphasizing a shared genetic component (8,10). Evidence for the enrichment of rare missense variants in genes encoding GABA_A_ receptors and GABAergic pathway genes in genetic generalized epilepsies pointed to the importance of the inhibitory pathway (9,10). We used the large-scale dataset collected by the Epi25 Collaborative (10) for a comprehensive, exome-based case-control study to examine the burden of Ultra-Rare Variants (URVs) in a large number of candidate gene-sets for three different epilepsy forms (DEE, GGE, NAFE), aiming to understand the specific roles of deleterious URVs in key pathways implicated in epileptogenesis. Focusing on regional constraint and paralog conservation, we identified relevant and specific gene-set associations in these three epilepsy forms.

## Methods

### Study Samples

The Epi25 Collaborative collected and generated phenotyping and exome sequencing data from individuals with different subtypes of epilepsy (10). We analyzed subjects from recruitment years 1 and 2 (*n*=13,197). The epilepsy classification, phenotyping and consent procedures have been previously described (10). Five control cohorts, from the Epi25 Collaborative and the database of Genotypes and Phenotypes (dbGAP) (11), were available for this analysis (*n*=13,299), including Italian controls from the Epi25 Collaborative, the Swedish Schizophrenia Study controls (dbGAP: phs000473), and three Myocardial Infarction Genetics (MIGen) Consortium cohorts: Leicester UK Heart Study (dbGAP: phs001000), Ottawa Heart Study (dbGAP: phs000806) and the Italian Atherosclerosis, Thrombosis, and Vascular Biology (ATVB) Study (dbGAP: phs001592). Sequencing was performed on an Illumina HiSeq 2000 or 2500 platform at the Broad Institute (Cambridge, MA, USA). The data generation process has been previously described (10).

### Baseline sample quality control

For this analysis, we considered samples from three epilepsy categories (DEE, GGE, and NAFE) as classified by the Epi25 phenotype review (10). Controls with Coronary Artery Disease (CAD) diagnosis from the MIGen cohorts were not considered. Samples with genotyping rate less than 90%, outliers on key sample calling metrics (Fig. S1), duplicates and related samples up to the 3rd degree (one from each pair), and samples with ambiguous/discordant sequencing gender (Fig. S2) were removed. Using multidimensional scaling (5 principal components; see Additional file 1), the genotypes of the remaining samples were projected on the 1000 Genomes (12) space (Fig. S3). The major continental ancestry was then predicted using a Support Vector Machine (see Additional file 1). Samples labeled as European were further matched using the top-two principal components (PCs) from multidimensional scaling projection (total of 10 PCs) on the 1000 Genomes European samples space (Fig. S3). Following this baseline filtering, 7,836 cases and 8,822 controls (out of 13,197 cases and 13,299 controls) remained for subsequent analysis. These filtering steps were performed using the Genome Analysis Toolkit (GATK) v4.1.4.1 (13), PLINK v1.9 (14) and KING v2.2.4 (15). For additional details, see the supplemental methods for *Baseline sample quality control* (Additional file 1).

### Baseline variant QC

Variants located outside Gencode v.33 (16) coding sequencies (CDS) boundaries or in low complexity regions (17) were not considered. Multi-allelic calls were split, and the variants were normalized. Variants with low variant quality score log-odds, covered at mean depth < 10x in the baseline filtered sample set, at minimum depth of 10x in less than 95% of cases and controls, with large difference in mean depth or call rate between the cases and controls, or with allele count equal to 0 were removed. Low depth/quality genotype calls were set to missing. Variant filtering was performed using bcftools/htslib v1.10.2 (18), vt v0.57721 (19) and GATK (13) as detailed in the supplemental methods for *Baseline variant quality control* (Additional file 1).

### Case control matching and call rate harmonization

We performed multiple iterations of principal component analysis (PCA) on our baseline filtered dataset. Following the first round of PCA (10 PCs) using PLINK (14), the Swedish Schizophrenia Study control cohort showed poor clustering with the rest of study samples on the top-two components and was removed. Subsequently, outliers on top-two principal components were filtered using EIGENSTRAT v6.1.4 (20,21). For details, see the supplemental methods (Additional file 1). To handle the residual stratification caused by platform differences, we removed all variants with call rate < 95% in any of the remaining sequencing cohorts (Epi25, Leicester, Ottawa, ATVB) or with difference in call rate > 0.5% between any pair of cohorts. Also, variants with Hardy-Weinberg Equilibrium *p* value less than 1×10^-6^ (on the combined case-control cohort) were excluded. These variant calling metrics were obtained using bcftools/htslib (18) and PLINK (14). The final analysis set included 7,589 cases (DEE=1,003, GGE=3,064, NAFE=3,522) and 3,962 matched controls (ATVB = 1,673, Leicester=1,082, Ottawa=924, Epi25 Italian=283) of non-Finnish European ancestry (Table S1 and Table S2). We observed a total of 1,267,392 variants in the final dataset, comprising 1,247,342 SNVs and 20,050 indels. The quality-controlled case-control cohort (Fig. S5) showed a balanced distribution of variants and comparable variant calling metrics (Fig. S6). The use of predominantly male or male-only control cohorts from ATVB and Leicester studies resulted in a misbalanced sample sex ratio (53.6% female cases vs. 19.4% female controls). The effect of this imbalance was addressed in a secondary analysis as will be detailed.

### Qualifying Variants

The variants were annotated using snpEff v4.3 (22) and Annovar v20191024 (23). We focused on URVs as these have shown a strong burden of deleterious pathogenic variants in multiple studies of epilepsy and other neurological disorders (8,10,24–28). Here, URVs were defined based on their population Minor Allele Frequencies (MAFs) in DiscovEHR (MAF_DiscovEHR_) and gnomAD r2.1 (MAF_gnomAD_) population database (29,30) and their Minor Allele Counts (MACs) calculated separately for each analysis (MAC_DEE+Controls_, MAC_GGE+Controls_, MAC_NAFE+Controls_), as follows: (i) MAF_DiscovEHR_ = 0 (ii) MAF_gnomAD_ ≤ 2×10^-5^ (iii) MAC_Epilepsy+Controls_ ≤ 3 in the respective analysis group. Performing three separate analyses for the three epilepsy phenotypes, with independent calculation of MAC in each analysis, was intended to provide stringent control for inflation. Accordingly, the reported variant counts in the control sets may differ slightly between the three analyses. URVs were categories further into multiple conditions based on their functional consequences and collapsed by gene as qualifying variants (QVs). We considered twelve non-synonymous variant conditions including protein-truncating (presumed loss-of-function) and multiple groups of missense variants (mix of loss- and gain-of-function mechanisms) as well as a (thirteenth) synonymous control condition (presumed neutral). The grouping of missense QVs in multiple (partially overlapping) conditions focused on three perspectives: conventional in-silico deleteriousness, constraint and paralog conservation. It was based on multiple predictions, namely, PolyPhen2 (PPh2) (31), Sorting Intolerant From Tolerant (SIFT) (32), Missense Badness Polyphen and Constraint (MPC) (33), Missense Tolerance Ratio (MTR) (34), Constrained Coding Regions (CCR) (35) and para-Z-score for paralog conservation (36). PPh2 (31) and SIFT (32) are two conventional, commonly used, in-silico missense deleteriousness scores that are widely used in genetic studies to identify likely benign and likely deleterious variants based on a number of features including the sequence, phylogenetic and structural information. MPC score (33) aims to identify regions within genes that are specifically depleted of missense variation and combines this information with variant-level metrics that measures the increased deleteriousness of amino acid substitutions when they occur in missense-constrained regions. MTR score (34) estimates the intolerance of genic regions by comparing the observed proportion of missense variation to the expected proportion in the sequence context of the protein-coding region under study. While MPC and MTR scores are scaled down to individual missense alterations, CCR score (35) aims to identify coding regions that are completely devoid of variation in population databases. Functionally critical protein regions are usually encoded by bases in regions with high CCR scores. Paralog conservation-based missense variant analysis was recently shown to aid variant prioritization in neurodevelopmental disorders (36). It has been proposed that most disease genes in humans have paralogs (37). The analyzed variant conditions (Table S6) were: (i) Benign missense variants: as predicted by PPh2 and SIFT. (ii) Damaging missense variants as predicted by PPh2 and SIFT. (iii) Protein Truncating Variants (PTVs) that included stop-gained, start-lost, frameshift, splice-donor and splice-acceptor variants. (iv) All functional variants combined PTVs, in-frame indels and damaging missense variants. (v) “MPC 1” missense variants: constrained missense with MPC score ≥ 1. (vi) “MPC 2” missense variants: highly constrained missense with MPC score ≥ 2 (enriched for *de novo* variants). (vii) “MTR ClinVar” missense variants: constrained missense with MTR score ≤ 0.825 which is the median for ClinVar variants not denoted as *de novo*. (viii) “MTR De Novo” missense variants: highly constrained missense with MTR score ≤ 0.565 which is the median for ClinVar *de novo* variants. (ix) “CCR 80” missense variants: highly constrained missense variants in regions with CCR score ≥ 80, with MPC score ≥ 1, and MTR score ≤ 0.825. (x) “paralog-non-conserved”: missense variants located in sites not conserved across paralog genes as indicated by a para-Z-score ≤ 0. (xi) “paralog-conserved”: missense variants located in sites conserved across paralog genes as indicated by a para-Z-score > 0. (xii) “paralog highly conserved”: missense variants in highly conserved sites between paralog genes with para-Z-score ≥ 1. (xiii) “Synonymous” variants that served as a control condition for inflation.

### Gene-sets

92 gene-sets were tested. In addition to exome-wide burden testing (one gene-set of all protein coding genes), we defined additional 91 specific gene-sets as follows: (a) 34 sets based on gene expression patterns in the brain: brain-expressed genes grouped by their intolerance profiles including loss-of-function intolerant and missense intolerant genes (30,38), genes grouped by their regional brain expression in the cortex and hippocampus from Genotype-Tissue Expression project v8 (39), developmentally-relevant genes (40–42), brain-enriched genes from the Human Protein Atlas v20.1 (43,44), genes enriched in specific cell types (45,46) (neurons, excitatory and inhibitory neurons, glia, astrocytes, microglia, oligodendrocytes, and endothelial cells) and localization in certain neuronal compartments (47,48); (b) 28 functional groups including ion channels (8), GABA_A_ receptors (9), excitatory receptors (9), GABAergic pathway (9), PSD-95 interactors (8), Gene Ontology (GO) genesets of GABAergic and glutamatergic synapses (40,41,49), neuronal pathways from Kyoto Encyclopedia of Genes and Genomes (KEGG) (50) and neuronal gene-sets from Reactome database (51); (c) 14 gene-sets of known disease-related genes including monogenic epilepsy-causing genes (8–10), epilepsy genome-wide association study (GWAS) top-ranking genes from positional mapping within a window of 250 kb around significant loci plus mapping based on chromatin interaction between gene promoters and the significant locus (52), co-regulated genes in the brain (53,54); and (d) 15 nonneuronal gene-sets including genes not expressed in the brain (39,43,44), cancer and metabolic pathways (50). The gene-sets are outlined in Table 1, Table S7 and Table S8 (see Additional file 1 and Additional file 2).

**Table (1):**
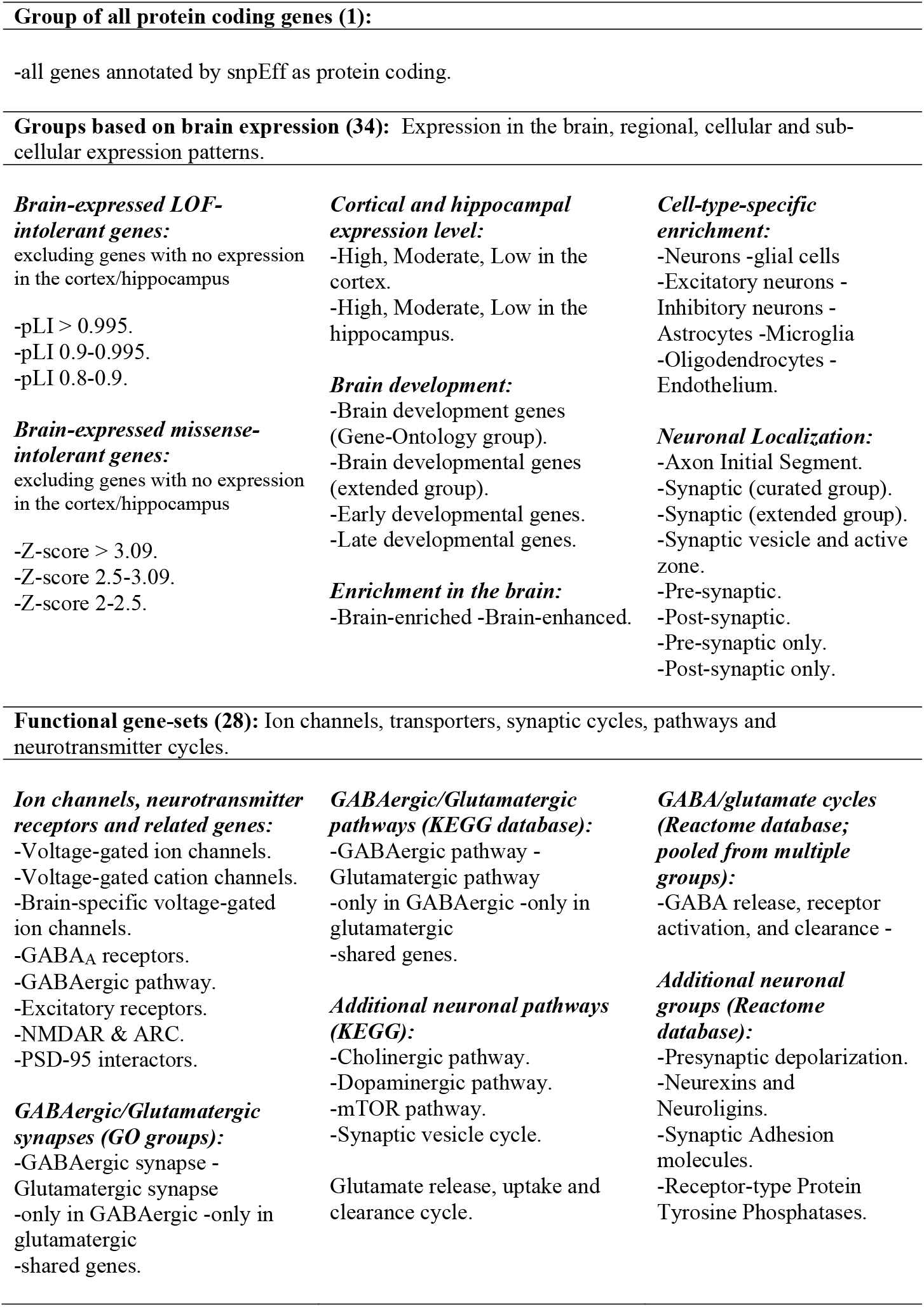
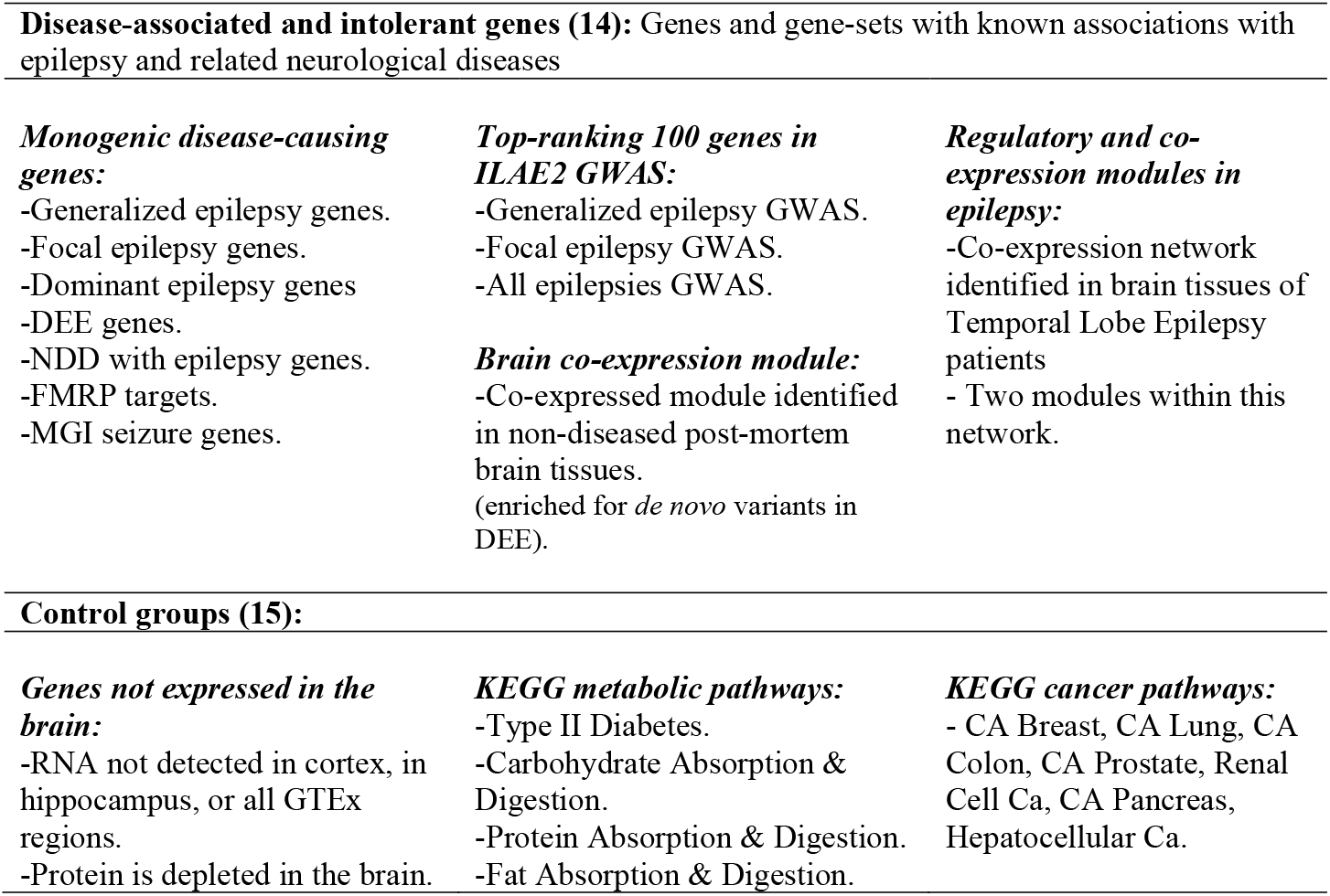
Gene-sets investigated in this study.

### Gene-set burden analysis

We examined the burden of QVs in thirteen variant conditions (Table S6) for 92 gene-sets in three epilepsy phenotypes (DEE, GGE, and NAFE) against a set of matched controls. Gene-set burden testing was done using logistic regression by regressing case-control status on the individual QVs counts. In each sample, ultra-rare variants (URVs) that matched the specific analysis condition were collapsed by gene into QVs and these QVs were aggregated (summed) across a target gene-set to get a burden score (assuming equal weights and direction of effects) which was used as a predictor in a binomial model while adjusting for additional covariates (sex, top ten principal components, exome-wide variant count, and exome wide singletons count) using *glm*() function from *stats* package (55). Likelihood ratio test (LRT) from *lmtest* package (56) was used to compare a model with QVs burden and covariates as predictors against a null model (covariates only). Log-odds from LRT and their respective 95% confidence intervals and *p* values are presented here as a measure of enrichment in tested gene-sets (Additional file 3). We employed a Benjamini-Hochberg false discovery rate (FDR) multiple testing adjustment for *p* values that accounted for 3312 tests (92 gene-sets x 3 epilepsy phenotypes x 12 non-synonymous variant conditions, excluding the control condition of synonymous variants) as implemented in *p.adjust*() function from *stats* package (55). The cut-off for substantial enrichment was defined as FDR-adjusted *p* value < 0.05. For simplicity, *p* values (FDR-adjusted except for synonymous variants) are indicated throughout the presented plots using stars as follows: no star > 0.05, * < 0.05, ** < 0.005, *** < 0.0005, **** < 0.00005. To estimate the extent of bias that might have been introduced by the imbalance in male-to-female ratios between cases and controls, we performed a secondary analysis excluding chromosome X genes (Additional file 4). Also, to ensure adequate control for any bias introduced by differences in capture kits, we performed another secondary analysis between two groups of control samples (Leicester study controls vs. Ottawa and ATVB controls) representing two main enrichment kits (Illumina ICE vs. Agilent SureSelect kits). The statistical analysis was performed in R 3.3.3 (55). The analysis approach is outlined in Fig. 1 and the methods are detailed in Additional file 1. Tables of tested gene-sets and genes in each set are provided in Additional file 2.

**Fig. 1:**
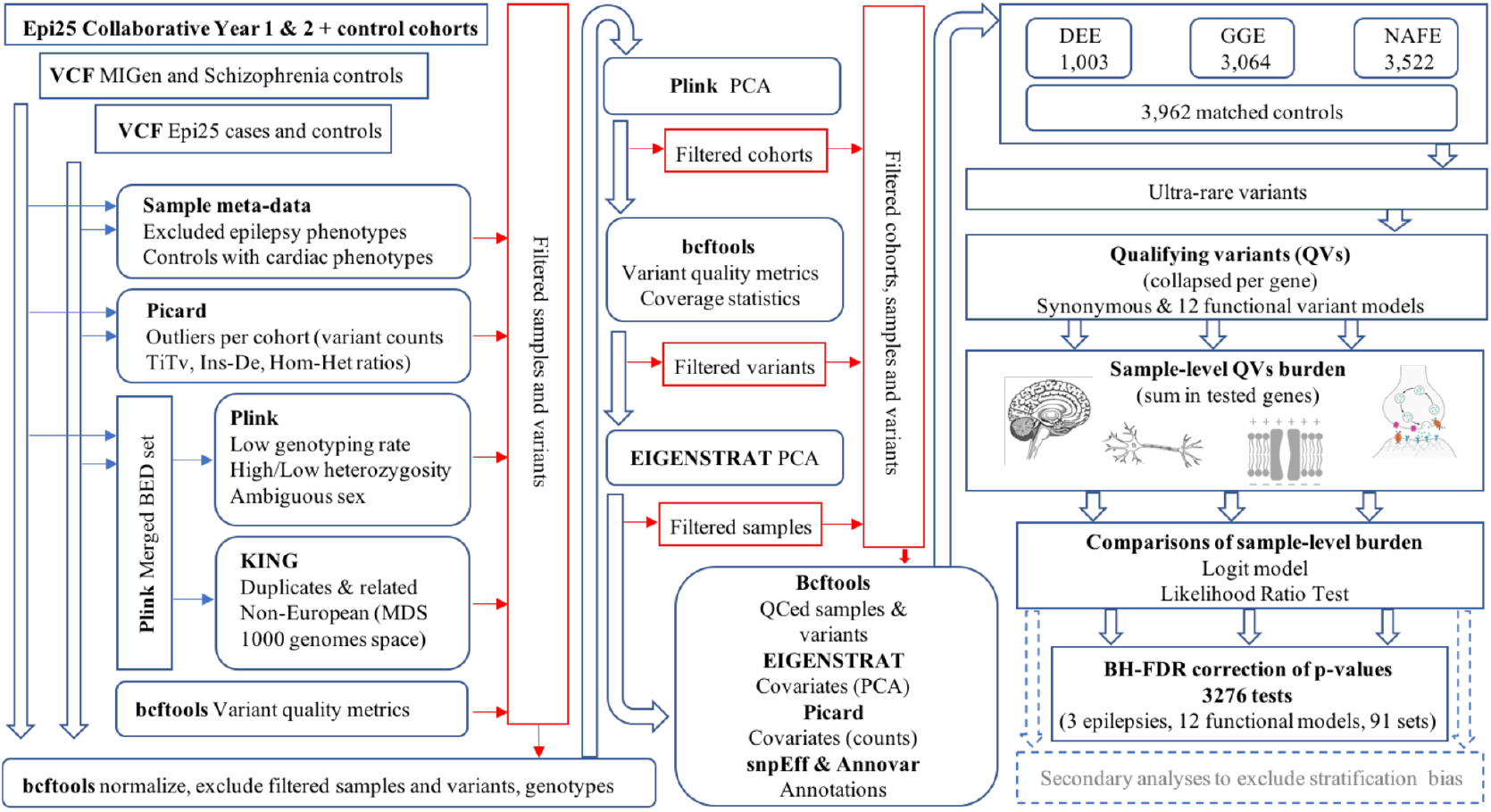
Outlines of the burden analysis method. Thirteen (twelve functional and one synonymous) variants models with focus on missense variants in constrained or paralog-conserved sites were tested in the three epilepsy phenotypes against a shared set of matched controls. The burden was examined in 92 gene-sets (detailed in table 1) using a logistic regression model with the count of qualifying variants per sample as a predictor and sample sex, ten principal components, singletons and exome-wide variant counts as covariates. Two secondary analyses were performed: one analysis restricting gene-sets to autosomal genes (to exclude bias introduced by male-to-female ratio imbalances) and another analysis testing controls prepared for exome sequencing using Illumina ICE capture kits against controls prepared with Agilent SureSelect capture kits (to exclude bias caused by differences in enrichment kits).

## Results

### URVs excess in brain-expressed genes

First, we investigated the burden of URVs across all protein coding genes following the analysis approach outlined in Fig. 1. This revealed a clear enrichment in constrained missense variants that was maximum in consensus constrained coding regions predicted by Missense-badness Polyphen and Constraint (MPC), Missense Tolerance Ratio (MTR) and Consensus Coding Regions (CCR) scores (Fig. 2). The combination of the three metrics identifies highly deleterious variants in functionally critical genic regions (see methods). In this particular analysis in all three phenotypes, about half of the cases, in contrast to roughly one-fourth of controls, harbored one or more QVs in highly constrained regions (Fig. S10). A previous similar analysis of this dataset (10) examined loss-of-function intolerant genes and demonstrated an increased burden in ultra-rare constrained as well as protein truncating variants (PTVs). Here, the examination of brain-expressed intolerant genes showed, similarly, a marked enrichment in PTVs in addition to a burden in constrained missense variants that is comparable to what is seen exome-wide (Fig. 2 and Fig. S11).

**Fig. 2:**
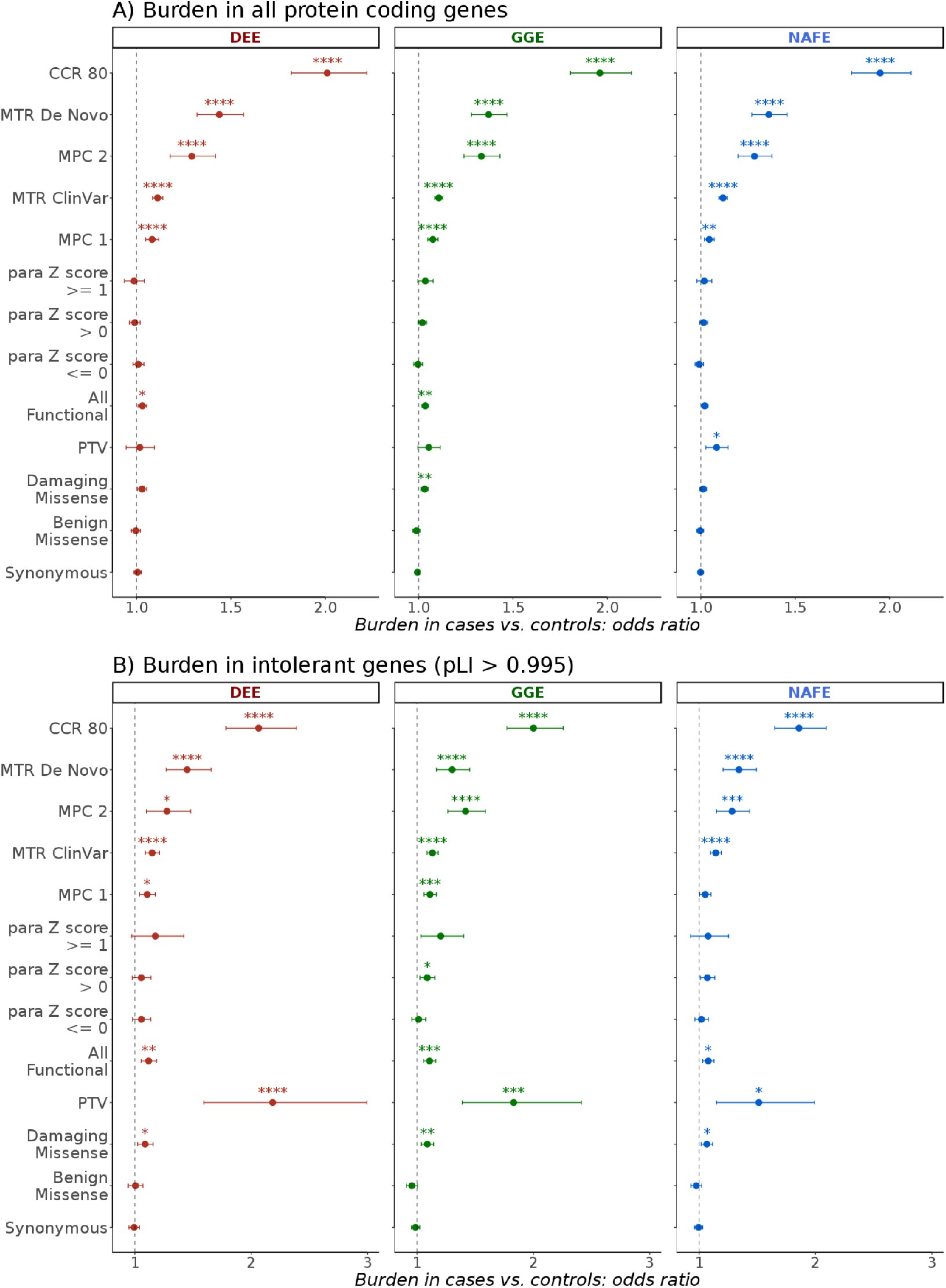
Exome-wide burden of ultra-rare variants in the epilepsies. The burden in developmental and epileptic encephalopathies (DEE), genetic generalized epilepsies (GGE) and non-acquired focal epilepsies (NAFE) in 19,402 protein coding genes **(A)** and 1,743 loss-of-function intolerant genes with pLI score > 0.995 **(B)** is shown in multiple test and control conditions (y-axis) as odds ratio (x-axis) from Likelihood Ratio Test (bars indicate 95% confidence intervals). FDR corrected *p* values (not corrected for synonymous variants) are indicated with stars as follows: no star > 0.05, * < 0.05, ** < 0.005, *** < 0.005, **** < 0.0005. There is an incrementing burden with a higher level of missense constraint. Both synonymous and benign missense alterations are not enriched, suggesting sufficient control for inflation.

When we examined protein coding genes grouped by their relative brain expression, damaging missense variants were only substantially enriched in genes highly expressed in the cortex or hippocampus, whereas those expressed at medium or low levels only showed an enrichment for the most constrained missense variants (Fig. 3). Genes with depleted expression in the brain did not show a substantial enrichment for any variant type (Fig. S15). Genes showing a higher expression in the adult brain compared to other tissues (brain-enriched & brain-enhanced) were also preferentially enriched, as well as genes associated with brain development. Genes related to late rather than early development showed a slightly higher enrichment in all three phenotypic groups (Fig. 3).

**Fig. 3:**
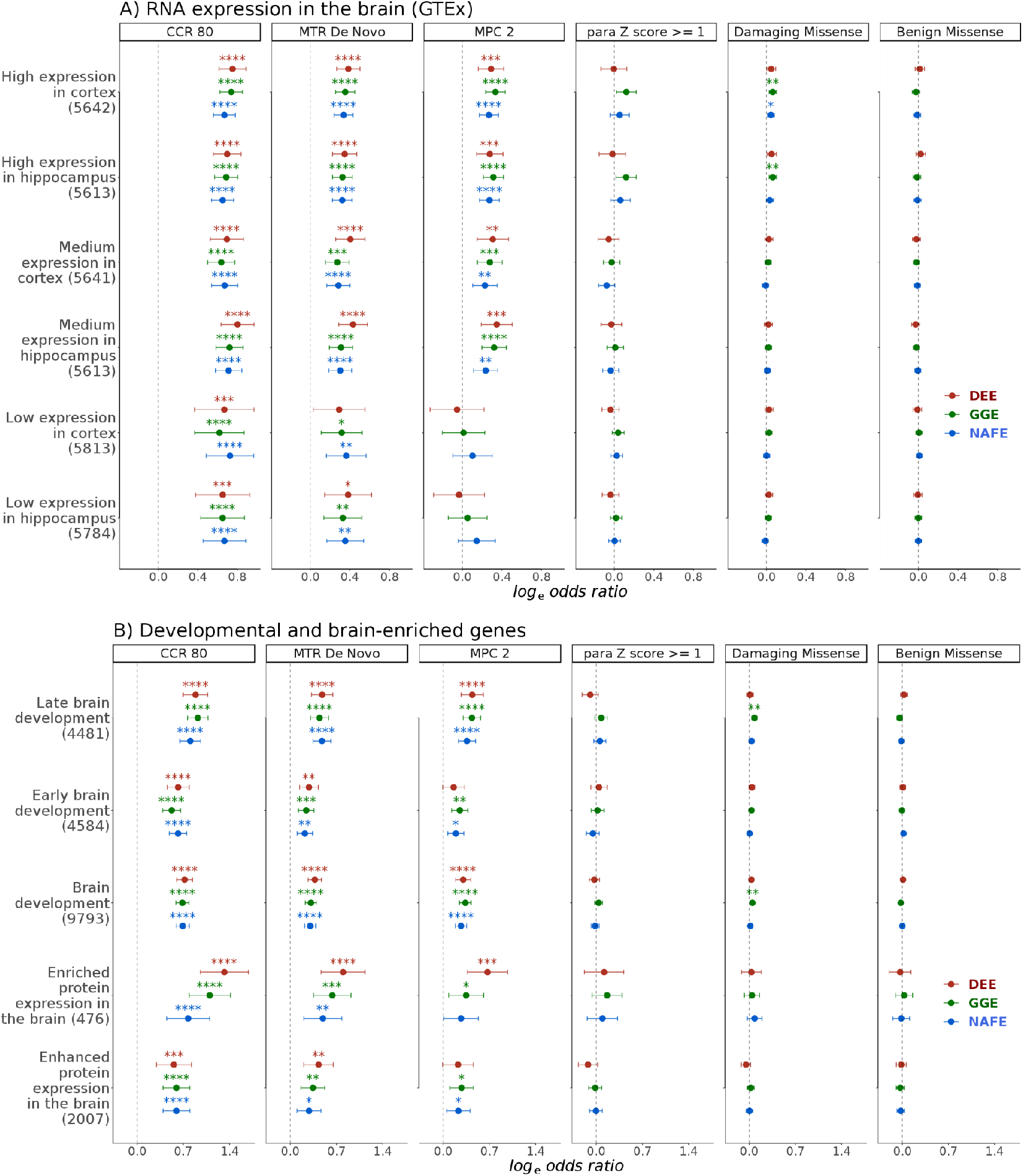
Burden of ultra-rare missense variants in brain expressed and developmental genes. The burden of benign or damaging missense variants and missense variants in highly paralog-conserved or highly constrained sites in developmental and epileptic encephalopathies (DEE), genetic generalized epilepsies (GGE) and non-acquired focal epilepsies (NAFE) is shown in gene-sets based on levels of RNA/protein expression in the cortex and hippocampus **(A)** or enrichment in adult or developing brain **(B)**. Gene-sets are shown on the y-axis (number of genes between brackets). Log odds ratio (Likelihood Ratio Test) are shown on the x-axis (error bars indicate 95% confidence intervals). The variant conditions are shown in vertical panels. FDR corrected *p* values (not corrected for synonymous variants) are indicated with stars as follows: no star > 0.05, * < 0.05, ** < 0.005, *** < 0.005, **** < 0.0005. Brain enriched genes (Human Protein Atlas: more than four-fold expression compared to other tissues) show higher burden in DEE compared to those genes with only enhanced expression (higher but less than four-fold expression in the brain). This difference is less prominent in GGE and NAFE. Developmental genes enriched in late development show higher burden than genes enriched in early development.

Focusing further on cell-type specific expression, neuron-specific genes were preferentially affected compared to those enriched in glial cells, particularly in GGE (Fig. 4). To obtain further insight into the nature of this neuronal enrichment, we used sets of genes representing paralogs of mouse genes found to be enriched in excitatory or inhibitory neurons (see Additional file 1). Interestingly, genes preferentially expressed in inhibitory neurons showed an increased burden only in GGE, whereas those preferentially expressed in excitatory neurons showed a more prominent signal in NAFE. Since well-established epilepsy genes, like ion channels and receptors, show differential distributions in different neuronal compartments (57,58), we examined further sets of genes based on subcellular localization. We found that pre- and postsynaptic genes were enriched with variants in cases vs. controls, as well as a very small set of 17 genes located in axon initial segments (most prominent in DEE) (Fig. S13).

**Fig. 4:**
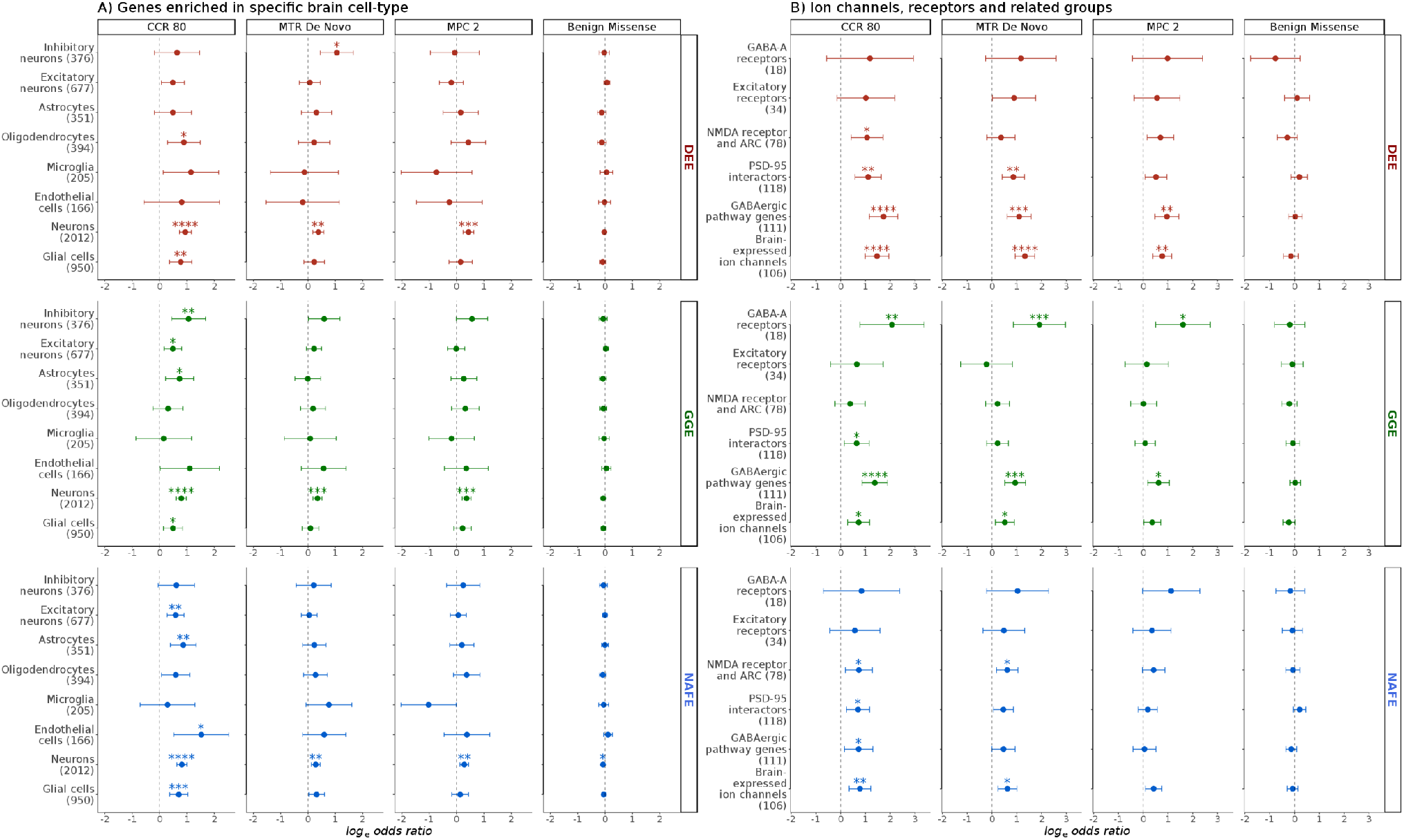
Burden in neuronal and glial cells, ion channels, receptors and related interactors. The burden in developmental and epileptic encephalopathies (DEE), genetic generalized epilepsies (GGE) and non-acquired focal epilepsies (NAFE) in shown on the x-axis (log-odds from Likelihood Ratio Test; error bars indicate 95% confidence intervals). Gene-sets are shown on the y-axis (number of genes between brackets). The variant conditions are shown in vertical panels. FDR corrected *p* values (not corrected for synonymous variants) are indicated with stars as follows: no star > 0.05, * < 0.05, ** < 0.005, *** < 0.005, **** < 0.0005. **(A)** Comparisons based on the cellular expression patterns show that the enrichment is prominent in neuron-enriched compared to glial genes, particularly in GGE. GGE shows relatively higher enrichment in inhibitory neuron genes while NAFE shows a relatively higher burden in excitatory neuronal genes. **(B)** Variants in GABA_A_ receptors and GABAergic pathway are preferentially enriched in GGE compared to groups of genes coding for NMDA receptor and neuronal activity-regulated cytoskeleton-associated protein (ARC). In NAFE, the variants are enriched in the NMDA receptor & ARC gene-sets, but not GABA_A_ receptors.

### Burden of URVs in ion channel, neurotransmitter receptor encoding and related genes

Next, we examined functional gene-sets that could, more specifically, underlie the observed enrichment in neuronal and synaptic genes. Ion channels, neurotransmitter receptors and transporters are widely implicated in epilepsy, especially in monogenic and familial forms, displaying considerable phenotypic heterogeneity and presenting as mild or severe epilepsies (59–61). Variants in GABA_A_ receptors were enriched in GGE but not in DEE or NAFE while those in gene-sets representing genes encoding N-Methyl-D-Aspartate receptor and Activity-Regulated Cytoskeleton protein (NMDAR-ARC) were enriched in NAFE and DEE. A comprehensive gene-set for the GABAergic pathway genes (9) showed a prominent signal in GGE and DEE, and less in NAFE. In contrast, a gene-set representing PSD-95 interactors showed comparable enrichment in NAFE and GGE (Fig. 4). Brain-expressed ion channels were found to be enriched for highly constrained missense variants (CCR 80 condition) in common as well as rare epilepsies (Fig. 4).

### Patterns of burden in gene-sets representing inhibitory vs. excitatory signaling

We then compared the patterns of URVs burden in genes involved in the GABAergic (main inhibitory) pathway and synapse against those in the glutamatergic (main excitatory) pathway and synapse in the brain, by examining their unique and overlapping genes based on KEGG pathways (50) or GO synaptic gene-sets (41) and sets of specific receptors (Fig. 5). GGE showed a higher burden in GABAergic vs. glutamatergic synapse (GO) and pathway (KEGG) genes, in GABA_A_ receptors vs. excitatory receptors/NMDAR-ARC genes, and in GABAergic pathway genes (comprehensive gene-set) vs. PSD-95 interactors, thus matching the higher burden in genes representing inhibitory vs. excitatory neuronal signaling. The CCR 80 analysis of GO gene-sets in NAFE showed a higher burden in glutamatergic vs. GABAergic synapse genes, akin to the pattern seen in genes enriched in excitatory vs. inhibitory neurons. The analysis of KEGG glutamatergic vs. GABAergic pathway genes did not confirm this finding (Fig. 5). It is notable that the overlap between GO synapse and KEGG pathway gene-sets is minimal (Fig. S18), and the size of GO and KEGG gene-sets was comparable in GABAergic but discordant in glutamatergic genes.

**Fig. 5:**
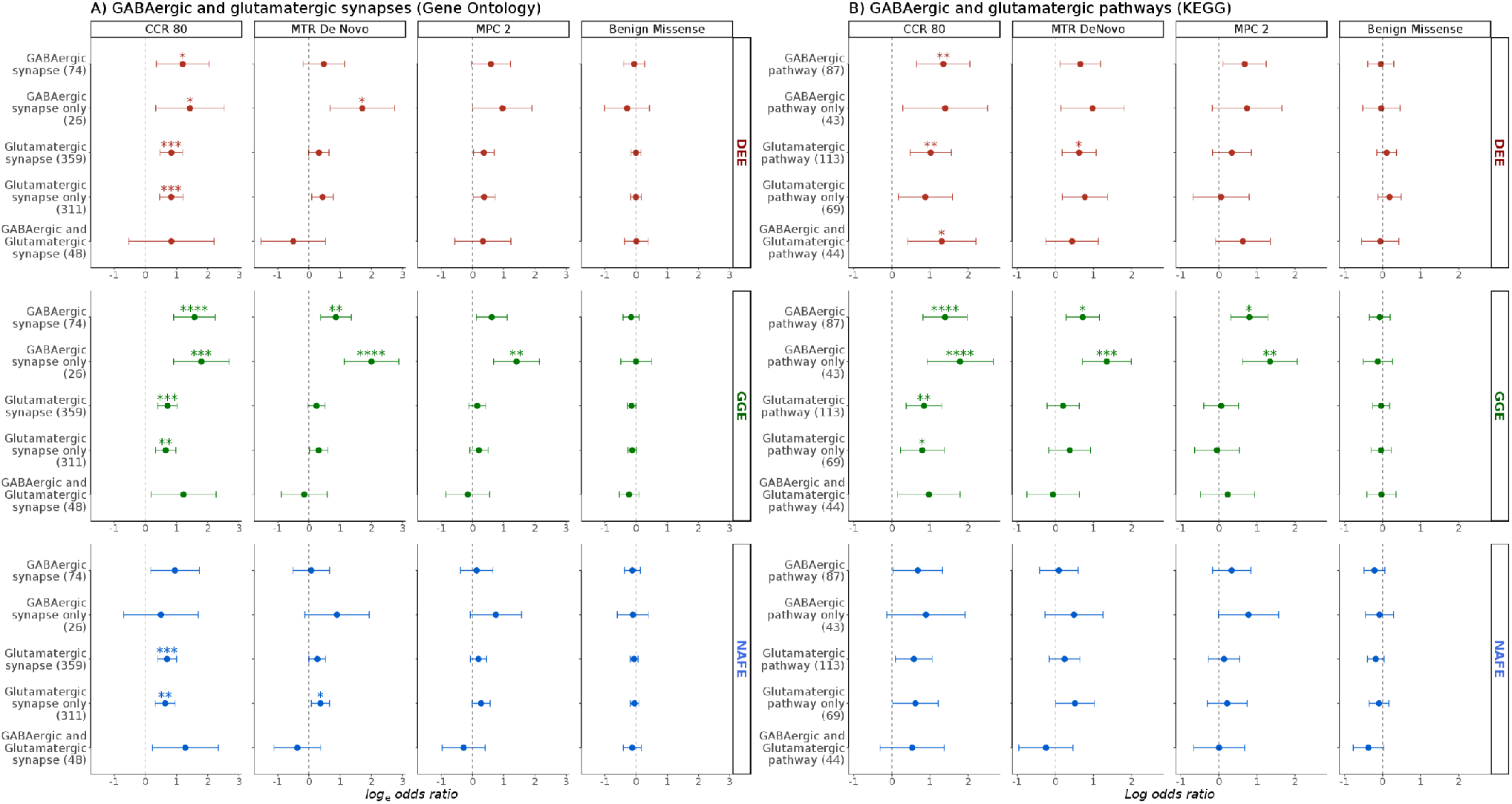
Enrichment in major neuronal synapses and pathways. Comparison of enrichment patterns in GABAergic and glutamatergic synapses and pathway genes based on Gene-Ontology (GO) **(A)** or Kyoto Encyclopedia for Genes and Genomes (KEGG) **(B)** The burden in developmental and epileptic encephalopathies (DEE), genetic generalized epilepsies (GGE) and non-acquired focal epilepsies (NAFE) in shown on the x-axis (log-odds from Likelihood Ratio Test; error bars indicate 95% confidence intervals). Gene-sets are shown on the y-axis (number of genes between brackets). The variant conditions are shown in vertical panels. FDR corrected *p* values (not corrected for synonymous variants) are indicated with stars as follows: no star > 0.05, * < 0.05, ** < 0.005, *** < 0.005, **** < 0.0005. Complete groups, genes specific to one of the two synapses/pathways as well as their intersection were tested. Both GO and KEGG gene sets show an increased burden in GGE when “GABAergic only” genes are tested. GGE shows a relatively higher burden in GABAergic compared to glutamatergic gene sets.

Altogether, these comparisons of the burden in missense variants in highly constrained sites between GGE and NAFE (Fig. 4 and Fig. 5) suggest the following patterns: (i) brain-expressed ion channels, genes enriched in excitatory neurons, enriched in astrocytes, PSD-95 interactors, GABAergic and glutamatergic synapse/pathway genes show an increased burden in cases vs. controls both in GGE & NAFE; (ii) in GGE, this enrichment is coupled with a stronger enrichment in inhibitory neuronal genes, in GABA_A_ receptors and in GABAergic synapse-specific genes (higher burden in inhibitory vs. excitatory gene-sets). (iii) in NAFE, this is accompanied by an absence of enrichment in the later genesets and increased burden in the NMDAR-ARC gene-set (higher burden in excitatory vs. inhibitory gene-sets).

### Burden in gene-sets of known epilepsy-related genes

The previous Epi25 Collaborative analysis (10) demonstrated a high burden of missense variants in constrained sites (with MPC score ≥ 2) in DEE, GGE, and NAFE, seen in dominant epilepsy genes, DEE genes, and NDD-Epilepsy genes. We observed similar enrichment patterns (Fig. 6) in MPC 2 and MTR De Novo conditions (enriched for de *novo* mutations). In addition, we saw a substantial enrichment for other variant conditions (MPC score ≥ 1 and MTR ClinVar) with lower odds ratios. Limiting the analysis to highly constrained genic regions (CCR 80 condition) resulted in a marked increase in URVs burden, as was the trend in all the tested gene-sets so far. Testing these sets also unraveled strong enrichment in PTVs and missense variants in paralog-conserved sites. PTVs and missense variants in paralog-conserved sites did not show substantial enrichment in exome-wide analysis and most of other expression-based, localization-based or pathway-based gene-sets. However, we saw a modest increase in PTV burden in highly intolerant genes with probability of Loss-of-function Intolerance (pLI) > 0.995 in all epilepsies (Fig. 2 and Fig. S11). In a gene-set of known DEE genes, where highly intolerant genes are rather prevalent, we saw a prominent enrichment in PTVs burden in DEE. Also, there was an increased burden in missense variants in paralog-conserved sites in sets of epilepsy-related disease genes (DEE genes, dominant Epilepsy genes, NDD-Epilepsy genes). This burden was very strong in DEE but not as remarkable in GGE and NAFE (Fig. 6).

**Fig. 6:**
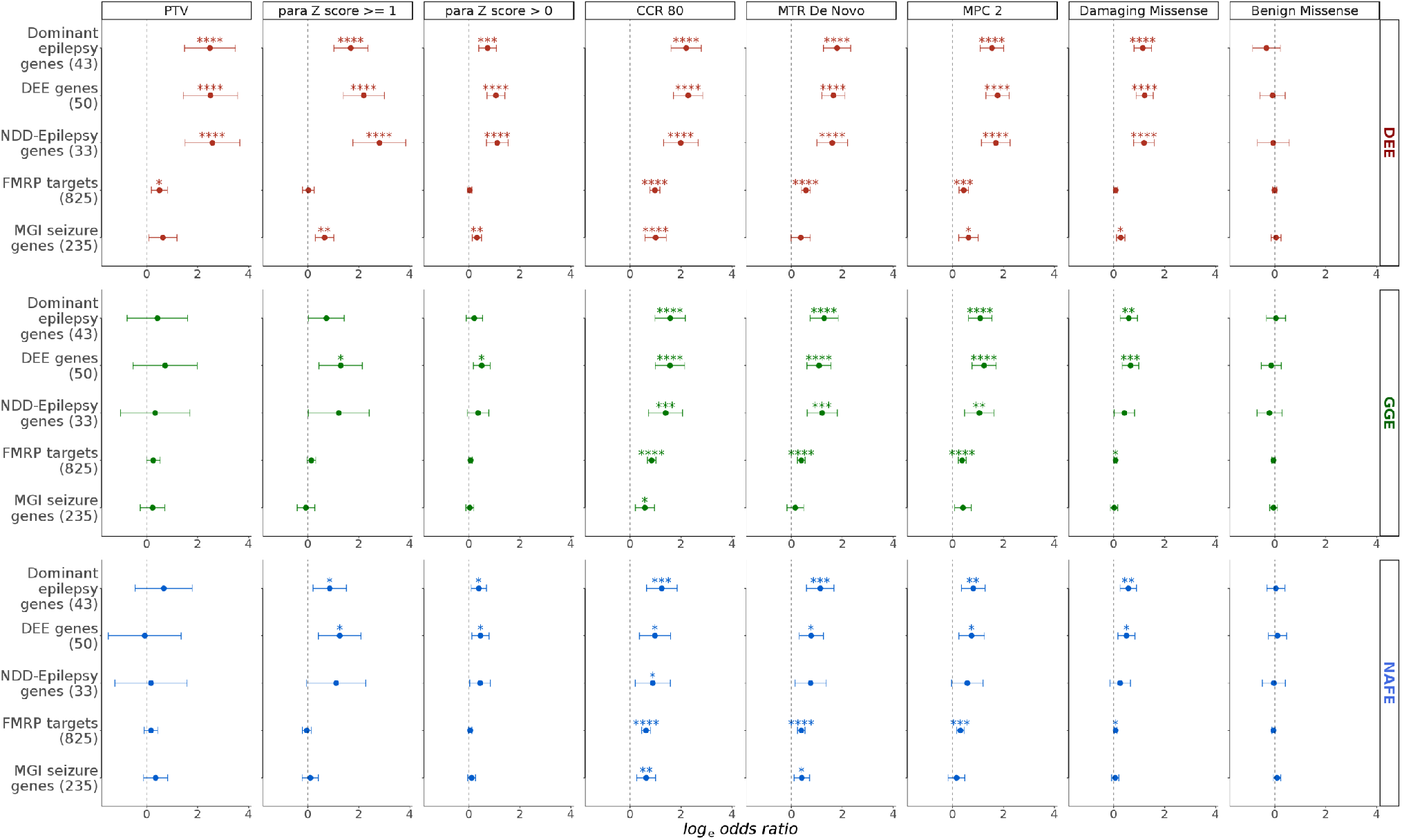
Burden of ultra-rare variants in groups of epilepsy-related known disease genes. The burden in five gene-sets (y-axis; number of genes between brackets) in developmental and epileptic encephalopathies (DEE), genetic generalized epilepsies (GGE) and non-acquired focal epilepsies (NAFE) (horizontal panel) in selected variant conditions (vertical panels) is shown on the x-axis (log odd ratios from Likelihood Ratio Test; error bars indicate 95% confidence intervals). FDR corrected *p* values (not corrected for synonymous variants) are indicated with stars as follows: no star > 0.05, * < 0.05, ** < 0.005, *** < 0.005, **** < 0.0005. NDD-Epilepsy: neurodevelopmental disorders with epilepsy. FMPR: Fragile-X Mental Retardation Protein targets. MGI: Mouse Genome Informatics database.

### Enrichment in top GWAS hits captures divergence between common epilepsies

Recent efforts from the ILAE consortium on complex epilepsies identified multiple associations in a large GWAS of common epilepsies (52). To examine the hypothesis that genes located near the top GWAS hits are also affected by rare variants, we tested the enrichment in sets of the 100 top-ranking genes derived from the ILAE GWAS in generalized, focal, and all epilepsies. Interestingly, when limiting the analysis to Consensus Coding Regions (CCR80 condition), top-ranking genes derived from the GWAS of either generalized or focal epilepsies were preferentially enriched for rare variants in the respective phenotypic groups of GGE and NAFE (Fig. 7). Although the observed enrichment was rather subtle, this result was corroborated by a similar pattern for two, rather small, sets of known epilepsy genes that are predominantly associated with either generalized or focal epilepsy (9).

**Fig. 7:**
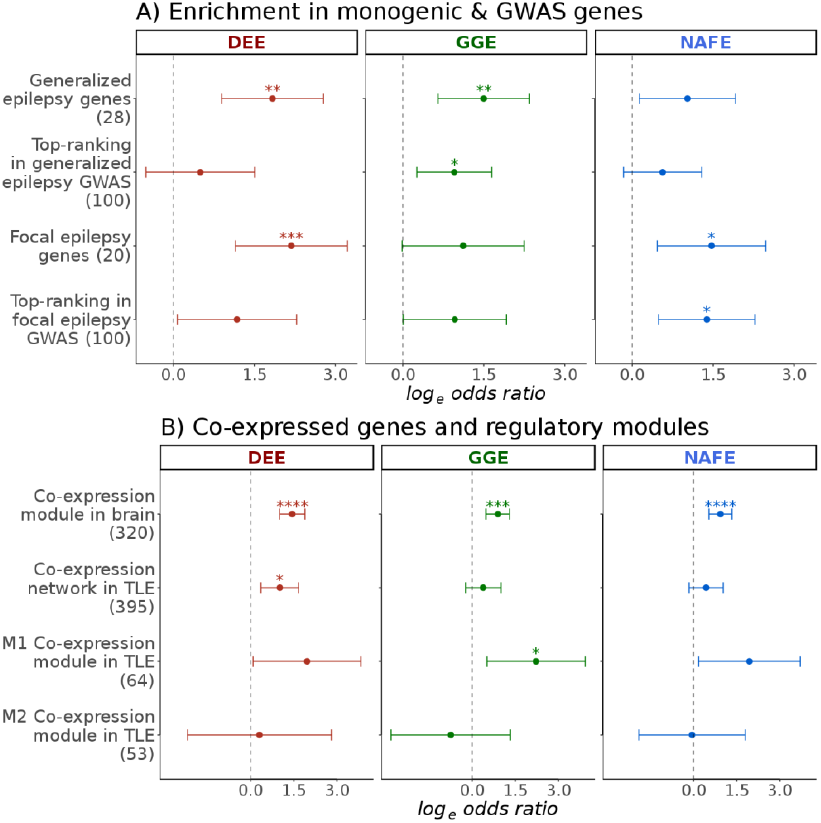
Risk elements in GWAS top-ranking genes and co-expression modules. The burden of missense variants in highly constrained sites (log-odds on the x-axis; error bars indicate 95% confidence intervals) in developmental and epileptic encephalopathies (DEE), genetic generalized epilepsies (GGE) and non-acquired focal epilepsies (NAFE) is shown in monogenic epilepsy genes, top-ranking epilepsy GWAS genes, brain co-expression modules and TLE-related co-expression modules (y-axis; number of genes between brackets). FDR corrected p values (not corrected for synonymous variants) are indicated with stars as follows: no star > 0.05, * < 0.05, ** < 0.005, *** < 0.005, **** < 0.0005. (A) Generalized or focal epilepsies (presumed monogenic) as well as top-ranking 100 genes from GWAS of generalized and focal epilepsies are preferentially enriched for constrained missense variants (CCR 80) in respective phenotypic cohorts indicating a possible convergence between common and rare variants in GWAS genes. (B) Enrichment in co-expressed genes identified in postmortem brain tissues of healthy individuals (module of 320 genes) or in brain tissues from TLE patients (network of 395 genes) as well as two sub-modules of this network (M1 and M2).

### Brain- and epilepsy-related co-expression modules

We also aimed to touch upon the role of brain co-expression modules identified in post-mortem brain tissues from healthy individuals (54) and contrast these to the networks and modules identified in brain tissue derived from epilepsy patients (53). A brain expression module was found to be substantially enriched for rare deleterious variants in an independent cohort of DEE (54). A link to common epilepsy phenotypes was also inferred, but a burden in ultra-rare variants was not examined so far. This module showed a non-specific enrichment in all three epilepsy subtypes with highest odds in DEE. It is noteworthy that this module overlaps largely with known epilepsy genes (Fig. S19). In resected hippocampi of individuals with temporal lobe epilepsy (TLE), Johnson and colleagues identified two co-expression modules within a gene-regulatory transcriptional network (53). A subtle enrichment was seen in these modules in DEE and GGE, but not NAFE (Fig. 7).

### Additional neuronal and non-neuronal pathways

Other neuronal gene-sets were enriched in our analysis (Fig. S14). Genes encoding neurexins and neuroligins, important elements of pre- and post-synaptic interaction promoting adhesion between dendrites and axons (62), were enriched in DEE (Fig. S14). Also, the synaptic vesicle cycle pathway (KEGG) showed a prominent signal in both DEE and GGE. We also examined the burden in the mTOR pathway (KEGG), hypothesizing that it could have potential relevance to focal epilepsies, but did not detect a substantial enrichment (Fig. S14). Interestingly, NAFE analysis displayed a burden in endothelial and astrocyte-specific genes in highly constrained genic regions (Fig. 4). Detailed results from all tested conditions including the counts of genes with observed QVs, variant counts in cases and controls, logistic regression odds of the individual QVs burden in cases vs. controls and related *p* values are provided in Additional file 3.

### Specificity of the observed enrichment patterns

Four sets of genes not expressed in the brain that were tested (high confidence genes with depleted RNA and protein expression in the brain, genes with no RNA detected in the cortex, the hippocampus or any brain tissue) were not substantially enriched in most the tested variant conditions (Fig. S15). Also, we examined eleven metabolic and cancer pathways (KEGG) to have additional insights into the specificity of the observed signals to neuronal processes and genes. Among 540 tests targeting functional variants in these non-neuronal gene-sets (3 epilepsy subtypes, 15 sets representing genes not expressed in the brain, KEGG metabolic and cancer pathways, 12 conditions/types of variants), 18 tests (3.3%) had an FDR-adjusted *p* values < 0.05. At least for some of those, the enrichment could be explained by an overlap with genes known to play a role in epilepsy. For instance, genes forming the Type II Diabetes KEGG pathway are substantially enriched in DEE (FDR-adjusted *p* values of 0.007 for MTR DeNovo and 0.01 for CCR 80 condition). This pathway contains two genes that are known to cause DEE, namely, *CACNA1A* (63) and *CACNA1E* (64). The enrichment is no longer prominent (*p* values > 0.05) after the removal of these two genes (Fig. S20).

### Bias and inflation in gene-set burden testing

The analysis for synonymous variants did not show more substantial enrichment than expected by chance, indicating sufficient control for inflation, particularly in exome-wide models and gene-sets with large number of genes. In this control condition, few tests showed *p* values < 0.05 (15 out of 276 tests of 92 gene-sets and 3 phenotypes: 5.4%). The analysis for benign missense variants, another condition that is not expected to show an increased burden in cases vs. controls (10), did not show substantial enrichment as well. Nine out of 276 tests for benign missense variants (3.2%) showed *p* values < 0.05 (only 2 with FDR-adjusted *p* values < 0.05). Possible alternative explanations for such subtle signals include residual population stratification, differences in exome capture not adjusted by covariates and the presence of synonymous variants with functional consequences (65). However, these proportions are close to the limit expected by chance under a true null hypothesis (5% with α = 0.05). A potential source of bias in our burden testing was the imbalance in male-to-female ratios between cases and controls (Table S4). We provide results from a secondary analysis that excluded all genes located on chromosome X, which shows that any bias not captured by the inclusion of sample sex as a covariate is likely marginal (Additional file 5). To exclude any major residual stratification resulting from the use of different enrichment kits, we additionally performed a controls-only analysis in which we compared control samples enriched with Illumina ICE capture kits (from Leicester study) to controls enriched using Agilent SureSelect kits (ATVB study and Ottawa study). This analysis reflected a good control for any potential bias introduced by different exome capture systems and also demonstrated that the mixing of controls included (Leicester and Ottawa) or not included (ATVB) in gnomAD is unlikely to have affected our main outcomes (Additional file 5).

## Discussion

By analyzing the sequencing data of 11,551 unrelated European individuals, we show an exome-wide burden in ultra-rare missense variants in epilepsy cases compared to controls. These variants are mainly found in constrained sites across three different subtypes of common generalized and focal epilepsies as well as rare and mostly severe developmental and epileptic encephalopathies. This ultra-rare variant burden was prominent for missense variants enriched for *de novo* alterations, yet also substantial in analyses examining a likely combination of inherited and *de novo* variants. While this burden was consistently higher with increased constraint in various gene-sets, PTVs were specifically relevant in known disease genes (Fig. 6) and brain-expressed loss-of-function intolerant genes in comparison to all protein coding genes (Fig. 2). This replicates the findings of the previous Epi25 analysis (10) which indicated an increased PTV burden in known disease-causing epilepsy related genes and genes with pLI > 0.995 but not those with pLI 0.9-0.995 (Fig. S11). In general, PTVs are known to be associated with several phenotypes with their effect being most prominent in LOF-intolerant genes (30,38,66). Similar to PTVs, we did not observe a substantial increase in the burden of missense variants in paralog-conserved sites in a group of all protein coding genes, but it was remarkably high in gene-sets of known disease genes. The increased burden was very prominent in DEE but not in GGE and NAFE. Missense variants in paralog-conserved sites showed prominent enrichment in neurodevelopmental disorders (36), a phenotypic category that overlaps largely with DEE. Although this may reflect a true disparity based on the importance of these highly conserved sites (with their disturbance possibly resulting in severe rather than mild epilepsy phenotypes), it is possible that our analysis lacked power to detect small effect sizes in common epilepsies.

In addition to their utility in estimating the contribution of certain variant conditions/types in different epilepsy phenotypes, gene-sets with known relation to epilepsy, especially to monogenic forms, constitute high-effect-size gene-sets that can also serve as technical validation sets. The presented results are consistent with previous analyses of missense variants in a small number of gene-sets examined in similar cohorts (8–10). The systematic analysis of gene-sets and more different conditions/types of variants revealed interesting findings about the neurobiology of distinct types of epilepsy and clearly revealed that the pathological burden is markedly higher in constrained coding sites and regions. Although associated with higher odds ratios of an epilepsy phenotype, these variants are not deterministic on their own, since about one-fourth of the controls also carry qualifying variants in the CCR 80 analysis (Fig. S10). As such, the phenotype is determined by a constellation of other factors, possibly including patterns of multiple variations, oligogenic contribution from rare variants (67), and polygenic risk from common variants (68).

The excess of ultra-rare missense variants in the epilepsies clearly stemmed from genes with enriched expression in the brain compared to other tissues, including developmentally relevant genes. A strong link exists between brain development and rare DEE and NDD, since both present with marked developmental deficits accompanied with different seizure types (69,70). The relation of developmental genes to common epilepsies is not as clear. At the cellular level, the enrichment was primarily seen in neuron-specific genes. Synaptic genes with their various pre- and postsynaptic localizations were main drivers in all epilepsies, with prominent signals in postsynaptic genes (Fig. S13). A similar enrichment in synaptic genes has been observed in neurodevelopmental disorders with epilepsy (70,71), schizophrenia (24) and autism (72), highlighting a shared genetic architecture not only between epilepsy subtypes but also with other related neurological disorders, as has been shown previously for common variants (73). Ion channel genes were enriched for ultra-rare, constrained variants in all epilepsy phenotypes (Fig. 4), as has been shown in previous work focusing on ultra-rare variants (8,10).

Despite the common genetic and phenotypic features, DEE, GGE and NAFE represent well-recognized phenotypic clusters with defined electro-encephalographic and clinical characteristics. Given the phenotypic severity of DEE, the prevalence of *de novo* variants and monogenic cases in DEE (those with pathogenic and likely pathogenic variants in known monogenic genes), and the description of phenotypic spectra for genes involved in DEE that also span the milder GGE or NAFE, the distinction between severe and mild epilepsies could be attributed, at least to some extent, to the severity of the genetic defects, their functional effects or their localization within certain channel regions (61,74–77). The distinction between GGE and NAFE, however, is probably functional, at least in part, as suggested by previous work demonstrating the centrality of GABAergic genes in generalized epilepsies (9,10). Also, it is well recognized that few genes present with focal, but not generalized, epilepsy syndromes (78). Here, phenotype-specific patterns were seen in comparisons of GGE and NAFE. GABA_A_ receptor genes, GABAergic pathway genes and genes enriched in inhibitory neurons were preferentially affected in GGE in comparison to the glutamatergic pathway genes. In contrast, NMDA receptor and ARC genes, and genes expressed in excitatory neurons were enriched in NAFE. Corroborating this finding for GGE, we found that the enrichment in the GABAergic pathway is stronger in core genes that are not shared with the glutamatergic pathway. An opposing pattern (albeit, not as prominent) was seen in NAFE, where the burden in genes enriched in excitatory neurons and glutamatergic synapse was more prominent than the burden in their inhibitory and GABAergic counterparts, providing evidence for the importance of the former in focal epilepsies.

Additional disparities in key gene-sets point to a possible genetic-functional divergence, so that a common background of shared risk seems to be overlaid by specific risk entities. Groups of known genes implicated in focal vs. generalized epilepsy were enriched in NAFE vs. GGE. Interestingly, the same pattern was found for the 100 top-ranking genes associated with GWAS hits, which were preferentially enriched in respective phenotypic groups. The enrichment of rare variants in GWAS genes also supports the convergence of ultra-rare and common variants in conferring epilepsy risk. According to our findings, a link between common and rare variants is likely to be also relevant for the phenotypic heterogeneity observed in seizure disorders. Notably, polygenic risk scores also pointed out the specificity of the risk profiles in common epilepsies (68). We also found an enrichment of ultra-rare *de novo* variants in DEE in a previously identified brain co-expression module (54), and the same module was also enriched for constrained variants in GGE and NAFE. Although co-expressed genes are not necessarily part of a single pathway, they represent closely orchestrated networks with possible functional correlations. It is therefore conceivable that differentially expressed genes in individuals with epilepsy would highlight modules in which altered transcription, ultra-rare variants, or both contribute to cause both rare and common epilepsies.

### Study limitations

Despite their robustness, the associations presented in this work should be interpreted with the caveats of gene group testing in mind (79). Given that pathways and molecular processes are not consistently defined in different resources, it is not always easy to define genes that represent a certain pathway. For instance, GABAergic and glutamatergic pathway definitions based on GO terms and KEGG databases are widely discordant (Fig. S18). These differences may explain the discrepancies we observed in enrichment patterns in the same pathway. One gene with abundance of qualifying variants in the GO-based analysis of the glutamatergic pathway was *PPFIA3* (see Additional file 4), a highly intolerant gene that encodes a synaptic receptor tyrosine phosphatase highly expressed in the brain. However, *PPFIA3* is not part of the KEGG-based glutamatergic pathway. Attempting to overcome such discrepancies, we examined multiple overlapping gene-sets from different resources. Associations that are seen regardless of the gene-set source (e.g., the consistent pattern of enrichment in inhibitory vs. excitatory neuronal/pathway genes in GGE vs. NAFE) are, therefore, likely to underscore a genuine biological relevance.

The analysis presented here has additional limitations which we aimed to overcome using stringent analysis and quality control strategies. The limited use of about half of the controls from the primary analysis to maximize case-control matching affected the overall power. Nevertheless, we were able to reproduce most of the major signals from gene-sets with large effect sizes, the latter thereby active as positive controls. The male-to-female ratios were not well-balanced in our case and control sets. Based on a secondary analysis excluding X chromosomal genes, this does not seem to introduce a substantial bias (see Additional file 5). The overlap between the controls used in this study and gnomAD controls resulting from MIGen Leicester and Ottawa controls (Table S2) created some challenges in defining ultra-rare variants, usually defined as those variants not observed in population databases. For population frequency filtering, we allowed around five alleles in gnomAD (allele frequency of 2×10^-5^). This count that exceeds our internal filtering cut-off for ultra-rare variants (three alleles) would allow for the retention of ultra-rare variants from our control that are also seen in gnomAD while still filtering common variants and prevalent sequencing artifacts. Also, multiple in-silico algorithms for predicting missense deleteriousness and estimating constraint (including MPC, MTR, and CCR scores) were derived from or validated in the Exome Aggregation Consortium (80) and gnomAD databases. Examination of control conditions and control gene-sets that are not expected to show an enrichment (e.g., genes not expressed in the brain) did not indicate any prominent inflation and supported the validity of the overall analysis. Interestingly, some signals in ‘non-neuronal’ gene-sets could be explained by the inclusion of well-established epilepsy genes (Fig. S20), like *CACNA1A* (63) and *CACNA1E* (64), which are in fact key neuronal genes of synaptic transmission.

## Conclusions

Missense URVs affecting constrained sites in brain-expressed genes are key genetic drivers in epilepsy. Patients with both common and rare epilepsy subtypes show an increased exome-wide burden of such variants that is primarily derived from neuronal genes, where key gene-sets including ion channels, developmental and synaptic genes are enriched across the phenotypic spectrum of epilepsy. Genes implicated by common GWAS variants may also be disrupted by URVs in various epilepsy phenotypes, suggesting a convergence of rare disruptive variants, and common variants in the pathogenesis of epilepsy. Enrichment patterns of URVs-affected genes suggest a preferential involvement of inhibitory genes in GGE and excitatory genes in focal epilepsies.

## List of abbreviations

ARC: Activity-Regulated Cytoskeleton protein.
ATVB: Atherosclerosis, Thrombosis, and Vascular Biology Study.
CAD: Coronary Artery Disease.
CCR: Constrained Coding Regions.
CDS: Coding sequences.
dbGAP: Database of Genotypes and Phenotypes.
DEE: Developmental and Epileptic Encephalopathies.
FDR: False Discovery Rate.
FMRP: Fragile-X Mental Retardation Protein.
GATK: Genome Analysis Toolkit.
GGE: Genetic Generalized Epilepsy.
GO: Gene Ontology.
GWAS: Genome-Wide Association Study.
KEGG: Kyoto Encyclopedia of Genes and Genomes.
ILAE: International League Against Epilepsy.
MAC: Minor Allele Counts.
MAF: Minor Allele Frequencies.
MGI: Mouse Genome Informatics.
MIGen: Myocardial Infarction Genetics Consortium.
MPC: Missense Badness Polyphen and Constraint.
MTR: Missense Tolerance Ratio.
NAFE: Non-Acquired Focal Epilepsy.
NDD: Neuro-Developmental Disorders.
NMDA: N-methyl D-Aspartate.
PCA: Principal Component Analysis.
PCs: Principal Components.
PPh2: PolyPhen2
PSD-95: Post-synaptic density protein 95.
PTVs: Protein Truncating Variants.
QVs: Qualifying Variants.
SIFT: Sorting Intolerant From Tolerant
URVs: Ultra-Rare Variants.

## Declarations

### Ethics approval and consent to participate

This analysis utilized previously analyzed and published data from the Epi25 Collaborative and the MIGen Consortium. Subjects investigated by the Collaborative provided signed informed consent at the participating centers according to local national ethical requirements and their standards at the time of collection. Approval for data reuse and analysis was obtained from the Epi25 Collaborative (cases) and dbGAP (controls). The ethical approval and consents procedures for the individual cohorts were reported by the Epi25 Collaborative (see methods section).

### Consent for publication

not applicable.

### Availability of data and materials

The data/analyses presented in the current publication are based on the use of study data from the Epi25 Collaborative (http://epi-25.org/) and the dbGaP (https://ncbi.nlm.nih.gov/gap/) under accessions phs000473, phs001000, phs000806, phs001592 and phs001489. Data supporting the results reported in the article can be found in this published article and its supplementary information files.

### Authors’ contributions

HL, PM, MK conceived the study. HL, PM, RK, and MK contributed to the study design and/or data acquisition. MK and PM performed the analysis with input from HL, RK, TS, DRB and MN. MK, PM, and HL interpreted the results and wrote the manuscript. Authors from the Epi25 Collaborative recruited and phenotyped patients and/or contributed to data generation as indicated in the appendix. All authors reviewed the manuscript for intellectual content and approved the final version of the manuscript.

## Acknowledgements

We are cordially thankful to the Epi25 Collaborative staff and all patients participating in the Epi25 project, who all made this study possible.

## Epi25 Collaborative

Yen-Chen, Anne, Feng. Daniel, P, Howrigan. Liam, E, Abbott. Katherine, Tashman. Felecia, Cerrato. Tarjinder, Singh. Henrike, Heyne. Andrea, Byrnes. Claire, Churchhouse. Nick, Watts. Matthew, Solomonson. Dennis, Lal. Erin, L, Heinzen. Ryan, S, Dhindsa. Kate, E, Stanley. Gianpiero, L, Cavalleri. Hakon, Hakonarson. Ingo, Helbig. Roland, Krause. Patrick, May. Sarah, Weckhuysen. Slavé, Petrovski. Sitharthan, Kamalakaran. Sanjay, M, Sisodiya. Patrick, Cossette. Chris, Cotsapas. Peter, De Jonghe. Tracy, Dixon-Salazar. Renzo, Guerrini. Patrick, Kwan. Anthony, G, Marson. Randy, Stewart. Chantal, Depondt. Dennis, J, Dlugos. Ingrid, E, Scheffer. Pasquale, Striano. Catharine, Freyer. Kevin, McKenna. Brigid, M, Regan. Susannah, T, Bellows. Costin, Leu. Caitlin, A, Bennett. Esther, M C, Johns. Alexandra, Macdonald. Hannah, Shilling. Rosemary, Burgess. Dorien, Weckhuysen. Melanie, Bahlo. Terence, J, O’Brien. Marian, Todaro. Hannah, Stamberger. Danielle, M, Andrade. Tara, R, Sadoway. Kelly, Mo. Heinz, Krestel. Sabina, Gallati. Savvas, S, Papacostas. Ioanna, Kousiappa. George, A, Tanteles. Katalin, Štěrbová. Markéta, Vlčková. Lucie, Sedláčková. Petra, Laššuthová. Karl, Martin, Klein. Felix, Rosenow. Philipp, S, Reif. Susanne, Knake. Wolfram, S, Kunz. Gábor, Zsurka. Christian, E, Elger. Jürgen, Bauer. Michael, Rademacher. Manuela, Pendziwiat. Hiltrud, Muhle. Annika, Rademacher. Andreas, van Baalen. Sarah, von Spiczak. Ulrich, Stephani. Zaid, Afawi. Amos, D, Korczyn. Moien, Kanaan. Christina, Canavati. Gerhard, Kurlemann. Karen, Müller-Schlüter. Gerhard, Kluger. Martin, Häusler. Ilan, Blatt. Johannes, R, Lemke. Ilona, Krey. Yvonne, G, Weber. Stefan, Wolking. Felicitas, Becker. Christian, Hengsbach. Sarah, Rau. Ana, F, Maisch. Bernhard, J, Steinhoff. Andreas, Schulze-Bonhage. Susanne, Schubert-Bast. Herbert, Schreiber. Ingo, Borggräfe. Christoph, J, Schankin. Thomas, Mayer. Rudolf, Korinthenberg. Knut, Brockmann. Gerhard, Kurlemann. Dieter, Dennig. Rene, Madeleyn. Reetta, Kälviäinen. Pia, Auvinen. Anni, Saarela. Tarja, Linnankivi. Anna-Elina, Lehesjoki. Mark, I, Rees. Seo-Kyung, Chung. William, O, Pickrell. Robert, Powell. Natascha, Schneider. Simona, Balestrini. Sara, Zagaglia. Vera, Braatz. Michael, R, Johnson. Pauls, Auce. Graeme, J, Sills. Larry, W, Baum. Pak, C, Sham. Stacey, S, Cherny. Colin, H T, Lui. Nina, Barišić. Norman, Delanty. Colin, P, Doherty. Arif, Shukralla. Mark, McCormack. Hany, El-Naggar. Laura, Canafoglia. Silvana, Franceschetti. Barbara, Castellotti. Tiziana, Granata. Federico, Zara. Michele, Iacomino. Francesca, Madia. Maria, Stella, Vari. Maria, Margherita, Mancardi. Vincenzo, Salpietro. Francesca, Bisulli. Paolo, Tinuper. Laura, Licchetta. Tommaso, Pippucci. Carlotta, Stipa. Raffaella, Minardi. Antonio, Gambardella. Angelo, Labate. Grazia, Annesi. Lorella, Manna. Monica, Gagliardi. Elena, Parrini. Davide, Mei. Annalisa, Vetro. Claudia, Bianchini. Martino, Montomoli. Viola, Doccini. Carla, Marini. Toshimitsu, Suzuki. Yushi, Inoue. Kazuhiro, Yamakawa. Birute, Tumiene. Lynette, G, Sadleir. Chontelle, King. Emily, Mountier. S, Hande, Caglayan. Mutluay, Arslan. Zuhal, Yapıcı. Uluc, Yis. Pınar, Topaloglu. Bulent, Kara. Dilsad, Turkdogan. Aslı, Gundogdu-Eken. Nerses, Bebek. Sibel, Uğur-İşeri. Betül, Baykan. Barış, Salman. Garen, Haryanyan. Emrah, Yücesan. Yeşim, Kesim. Çiğdem, Özkara. Annapurna, Poduri. Beth, R, Shiedley. Catherine, Shain. Russell, J, Buono. Thomas, N, Ferraro. Michael, R, Sperling. Warren, Lo. Michael, Privitera. Jacqueline, A, French. Steven, Schachter. Ruben, I, Kuzniecky. Orrin, Devinsky. Manu, Hegde. Pouya, Khankhanian. Katherine, L, Helbig. Colin, A, Ellis. Gianfranco, Spalletta. Fabrizio, Piras. Federica, Piras. Tommaso, Gili. Valentina, Ciullo. Andreas, Reif. Andrew, McQuillin. Nick, Bass. Andrew, McIntosh. Douglas, Blackwood. Mandy, Johnstone. Aarno, Palotie. Michele, T, Pato. Carlos, N, Pato. Evelyn, J, Bromet. Celia, Barreto, Carvalho. Eric, D, Achtyes. Maria, Helena, Azevedo. Roman, Kotov. Douglas, S, Lehrer. Dolores, Malaspina. Stephen, R, Marder. Helena, Medeiros. Christopher, P, Morley. Diana, O, Perkins. Janet, L, Sobell. Peter, F, Buckley. Fabio, Macciardi. Mark, H, Rapaport. James, A, Knowles. Consortium, (GPC), Genomic Psychiatry Cohort. Ayman, H, Fanous. Steven, A, McCarroll. Namrata, Gupta. Stacey, B, Gabriel. Mark, J, Daly. Eric, S, Lander. Daniel, H, Lowenstein. David, B, Goldstein. Holger, Lerche. Samuel, F, Berkovic. Benjamin, M, Neale.

## Authors’ information

See Additional file 1 for the affiliations of the Epi25 Collaborative members.

## Additional files

- Additional file 1: PDF file. Supplemental methods, tables and figures; affiliations of the Epi25 Collaborative members.

- Additional file 2: Excel xlsx file. Tables of gene-sets and genes in each gene-sets.

- Additional file 3: Excel xlsx file. Gene-set burden analysis results.

- Additional file 4: Excel xlsx file. Top-ranking genes per gene-set in the CCR 80 analysis.

- Additional file 5: Excel xlsx file. Secondary analysis results.

