## Additional file 1 for "Ultra-rare constrained missense variants in the epilepsies: Shared and specific enrichment patterns in neuronal gene-sets"

**Additional file 1: Supplementary methods, tables, figures and author information.**

*Supplementary Methods:* ... *3*

Sample cohorts overview: ... 3

Baseline sample quality control: ... 3

Baseline variant quality control: ... 5

Residual stratification: ... 5

Qualifying Variants: ... 6

Burden testing in gene groups: ... 7

References: ... 9

*Supplementary tables and figures:* ... *12*

Table S1: Epilepsy samples analyzed in this study ... 12

Table S2: Control datasets analyzed in this study ... 13

Table S3: Summary of baseline sample-level quality control. ... 14

Table S4: Final sample counts. ... 15

Table S5: Final variant statistics. ... 15

Table S6: Variant types and conditions used for the gene group burden analysis. ... 16

Table S7: Gene-sets ... 17

Table S8: Genes in each gene-set ... 17

Table S9: Burden of qualifying variants in 92 gene-sets. ... 17

Table S10: Top-ranking genes per gene-set. ... 17

Table S11: Secondary gene-set burden analysis results. ... 17

Fig. S1: Variant calling metrics of sequencing cohorts grouped by capture kits. ... 18

Fig. S2: Heterozygosity and kinship filtering. ... 18

Fig. S3: Continental ancestry groups. ... 20

**Supplementary Methods:**

**Sample cohorts overview:**

Access to two sets of variant calls (separate but jointly called VCF files) mapped to the human genome build GRCh37 was granted by the Epi25 Collaborative (1). The first set (*n*=13,497) contained calls from patients (*n*=13,197) and controls (*n*=300) collected by the Epi25 Collaborative. The second set (*n*=12,999) included controls from the Swedish Schizophrenia Study (S-SCZ), patients and controls from the Myocardial Infarction Genetics (MIGen) Consortium cohorts with access permission granted from dbGAP (2). The data generation process has been previously described (1). The exome sequencing was performed on an Illumina HiSeq 2000 or 2500 (Illumina, USA) at the Broad Institute (different patches or timepoints) and utilized Illumina TruSeq/Nextera (Epi25), Illumina's ICE Capture (MIGen), or Agilent SureSelect Human All Exon Kits (MIGen and S-SCZ) (Agilent, USA). Following quality control and harmonization steps outlined hereafter, 58% of the initial cases (table S1) and 30% of the control samples (table S2) were included in the final analysis.

**Baseline sample quality control:**

Cases with a diagnosis other than a Developmental and Epileptic Encephalopathy (DEE), a Genetic Generalized Epilepsy (GGE) or a Non-Acquired Focal Epilepsy (NAFE) were removed. The case definitions from the Epi25 Collaborative can be accessed online (http://epi-25.org/epi25-data-checks). Controls from MIGen cohorts with a coronary artery disease were not included in the analysis to avoid any prominent overlap in genetic predisposition. Gencode coding sequence (CDS) boundaries (v33 lifted to b37) (3) were padded with 10 bp and masked for low complexity and repeat regions (stratification files v. March 9, 2017) obtained from the Global Alliance for Genomics and Health (4) using bedtools v2.29.2 (5). All subsequent sample quality control, variant quality control and final analysis was performed over these regions (totaling 38Mb). The variant calling metrics were gathered for the two datasets over the CDS boundaries described above using the Genome Analysis ToolKit v4.1.4.1 (gatk CollectVariantCallingMetrics) (6). Outliers beyond 4 absolute deviations on total snps/indels counts, TiTv ratio, Ins-Del ratio, or Hom-Het ratio (per cohort) were filtered (Fig. S1). The VCF files were converted to PLINK v1.9 binaries (plink --vcf --make-bed) and merged (plink --bmerge)

(7). The genotyping rate (plink --missing) per sample was then calculated over the target CDS boundaries. Samples with genotyping rate less than 90% were filtered (Fig. S1). PLINK was used to select a set of informative SNPs with missingness less than 0.01, minor allele frequency exceeding 0.05, and in Hardy-Weinberg Equilibrium (plink --snps-only --maf 0.05 --geno 0.01 --hwe midp include-nonctrl 10e-6). These were then pruned (plink --indep-pairwise 50 5 0.5) and used to estimate autosomal heterozygosity (plink --het) using the F-statistic. Outliers beyond 3 standard deviations were filtered (Fig. S2). Informative SNPs (as detailed above) located on chrX were split (plink --split-x b37), pruned, and then used to estimate the F-statistic over chrX (plink --check-sex). Following visualization, cut-offs of F <= 0.2 (females) and F >= 0.6 (males) were used for SNP-sex prediction. Samples with ambiguous (0.2 < F < 0.6) or discordant sequencing and reported sex were filtered (figure S2). KING v2.2.4 (8) was used to detect duplicate samples and estimate the relatedness (king --related –degree 3). For each pair from duplicates and related samples up to the 3rd degree (Fig. S2), the sample with the lower genotyping rate was filtered. KING was used to perform multidimensional scaling (MDS) (5 principal components) on genotyping data from 2,451 samples from 1000 Genomes Project phase 2 followed by projection of the case and control samples into the 1000 Genomes space (king --mds --projection) (9). A subset of variants (*n*=73,080) that are called both in the 1000 Genomes data and our dataset were selected for projection. The five principal components from a randomly selected subset of 1000 Genomes samples (80% of samples) were used to train a Support Vector Machine (SVM), as implemented in R package *e1071* (10). A radial kernel was used to recognize four major continental ancestry groups: non-Finnish Europeans “EUR” (excluding “FIN” population), African “AFR”, admixed American “AMR”, South and East Asian “ASI” (including “EAS” and “SAS” super-populations). The SVM was tested on the remaining 1000 Genomes samples (20%), were it correctly recalled all samples with European ancestry, then used to classify the cases and control study samples (Fig. S3). Samples with a predicted ancestry other than European were filtered. These filtering steps removed 7,511 samples. To maximize the case-control matching among the remaining 18,985 samples, MDS (10 principal components) was repeated on a subset of samples 1000 Genomes of European ancestry (*n*=500, Northern and Western Europeans from Utah “CEU”, British in England and Scotland “GBR”, Toscani in Italy “TSI”, Iberian from Spain “IBS”, Finnish in Finland “FIN”), with projection of those samples labeled as European by the SMV (variants selected as indicated above) on this MDS space of European 1000 Genomes populations (Fig. S3). Upon visualization of the first two principal components, samples clustering with Finnish Europeans were removed (PC1 > 0.04). The Euclidean distance between all pairs of remaining case and control samples (on PC1/PC2) was calculated. Outlier samples (beyond 3 median absolute deviations) were filtered. The final set of baseline-filtered samples (Fig. S4) constituted 7,836 cases and 8,822 controls (*n*=16,661) as detailed in table S3).

**Baseline variant quality control:**

Two VCFs (1) containing 6,429,324 jointly called sites annotated with variant quality scores log odds (VQS Lod) were merged using bcftools/htslib v1.10.2 (11) and sites located outside the target CDS boundaries (see above) and sites with low recalibrated variant quality scores (SNPs VQS Lod < -3.0998 and Indel VQS Lod < 0.8107 corresponding to VQSR sensitivity tranche 99.6 and 95.00, respectively) were filtered (bcftools merge -f "PASS,." -R; bcftools view -c1 -S). The variants were allele-split and normalized using bcftools (bcftools norm -w 500 -c w) and vt v0.57721 (vt sort -m local) (12). The merged and normalized VCF was subset to the baseline filtered samples identified as detailed above (bcftools view -c1 -S). Genotype calls with depth < 10, quality < 20, or half-missing calls were set to missing (bcftools +setGT). Heterozygous genotypes with allele depth to total depth ratio < 0.25 were set to reference calls (bcftools +setGT). Variants with allele count equal to 0 were removed (bcftools view -c 1). These filtering steps were performed on a binary VCF steam piped between the outlined commands. Afterwards, the depth of coverage per variant was calculated (bcftools query -f '[%FORMAT\tDP]\n'). Only variants covered at a minimum depth of 10x in 95% of the baseline filtered cases and control sets were kept. Additionally, the distribution of the difference in mean coverage and the percentage of samples covered at depth 10x was visualized. All outlier variants beyond 3 standard deviations were filtered. The statistical calculations were performed in R (13). This quality control process resulted in a well-harmonized coverage between cases and controls (Fig. S4).

**Residual stratification:**

To maximize the cohort, sample and variant matching, we performed multiple rounds of principal component analyses (PCA) coupled with coverage harmonization among cohorts. To remove poorly matching sample cohorts, a baseline round of PCA (10 principal components) was performed using

PLINK (plink --pca) on a set of pruned variants (pruning was performed as described for baseline filtering above). A cohort of Swedish controls (*n*=4,838) clustered poorly with the rest of the study samples on the top principal components (PC1/2) (Fig. S5) and was therefore excluded. We then calculated the variant call rates (bcftools +fill-tags) across the remaining cohorts (Epi25, Leicester, Ottawa, ATVB), and removed all variants where any given cohort had a coverage < 95% or if the difference in coverage between any two given cohorts exceeded 0.5%. Variants not in Hardy-Weinberg equilibrium (p-value $< 10^{-6}$) were identified (plink --hwe) and filtered. This filtering insured that the top principal components will capture the ancestry and not the exome capture kits differences (Fig. S5). A second round of PCA (10 principal components) using EIGENSTRAT v6.1.4 (14,15) was performed (smartpca -p; outlier vectors = 2, outliers sigma = 6, iterations = 5) complemented by removal of extreme outliers identified upon visual inspection (PC1/PC2). A small subset of poorly matched samples (*n*=272) was subsequently removed. A third and final round of PCA with identical EIGENSTRAT parameters showed a well-matched case-control cohort (Fig. S5). The variant calling metrics were balanced for this set (gatk CollectVariantCallingMetrics) (Fig. S6).

**Qualifying Variants:**

Variant effects and consequences were annotated using snpEff v4.3t (16). Annovar v20191024 (17) was then used to annotate population frequencies from gnomAD r2.1 (18) and DiscovEHR Freeze 50 (19) as well as the following missense *in-silico* pathogenicity predictions: Sorting Intolerant From Tolerant (SIFT) (20), PolyPhen2 (PPh2) Human Diversity-based predictions (21), Missense-badness PolyPhen2 and Constraint (MPC) score (22), Missense Tolerance Ratio (MTR) score (23), and Paralog conservation (para-Z) score (24). Consensus Coding Region (CCR) scores (25) were annotated using tabix (26). Ultra-rare variants (URVs) were defined as follows: 1. Allele Count ($AC_{Analysis}$) <= 3, where $AC_{Analysis} = AC_{Epilepsy\text{-}type} + AC_{Controls}$ (epilepsy types: DEE, GGE, or NAFE depending on the analysis). 2. Not present in DiscovEHR ($MAF_{DiscovEHR} = 0$). 3. Allele Count <= 5 in gnomAD ($MAF_{gnomAD} < 2x10^{-5}$). The inclusion of gnomAD variants with low frequency allowed the use of control sets that overlap with gnomAD (table S2), since gnomAD variants are filtered at a higher count (5 alleles) compared to the analysis set allele count (3 alleles). The ultra-rare variants were grouped in thirteen analysis conditions as detailed in table S6. The genotypes and annotations were queried using bcftools or snpEff and imported for statistical analysis in R v3.3 (13). These were collapsed in a dominant model (reference as 0, heterozygous, homozygous and hemizygous as 1) to obtain a matrix of samples vs. genes where the cells contained 0/1 indicators for the presence or absence of a qualifying variant (QV) in a specific sample and gene. Single gene collapsing analysis was performed using Fisher Exact Test (FET). The genomic inflation was estimated by performing a gene-level collapsing analysis for synonymous qualifying variants using FET. *P* values obtained from gene-level FET were compared to a permutation-based distribution obtained from 1000 permutations (shuffling of case-control labels followed by FET). This was performed with a parallel implementation of the *QQ-perm* method (27) using *parallel* package (13). The Genomic Inflation Factor (λ) was estimated using *QQ-perm*. The negative $\log_{10}$ of the observed *p* values was plotted against the negative $\log_{10}$ of the mean permutation *p* values to obtain the *Quantile-Quantile* plots shown in Fig. S7.

**Burden testing in gene groups:**

In total, 92 gene groups, presented in table S7 (see Additional file 2), were tested. The genes in each gene set are given in table S8 (see Additional file 2). The construction of gene groups leveraged multiple sources as detailed in Additional file 2. To ensure homogeneity between obtained gene groups and snpEff annotations, each gene set was limited to those genes annotated with snpEff as protein coding genes using Ensembl gene IDs on GRCh37. Where available, Ensembl gene IDs were obtained from sources of gene sets. Otherwise, *biomaRt* package (28) and gProfiler (29) were used to map Human Gene Nomenclature Consortium (HGNC) names and gene name synonyms to their Ensembl gene IDs. *biomaRt* was used to map mouse genes to their human paralogues. For each of the three phenotypic groups and analysis conditions, the qualifying variants tables were filtered for variants located one of the genes in the gene set under consideration. These were then added per gene group to calculate a group-level burden scores per sample (sample vs. QV burden table). Additional sample-level metrics were annotated (phenotype, sample sex, exome wide singletons and variant counts, and ten principal components per sample). These data handling steps were performed using R base, *data.table* (30) and *tidyverse* (31). The resulting table (sample vs. phenotypes, QV burden scores and covariates) was used as input to perform logistic regression. The case-control status (indicator variable) was regressed on covariates only (null model) or burden scores and covariates (test model) as additive predictors using *glm*(family=binomial) function from *stats* package (13). The null model was *glm*(sex + variant counts + singletons + PC1…PC10) and the test model was *glm*(QV burden + sex + variant counts + singletons + PC1… PC10). Likelihood Ratio Test (LRT) from *lmtest* package (32) was used to compare the test and null models. The LRT log-odds and their 95% confidence intervals were not corrected for multiple testing. *P* values of twelve test conditions were adjusted using Benjamini and Hochberg false discovery rate (FDR) method as implemented in *p.adjust*(method = "BH") from *stats* package. In total, FDR adjustments accounted for 3312 tests (92 gene sets x 12 conditions x 3 phenotypes). The *p* values for the synonymous condition were not FDR-adjusted. A secondary analysis was performed over the 92 gene-sets limited to autosomal genes (excluding all genes on chromosome X). The aim was to estimate the bias created by male-to-female ratios imbalance (table S4). Another secondary analysis was performed using MIGen Leicester samples (Illumina ICE capture kits) as cases MIGen Ottawa/ATVB samples as controls (Agilent SureSelect capture kits) to exclude the presence of significant residual stratification between capture kits (table S2). The results of these secondary analysis are presented in Additional file 5. Comparisons between samples prepared using Illumina Nextera/TruSeq and Illumina ICE or Agilent SureSelect were not performed as these are almost identical to the primary cases (Nextera/TruSeq) vs. controls (ICE & SureSelect) analysis (Fig. S5).

We presumed equal weights and direction of effects for the variants in the conditions under analysis by taking the sum of qualifying variants in a specific gene group per sample as a predictor for a binary phenotype in a regression model. While this assumption is fairly reasonable for highly deleterious and constrained variants, it is rather simplistic for milder genetic alterations. This approach is also not ideal to estimate the odds in data sets with low counts. However, the computational ease, the clarity in setting up the analysis parameters in comparison to other variance componence-based and hybrid methods, e.g., skat-o (33), are key advantages that motivated this choice. The use of similar regression models has been shown to capture the major signals in gene-set burden analysis in epilepsy and other neurological diseases (1,34,35).

**Supplementary tables and figures:**

**Table S1: Epilepsy samples analyzed in this study.**

| **Phenotype group** | **Total** | **Phenotype review** | **Initial QC** | **Final QC** |
|---|---|---|---|---|
| Developmental and Epileptic Encephalopathy | 1,474 | 1,467 | 1,040 (71%) | 1,003 (68%) |
| Genetic Generalized Epilepsy | 4,510 | 4,471 | 3,183 (71%) | 3,064 (68%) |
| Non-Acquired Focal Epilepsy | 5,321 | 5,304 | 3,616 (68%) | 3,522 (66%) |
| Febrile Seizures and GEFS spectrum | 301 | Not considered | | |
| Symptomatic / Lesional | 1,434 | Not considered | | |
| Other epilepsies, unclassified epilepsies, non-epileptic seizures or not available | 157 | Not considered | | |
| Total | 13,197 | 11,242 | 7,839 (59%) | 7,589 (58%) |

**Table S2: Control datasets analyzed in this study.**

| Control set | Capture kits | In dbGAP | In gnomAD | Phenotype | Total | Initial QC | Final QC |
|---|---|---|---|---|---|---|---|
| Epi25 Collaborative controls (Italy) | Illumina TruSeq/Nextera | No | No | Unaffected | 300 | 283 (94%) | 283 (94%) |
| Leicester Heart study (UK) | Illumina ICE | phs001000.v1.p1 | Yes | Unaffected | 1,100 | 1082 (98%) | 1,082 (98%) |
| | | | | Coronary Artery Disease | 65 | - | - |
| Ottawa Heart Study (Canada) | Agilent SureSelect | phs000806.v1.p1 | Yes | Unaffected | 987 | 946 (96%) | 924 (94%) |
| | | | | Coronary Artery Disease | 928 | - | - |
| Atherosclerosis Thrombosis & Vascular Biology study (Italy) | Agilent SureSelect | phs001592.v1.p1 | No | Unaffected | 1,802 | 1,673 (93%) | 1,673 (93%) |
| | | | | Coronary Artery Disease | 1,875 | - | - |
| Swedish Schizophrenia Study (Sweden) | Agilent SureSelect | phs000473.v2.p2 | Yes | Unaffected | 6,242 | 4,838 (78%) | - |
| Total | | | | | 13,299 | 8,822 (66%) | 3,962 (30%) |

**Table S3: Summary of baseline sample-level quality control.**

| Criteria | Filter | Failing/Total (%) |
|---|---|---|
| Phenotype | Cases other than DEE, GGE, NAFE | 1,773/13,197 (13.4%) |
| | Controls with cardiac phenotype | 2,867/13,299 (21.6%) |
| Variant calling metrics | Outliers > 4 absolute deviations on key metrics | 1,088/26,496 (4.1%) |
| Genotyping rate | < 90% in called variants in coding regions | 66/26,496 (0.2%) |
| Autosomal heterozygosity | Outliers > 3 standard deviations | 1011/26,496 (3.8%) |
| ChrX heterozygosity | 0.2 < F < 0.6 or discordant reported/predicted sex | 255/26,496 (1.0%) |
| Kinship | Duplicate, twin or related up to the $3^{rd}$ degree | 331/26,496 (1.2%) |
| Major continental ancestry | Non-European ancestry predictions from SVM trained on 1000 Genomes samples | 2,057/26,496 (7.8%) |
| Samples failing one or multiple filters | | 7,511/26,496 (28.3%) |
| | | **18,985 samples remaining** |
| Matching | Finnish (MDS)/outliers on PCA (PC1/2) | 2,324/18,985 (12.2%) |
| | | **16,661 samples remaining** |

**Table S4: Final sample counts.**

| Group | Cohort | Samples | | Females % |
|---|---|---|---|---|
| DEE | | 1003 | | 462 (46%) |
| GGE | | 3064 | | 1764 (57.6%) |
| NAFE | | 3522 | | 1841 (52.3%) |
| Controls | Epi25 | 283 | 3962 | 767 (19.4%) |
| | Ottawa | 924 | | |
| | Leicester | 1082 | | |
| | ATVB | 1673 | | |

**Table S5: Final variant statistics.**

| Category | | Count |
|---|---|---|
| Jointly called sites in 26,496 samples | | 6,481,248 |
| Baseline QC:In CDS regions, normalized, genotype-filtered, VQSLOD filtered, not in low-complexity regions, AC > 0 in baseline-filtered samples | | 2,224,099 |
| Baseline variant harmonization: depth and call-rate harmonization | | 1,674,222 |
| Final QC: AC > 0 in final case-control set, cohort-level call rate harmonization, HWE | | 1,267,392 |
| Total variants | SNVs | 1,247,342 |
| | Indels | 20,050 |
| Variants with Allele Frequency < 0.5 % | | 1,203,350 |
| Variants with Allele Counts 1-3 | | 1,054,919 |
| Singleton variants | | 806,046 |

**Table S6: Variant types and conditions used for the gene group burden analysis.**

| Variant conditions | Control | | Deleterious variants | | | Missense constraint | | | | | Paralog conservation | | |
|---|---|---|---|---|---|---|---|---|---|---|---|---|---|
| Effects (Sequence Ontology terms) | Synonymous | Benign Missense | Damaging Missense | PTV | All Functional | MPC1 | MPC2 | MTR ClinVar | MTR DeNovo | CCR 80 | Paralog non-conserved | Paralog conserved | Paralog highly conserved |
| synonymous_variant | + | - | - | - | - | - | - | - | - | - | - | - | - |
| missense_variant | - | + | + | - | + | + | + | + | + | + | + | + | + |
| missense_variant (additional filters) | - | PPh2 & SIFT benign | PPh2 & SIFT damaging | - | PPh2 & SIFT damaging | MPC >=1 | MPC >= 2 | MTR <= 0.825 | MTR <= 0.565 | CCR >= 80 MPC >=1 MTR <= 0.825 | Para-Z-score =< 0 | Para-Z-score > 0 | Para-Z-score >= 1 |
| stop_gained | - | - | - | + | + | - | - | - | - | - | - | - | - |
| splice_acceptor_variant | - | - | - | + | + | - | - | - | - | - | - | - | - |
| splice_donor_variant | - | - | - | + | + | - | - | - | - | - | - | - | - |
| exon_loss_variant | - | - | - | + | + | - | - | - | - | - | - | - | - |
| frameshift_variant | - | - | - | + | + | - | - | - | - | - | - | - | - |
| start_lost | - | - | - | + | + | - | - | - | - | - | - | - | - |
| stop_lost | - | - | - | - | + | - | - | - | - | - | - | - | - |
| conservative_inframe_insertion | - | - | - | - | + | - | - | - | - | - | - | - | - |
| disruptive_inframe_insertion | - | - | - | - | + | - | - | - | - | - | - | - | - |
| conservative_inframe_deletion | - | - | - | - | + | - | - | - | - | - | - | - | - |
| disruptive_inframe_deletion | - | - | - | - | + | - | - | - | - | - | - | - | - |

**Table S7: Gene-sets.**

Provided in a separate excel file (Additional file 2).

**Table S8: Genes in each gene-set.**

Provided in a separate excel file (Additional file 2).

**Table S9: Burden of qualifying variants in 92 gene-sets.**

Provided in a separate excel file (Additional file 3).

**Table S10: Top-ranking genes per gene-set.**

Provided in a separate excel file (Additional file 4).

**Table S11: Secondary gene-set burden analysis results.**

Provided in a separate excel file (Additional file 5).

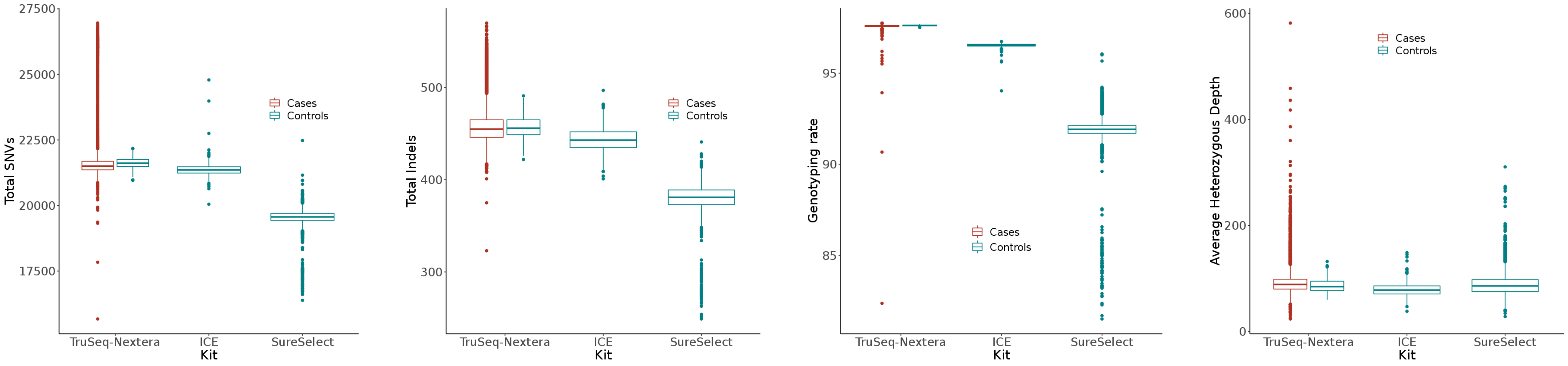

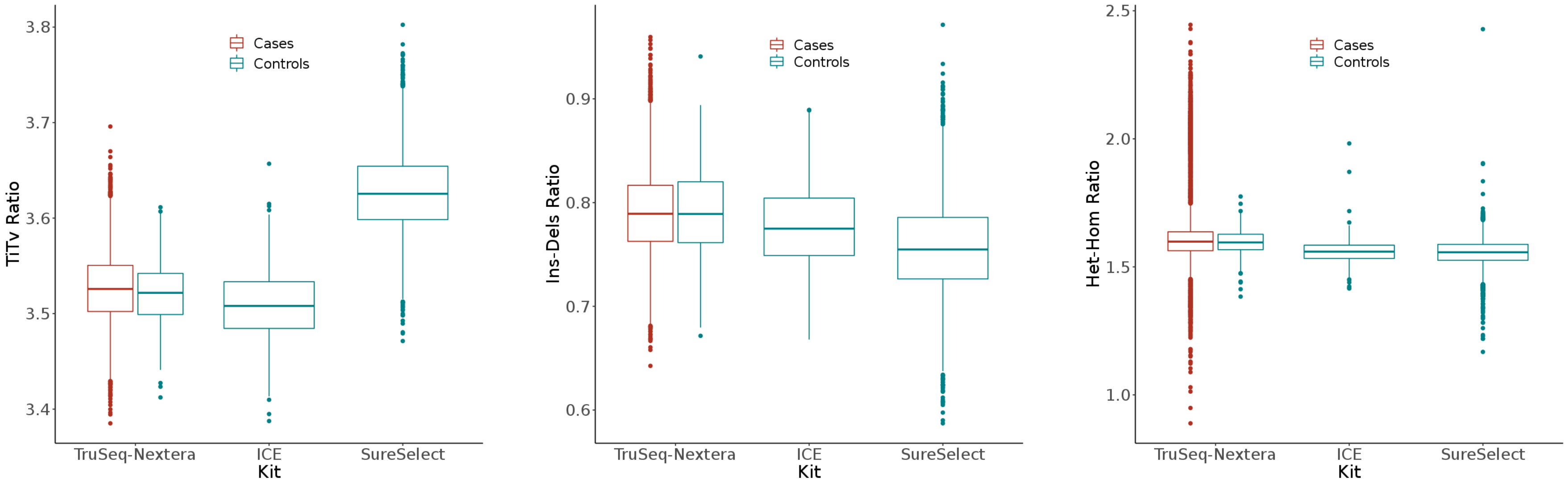

**Fig. S1: Variant calling metrics of sequencing cohorts grouped by capture kits.** These metrics were collected over Gencode coding regions, padded with 10 bps and masked for regions of repeats and low complexity. Samples with genotyping rate < 90% and outliers (> 4 absolute deviations per sequencing cohort) on SNV/Indel counts, TiTv ratio, Ins-Del ratio and Het-Hom ratio were filtered. SNVs: single nucleotide variations. Indels: insertions and deletions. TiTv: transitions-transversions. Ins-Del: insertions-deletions. Het-Hom: heterozygous-homozygous.

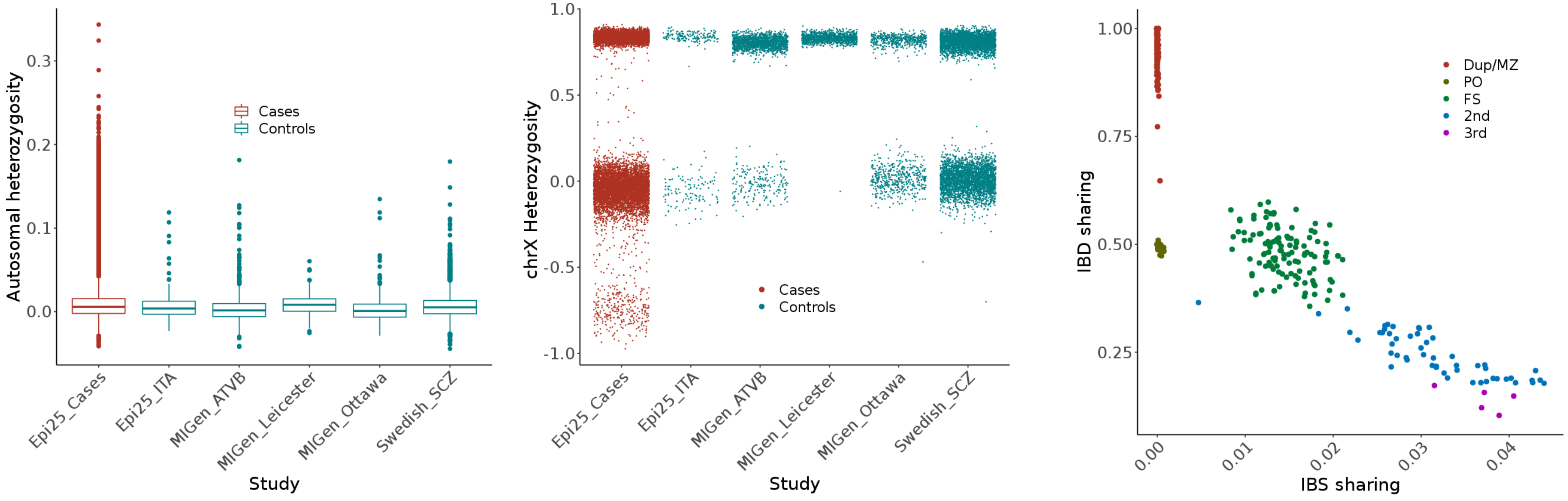

**Fig. S2: Heterozygosity and kinship filtering.** A set of common, pruned variants with high genotyping rate was used to calculate the F-statistic in autosomes (left) and chrX (center) using PLINK. Samples with low or excess autosomal heterozygosity (> 3 standard deviations) were filtered. For sex prediction (SNP-sex), cut-offs of 0.2 and 0.6 were used to separate female and male clusters from samples with ambiguous sequencing sex prediction. Integrated kinship predictions (right) using KING identified pairs of duplicates/twins and related samples. One sample from each pair was filtered. IBD: Identity by descent. IBS: identity by state. Dup/MZ: duplicates or monozygotic twins. PO: parent-offspring. FS: full-sibling. $2^{nd}$: second degree. $3^{rd}$: third degree.

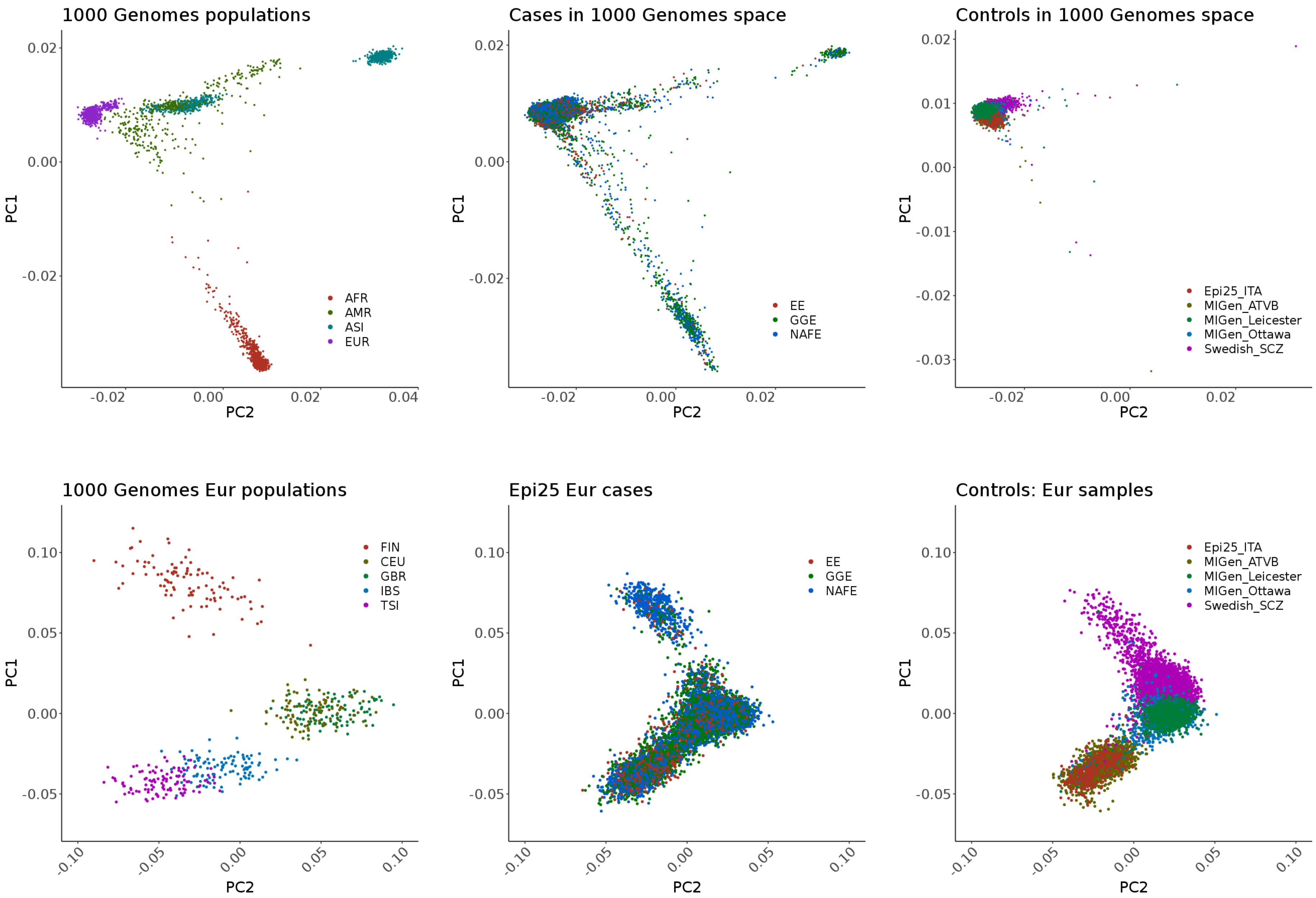

**Fig. S3: Continental ancestry groups.** KING multidimensional scaling (MDS) projection in the 1000 Genomes space (top left panel) was used to estimate the major ancestry components. The cases (top center panel) showed wide variability in continental ancestry. The controls (top right panel) were mostly of European ancestry. A support vector machine was trained on 1000 Genomes sample labels and used to identify Epi25 and control samples with likely European ancestry (bottom center and right panels). A second round of MDS was performed to project the principal components of 500 samples of European ancestry from the 1000 Genomes (bottom left) on the Epi25 cases and control samples classified as European (bottom middle and right panel) . See Fig. S4 for subsequent case-control matching.

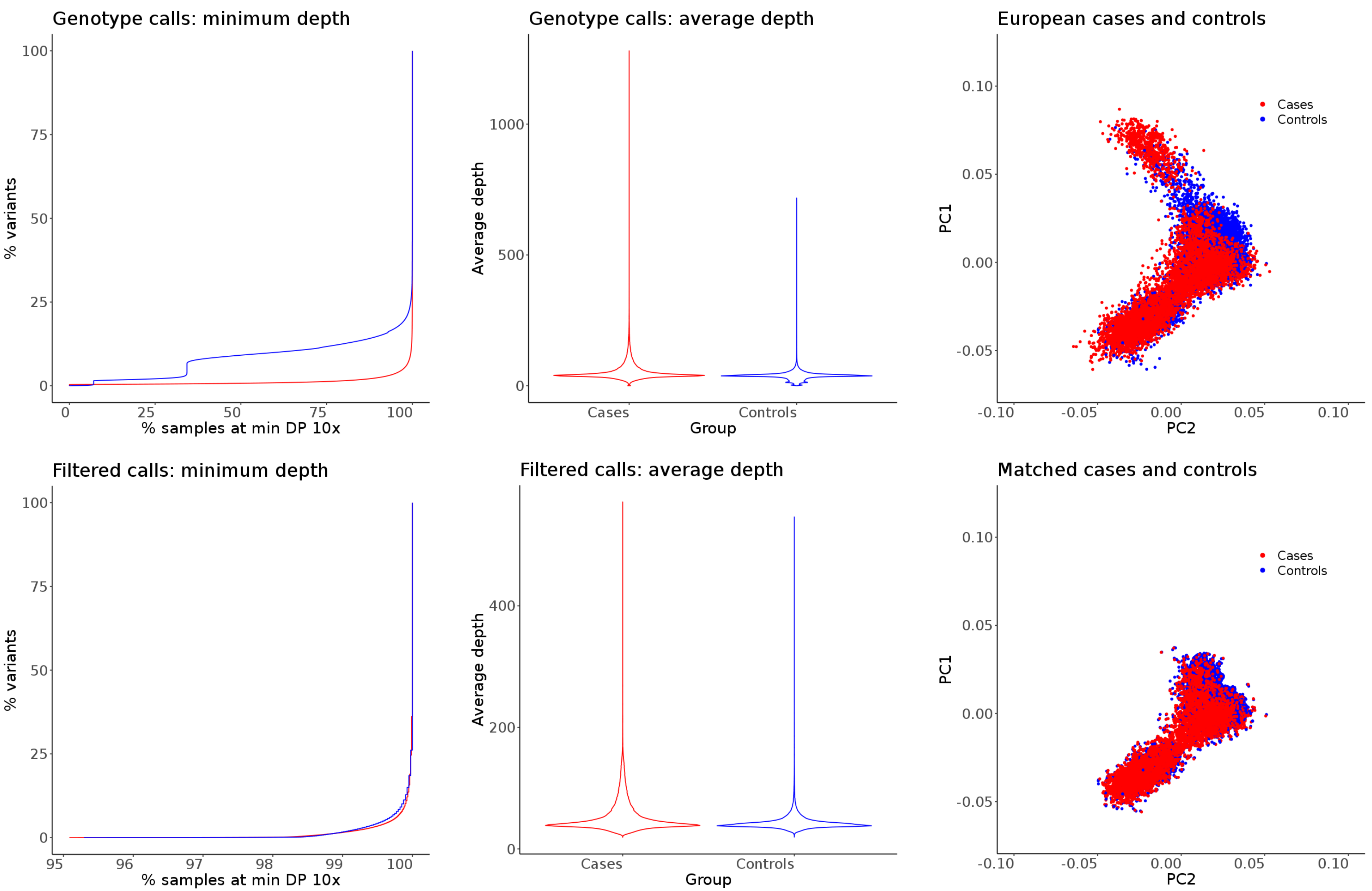

**Fig. S4: Baseline case-control matching and variant harmonization.** The precent of samples covered at a minimum depth of 10x (top left) and the average depth (top center) are shown for the cases (red) and controls (blue). Multidimensional scaling was used to estimate the major ancestral components (top-right; see Fig. S3 for details). To harmonize the ancestry and the variant calls, (a) about 20% of variants were removed where the percent of covered cases and controls was lower that 95%; (b) the difference in the average depth in cases and controls was calculated and outliers (> 3 standard deviations) were pruned out; (c) the difference in the percent of samples covered at depth 10x was calculated and variants with extreme differences (> 3 standard deviations) were also pruned; and (d) Poorly matched cases and controls on the top principal components PC1/PC2 and those of likely Finnish ancestry (PC1 > 0.04; see Fig. S3) were removed. This resulted in a homogeneous variant call rate (plots in bottom panels).

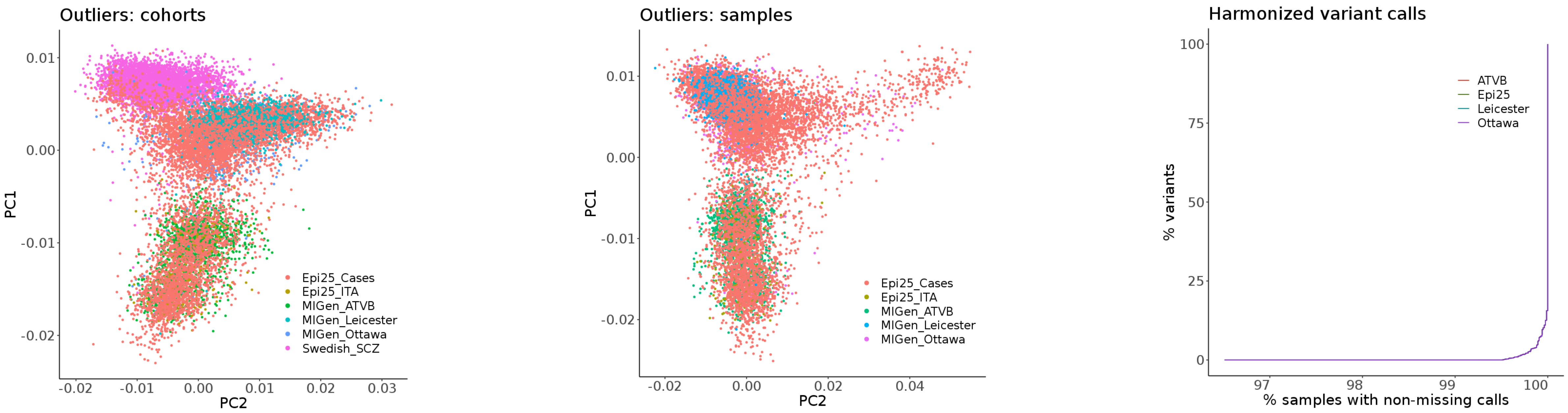

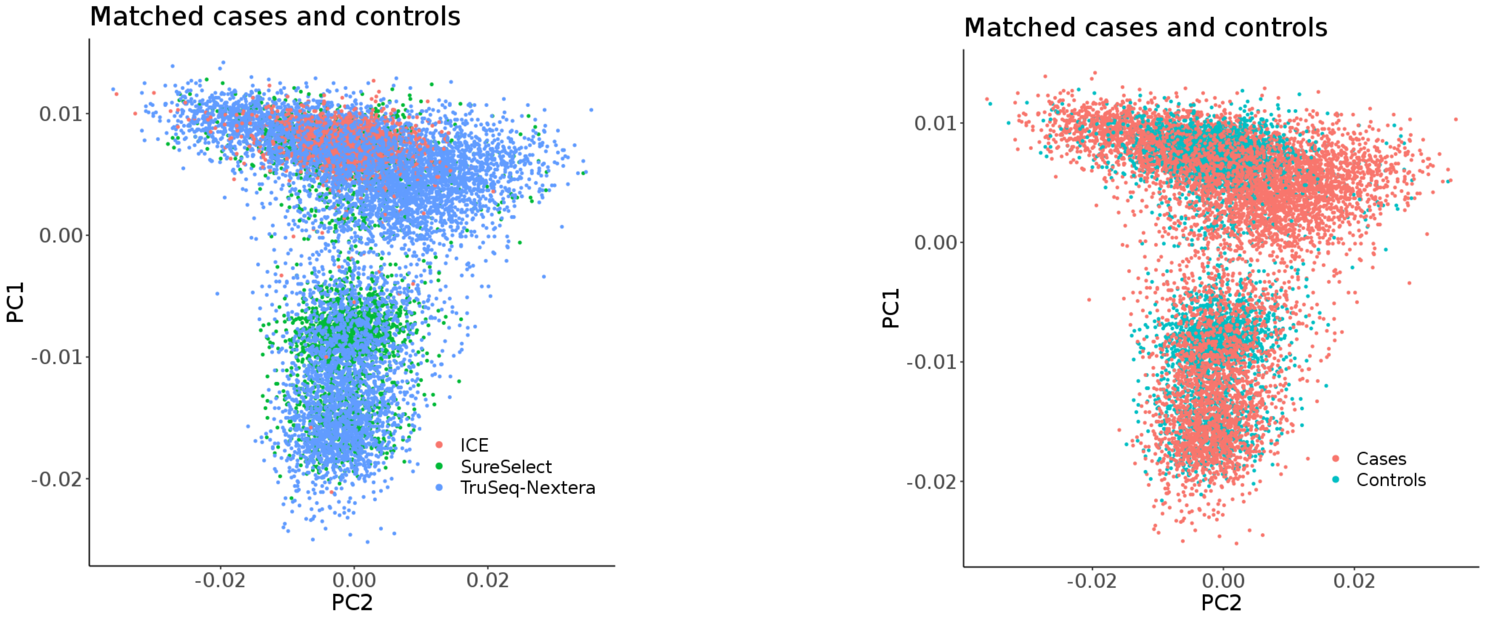

**Fig. S5: Final case-control matching.** Principal component analysis of baseline-filtered cases and controls showed residual population and cohort stratification (top left). Swedish controls (outliers on the first round of PCA; top left panel) and additional poorly matched samples (outliers on the second round of PCA; top center panel) were filtered. The call-rate was harmonized between different sequencing cohorts (top right) by removing all variants where the difference in call rate between pairs of individual cohorts exceeds 0.5%. These measures minimized the patch effects (bottom left). The first and second principal components of the final matched case control set (bottom center panel) capture the northern-southern and eastern-western European geographical axis, respectively

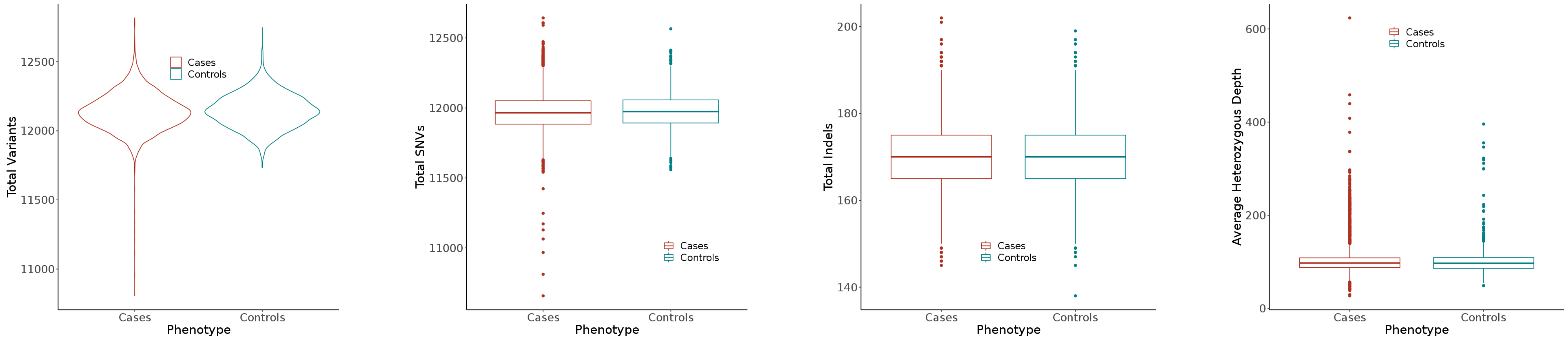

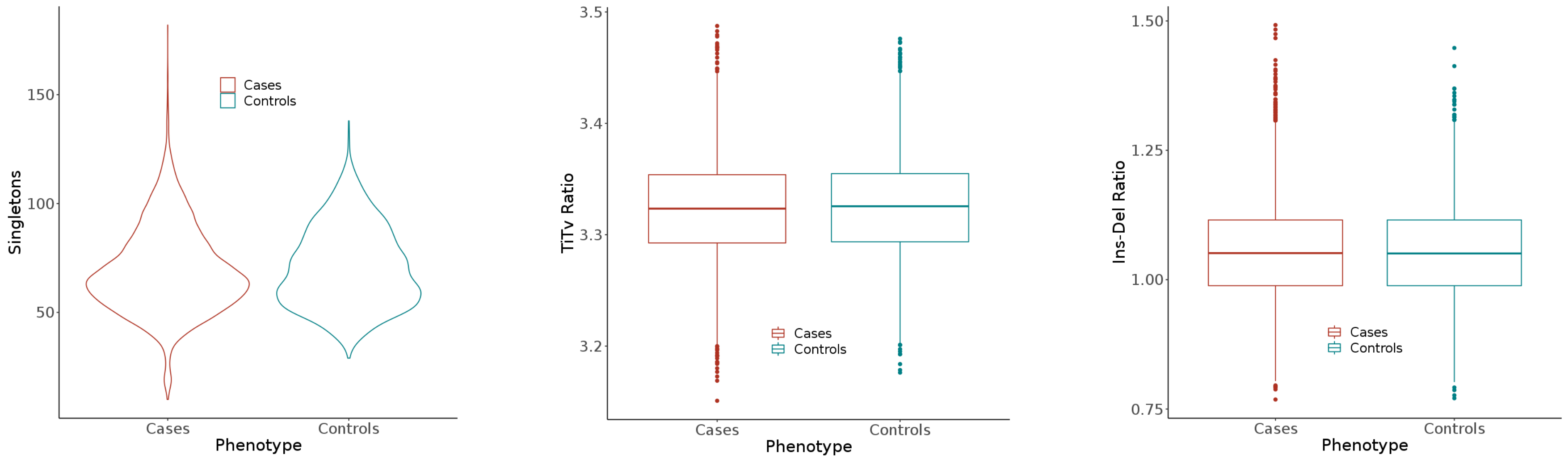

**Fig. S6: Variant counts and calling metrics in the final sample set.**

**A. Variant calling metrics stratified by phenotype.** The sample and variant homogenization steps resulted the removal of half of the raw variant calls in coding regions (see Fig. S1 for comparison) but also in a balanced SNP and Indel variant count distribution between the cases and controls. The residual difference in singletons count mirrors the European ancestry clusters seen in Fig. S5. SNVs: single nucleotide variations. Indels: insertions and deletions. TiTv: transitions-transversions. Ins-Del: insertions-deletions.

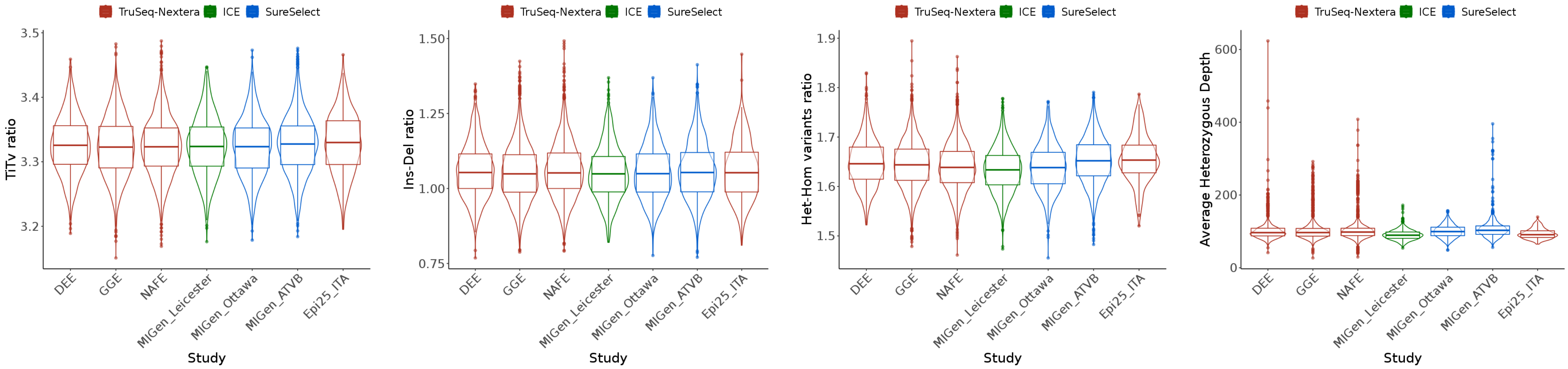

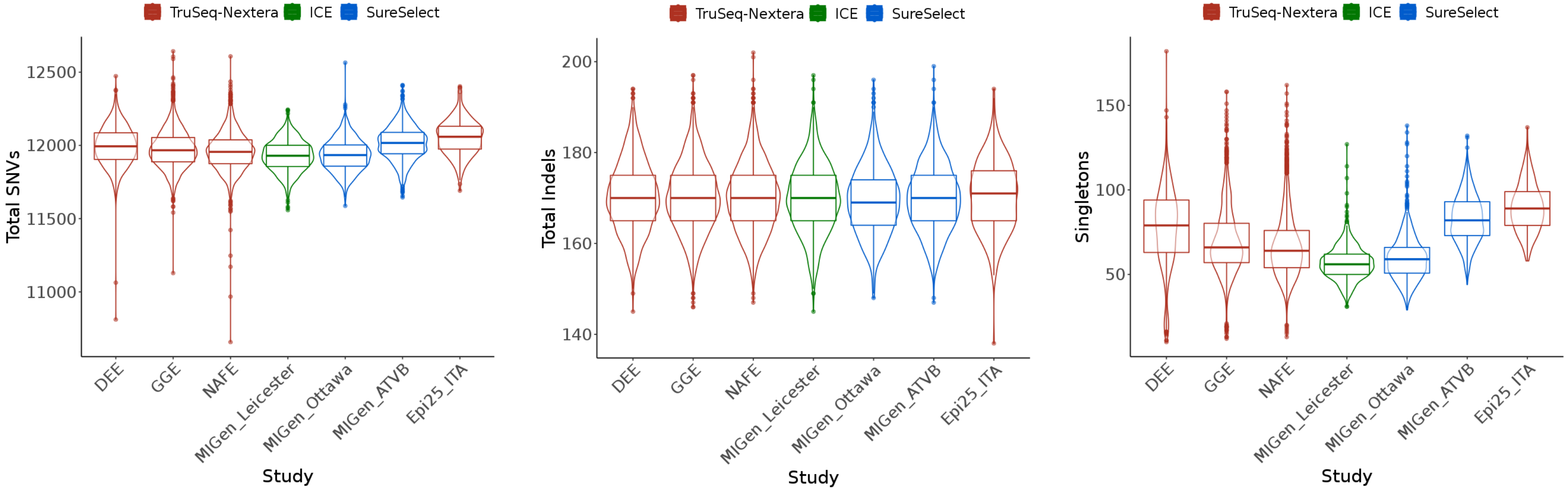

**Fig. S6: Variant counts and calling metrics in the final sample set.**

**B. Variant calling metrics stratified by enrichment kit and cohorts.** The residual differences in variant calling metrics between capture kits are minimal. Metrics that reflect the differences in ancestry as well (variant and singleton counts; bottom panels) were included as covariates in the regression analysis. SNVs: single nucleotide variations. Indels: insertions and deletions. TiTv: transitions-transversions. Ins-Del: insertions-deletions. Het-Hom: heterozygous-homozygous.

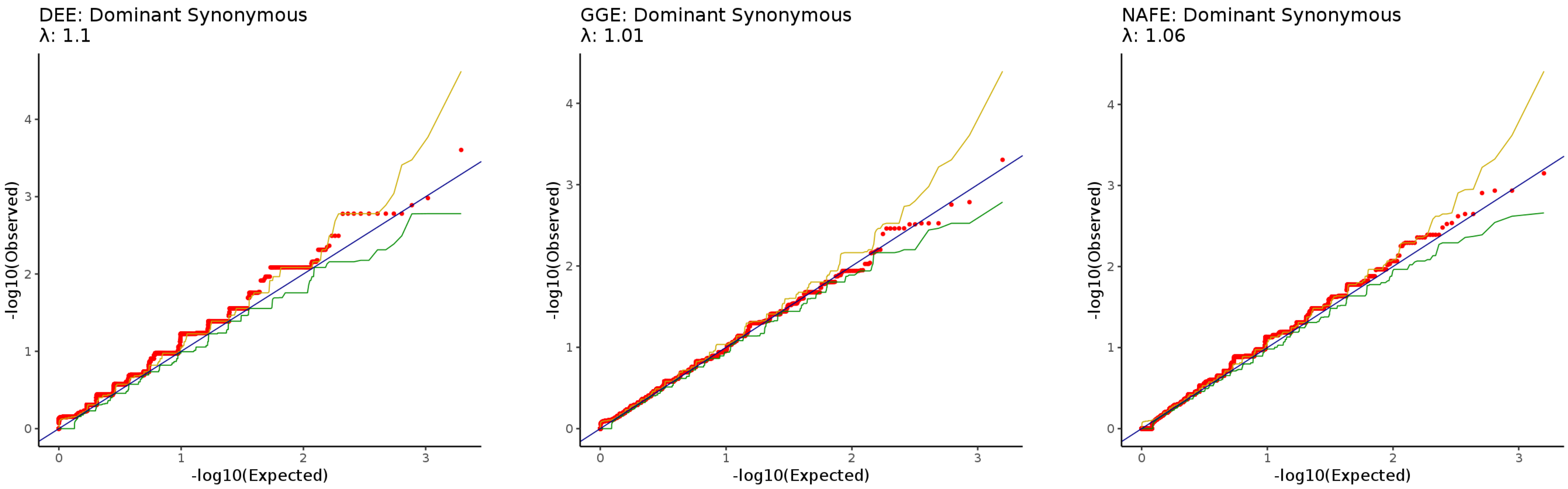

**Fig. S7: Quantile-Quantile plots of gene collapsing analysis of ultra-rare synonymous variants.** Observed *p* values are obtained from testing the significance of the difference in qualifying and unqualifying cases and controls counts (cases and controls with or without qualifying variants) using Fisher Exact Test. Expected *p* values indicate the mean *p* values obtained from 1000 permutations of sample labels followed by Fisher Exact Test. Green and golden lines indicate 2.5$^{th}$ and 97.5$^{th}$ centiles of permutation *p* values. Genomic Inflation Factor estimates (λ) were calculated from a comparison of the observed and mean permutation *p* values. DEE: developmental and epileptic encephalopathies. GGE: genetic generalized epilepsies. NAFE: non-acquired focal epilepsies.

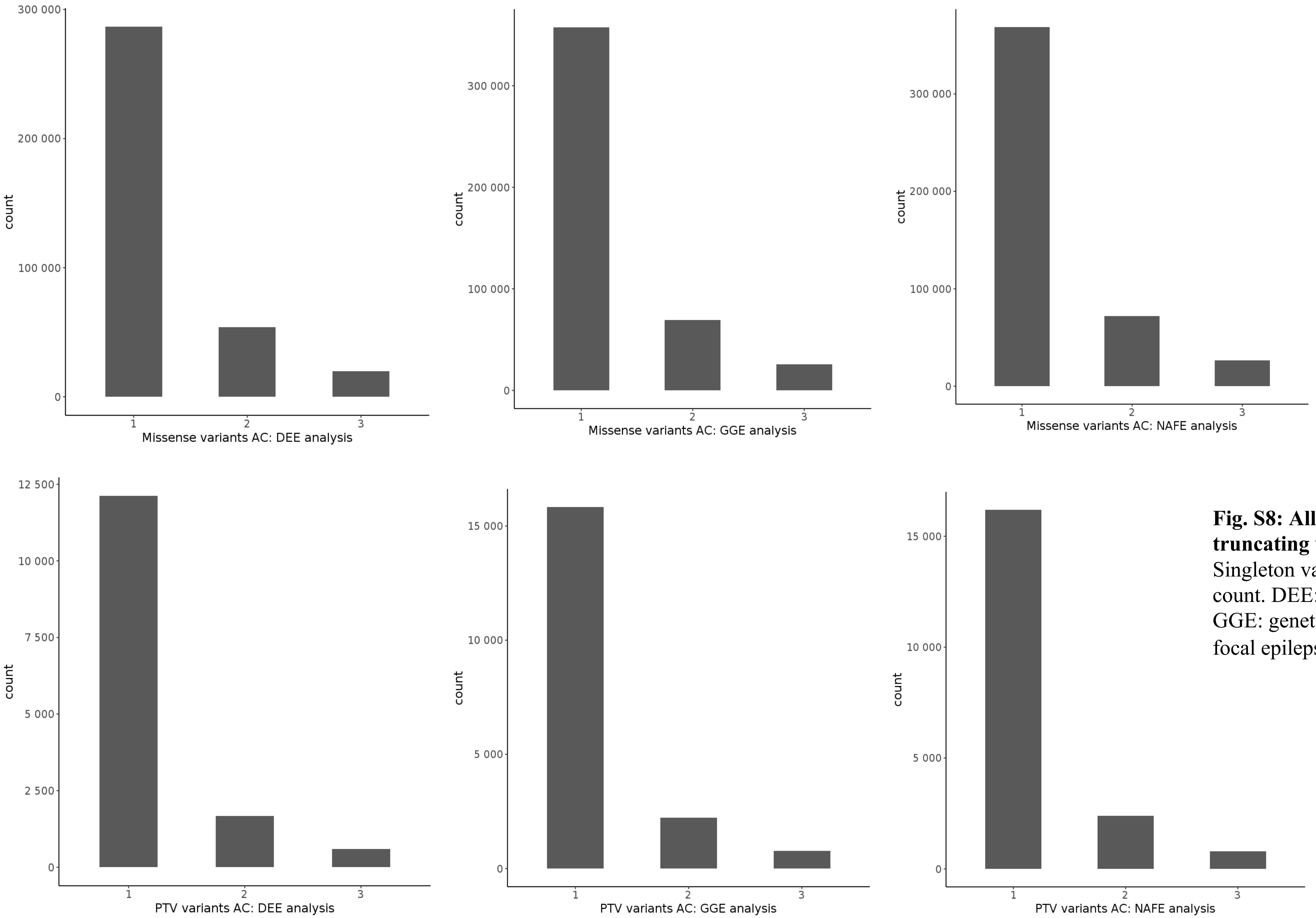

**Fig. S8: Allele counts of ultra-rare missense and protein truncating variants (PTV) observed in the study cohorts.** Singleton variants constitute most observations. AC: allele count. DEE: developmental and epileptic encephalopathies. GGE: genetic generalized epilepsies. NAFE: non-acquired focal epilepsies.

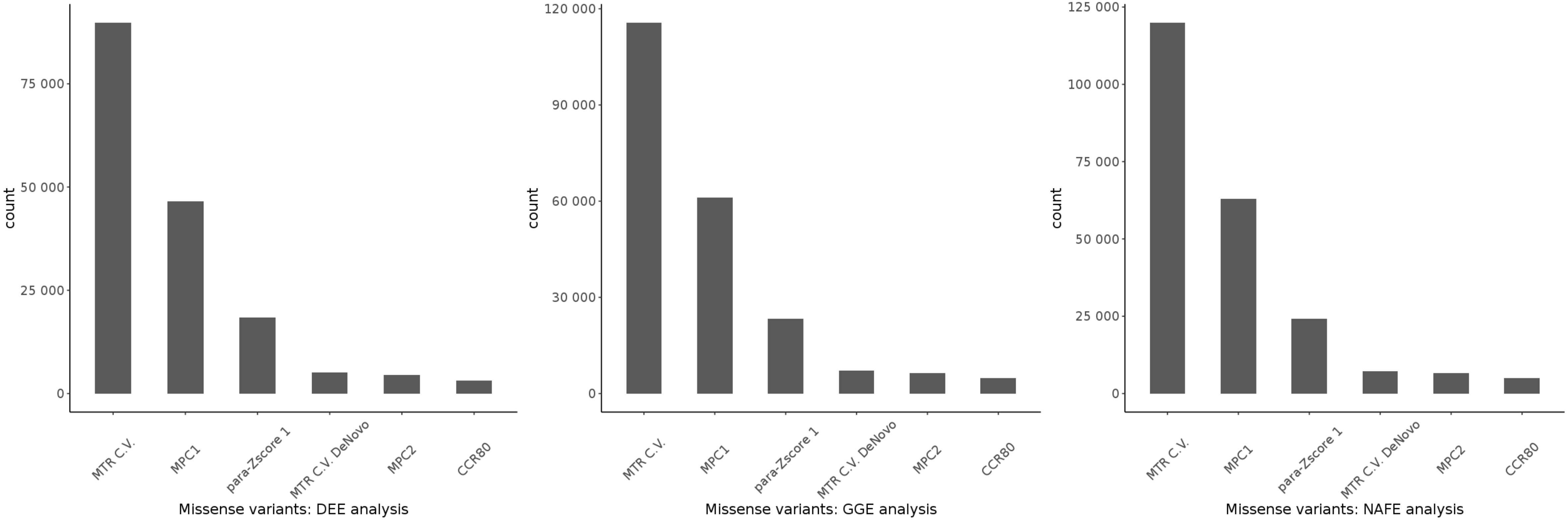

**Fig. S9: Ultra-rare variants count in selected missense conditions.** The variants are partially overlapping between these models, particularly because the same set of controls is used. DEE: developmental and epileptic encephalopathies. GGE: genetic generalized epilepsies. NAFE: non-acquired focal epilepsies. C.V.: ClinVar.

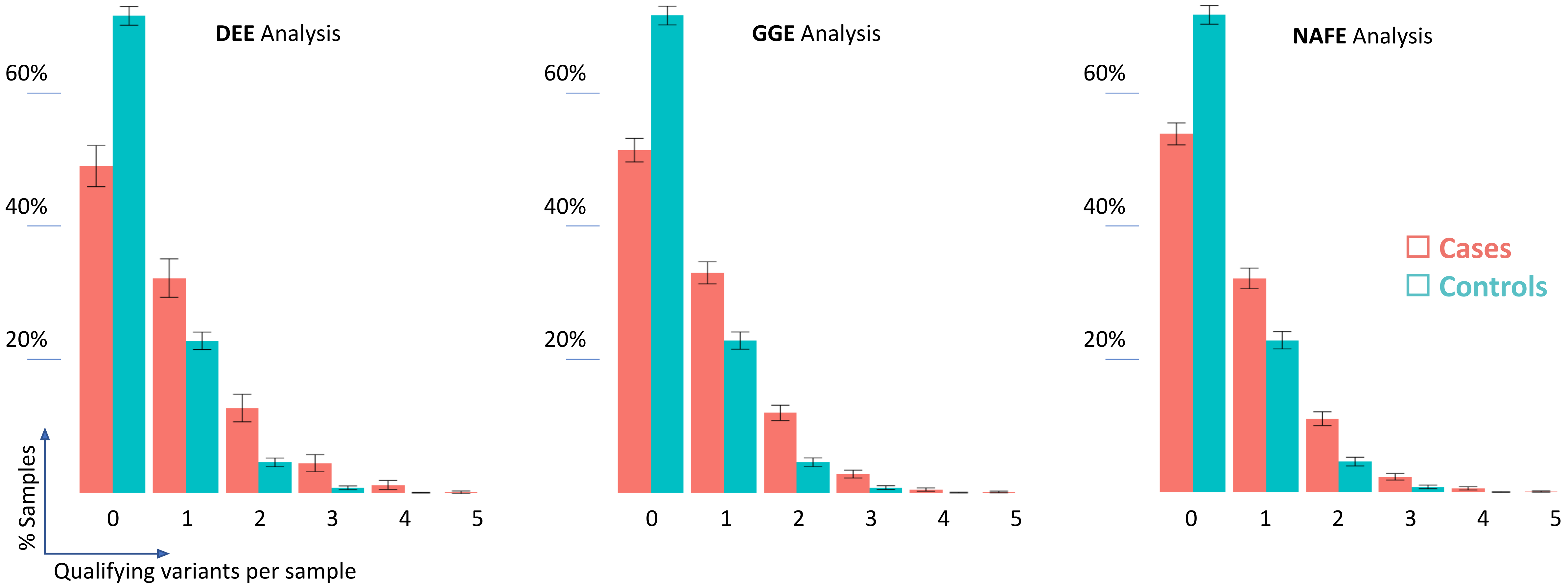

**Fig. S10: Variants load in three analysis sets in CCR80 condition.** Roughly, half of the cases compared to one fourth of the controls harbor one or more qualifying variant per exome in highly constrained sites. Error bars indicate the 95% confidence intervals calculated as follows: $p \pm 1.96 \times \sqrt{p(1-p)/n}$ where p is the proportion of samples and n is the total number of samples. DEE: developmental and epileptic encephalopathies. GGE: genetic generalized epilepsies. NAFE: non-acquired focal epilepsies.

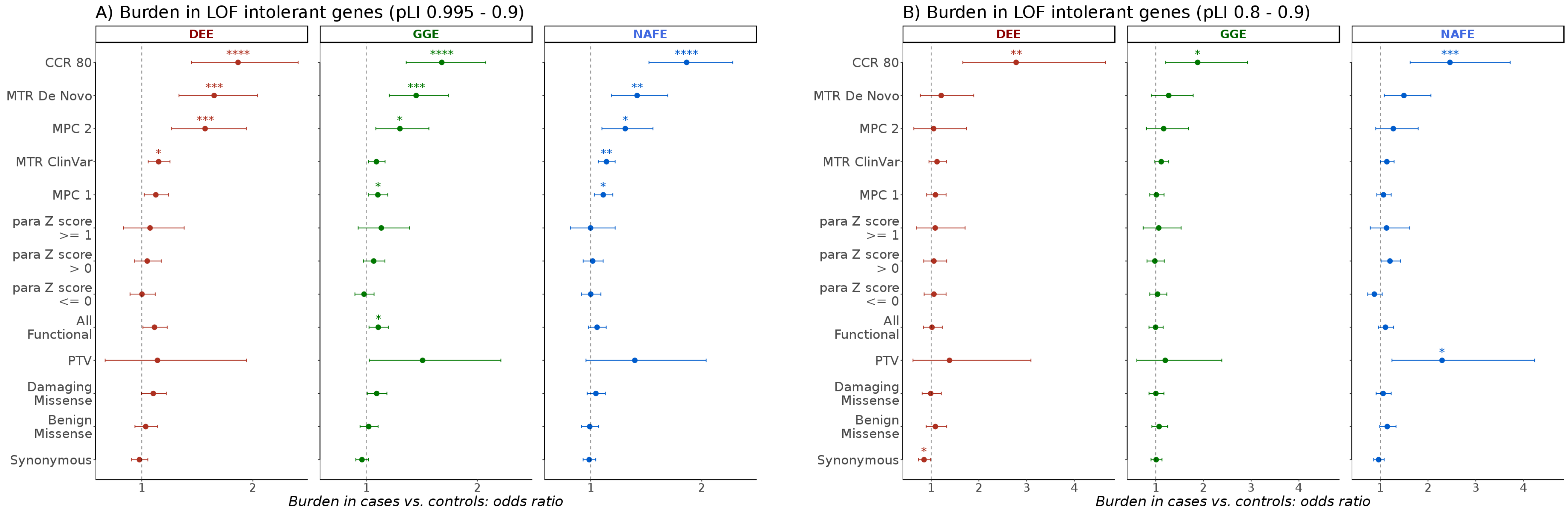

**Fig. S11: Burden of ultra-rare variants in loss-of-function intolerant genes.** *y* axis: variant conditions. *x* axis: odds ratio from regression analysis of individual burden of qualifying variants. Stars indicate FDR-adjusted *p* values: * < 0.05, ** < 0.005, *** < 0.0005, **** < 0.00005. Error bars indicate 95% confidence intervals of odds. DEE: developmental and epileptic encephalopathies. GGE: genetic generalized epilepsies. NAFE: non-acquired focal epilepsies. pLI: probability of loss-of-function intolerance.

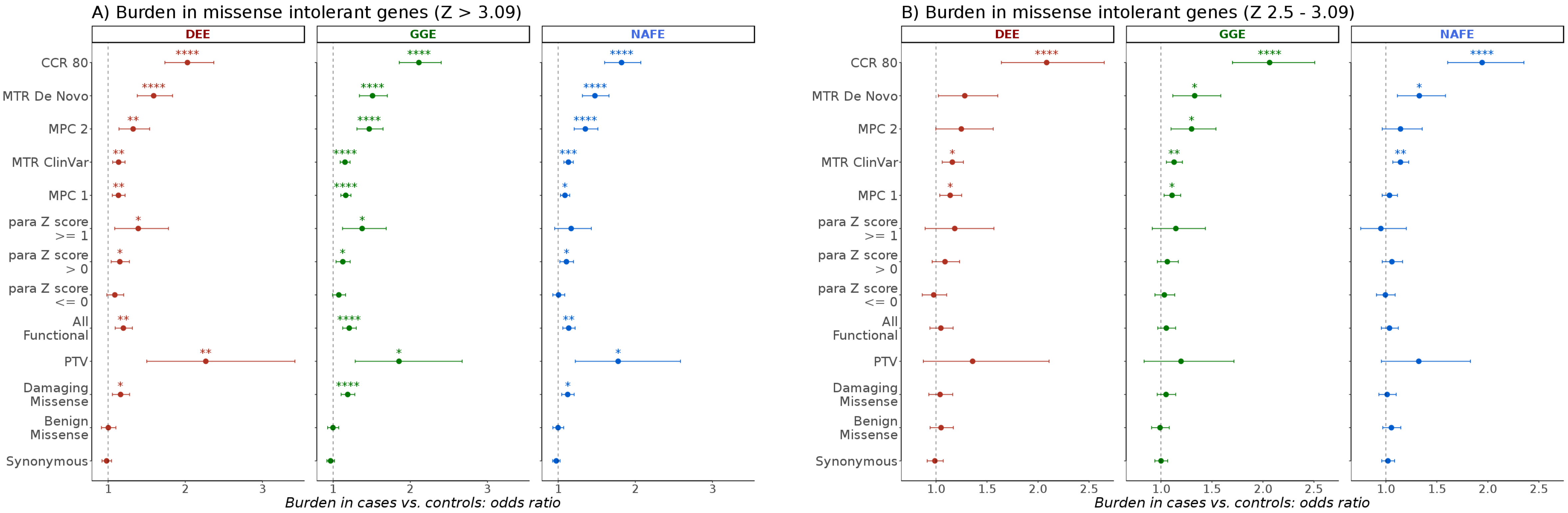

**Fig. S12: Burden in brain-expressed missense intolerant genes.** *y* axis: variant conditions. *x* axis: odds ratio from regression analysis of individual burden of qualifying variants. Stars indicate FDR-adjusted *p* values: * < 0.05, ** < 0.005, *** < 0.0005, **** < 0.00005. Error bars indicate 95% confidence intervals of odds. DEE: developmental and epileptic encephalopathies. GGE: genetic generalized epilepsies. NAFE: non-acquired focal epilepsies. Z: z-score of the probability of missense intolerance.

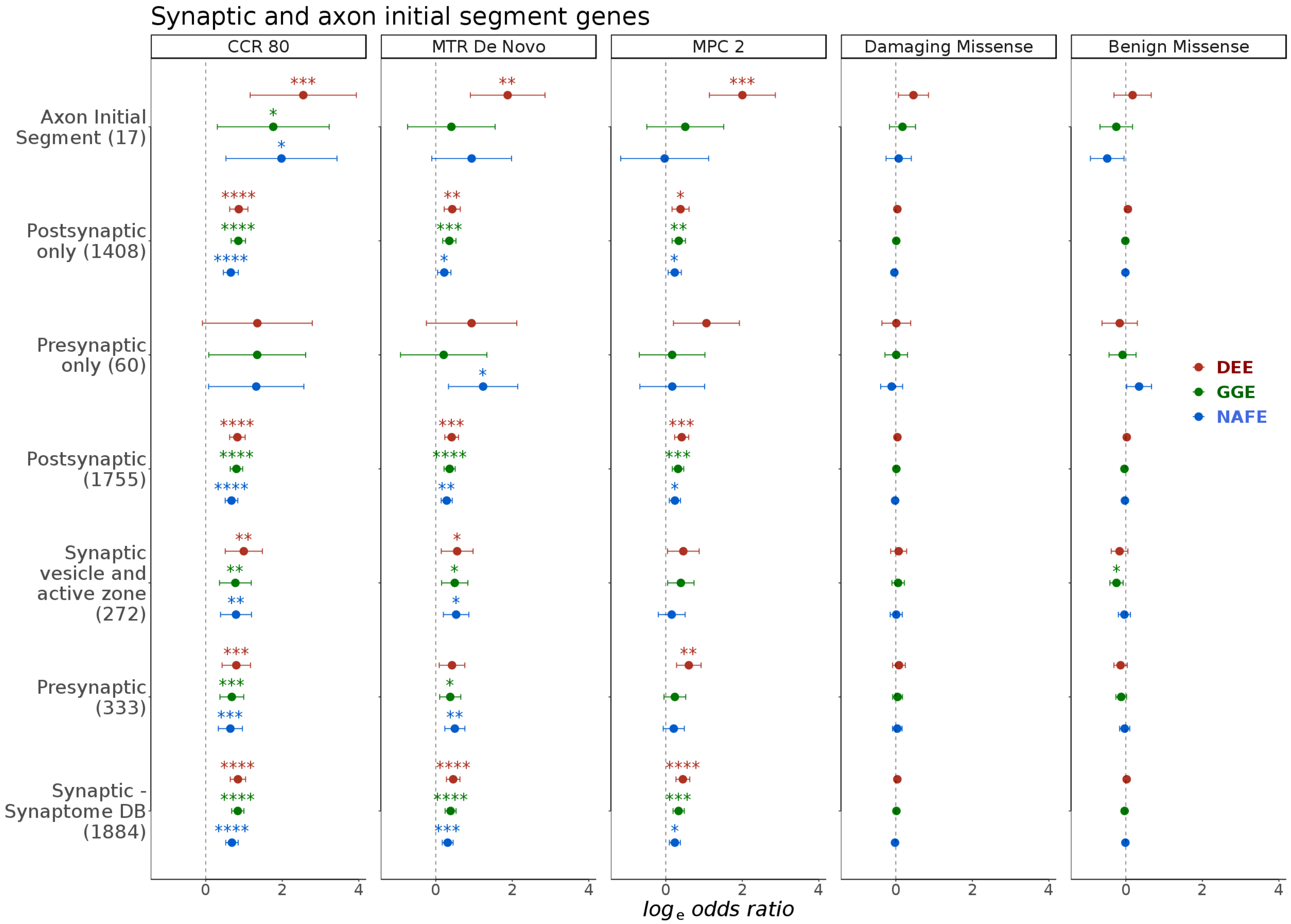

**Fig. S13: Burden in groups of axon initial segment and synaptic genes.** Panels: variant conditions. *y* axis: gene-sets (genes count between parenthesis). *x* axis: log odds ratio from regression analysis of individual burden of qualifying variants. Stars indicate FDR-adjusted *p* values: * < 0.05, ** < 0.005, *** < 0.0005, **** < 0.00005. Error bars indicate 95% confidence intervals of odds. DEE: developmental and epileptic encephalopathies. GGE: genetic generalized epilepsies. NAFE: non-acquired focal epilepsies.

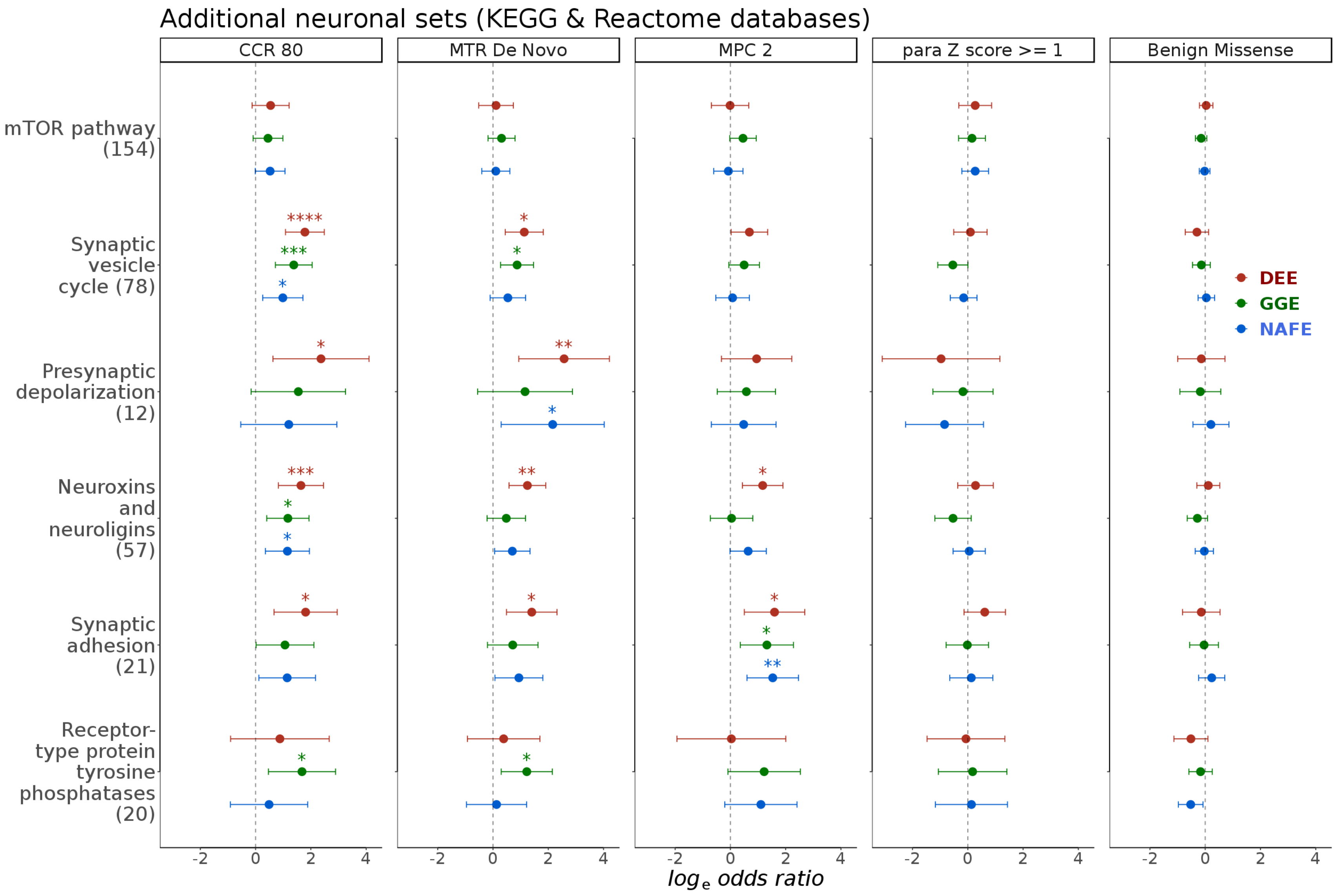

**Fig. S14: Burden in neuronal gene groups from KEGG and Reactome.** Panels: variant conditions. *y* axis: gene-sets (genes count between parenthesis). *x* axis: log odds ratio from regression analysis of individual burden of qualifying variants. Stars indicate FDR-adjusted *p* values: * < 0.05, ** < 0.005, *** < 0.0005, **** < 0.00005. Error bars indicate 95% confidence intervals of odds. DEE: developmental and epileptic encephalopathies. GGE: genetic generalized epilepsies. NAFE: non-acquired focal epilepsies.

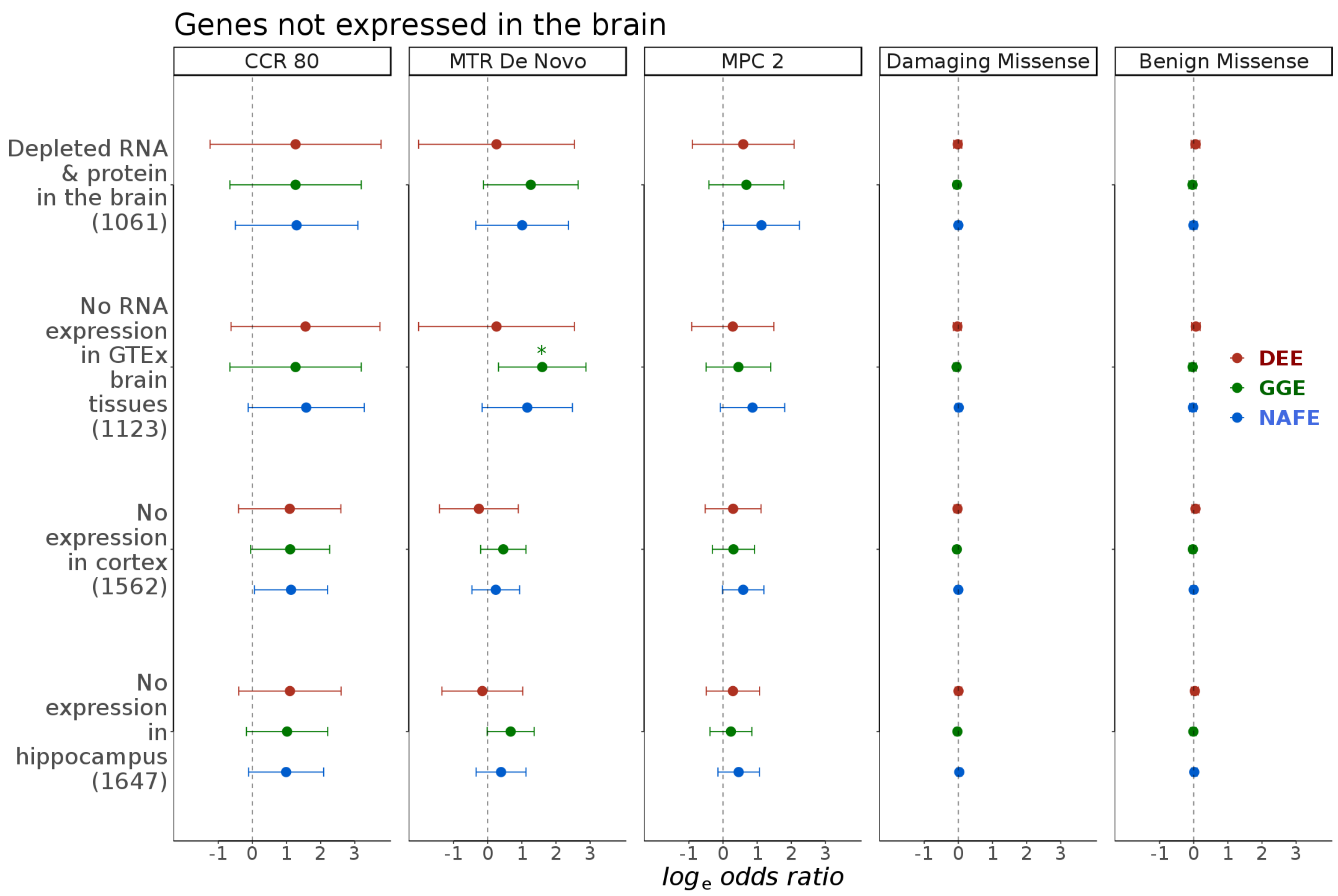

**Fig. S15: Burden in groups of genes not expressed in the brain.** Panels: variant conditions. *y* axis: gene-sets (genes count between parenthesis). *x* axis: log odds ratio from regression analysis of individual burden of qualifying variants. Stars indicate FDR-adjusted *p* values: * < 0.05, ** < 0.005, *** < 0.0005, **** < 0.00005. Error bars indicate 95% confidence intervals of odds. DEE: developmental and epileptic encephalopathies. GGE: genetic generalized epilepsies. NAFE: non-acquired focal epilepsies.

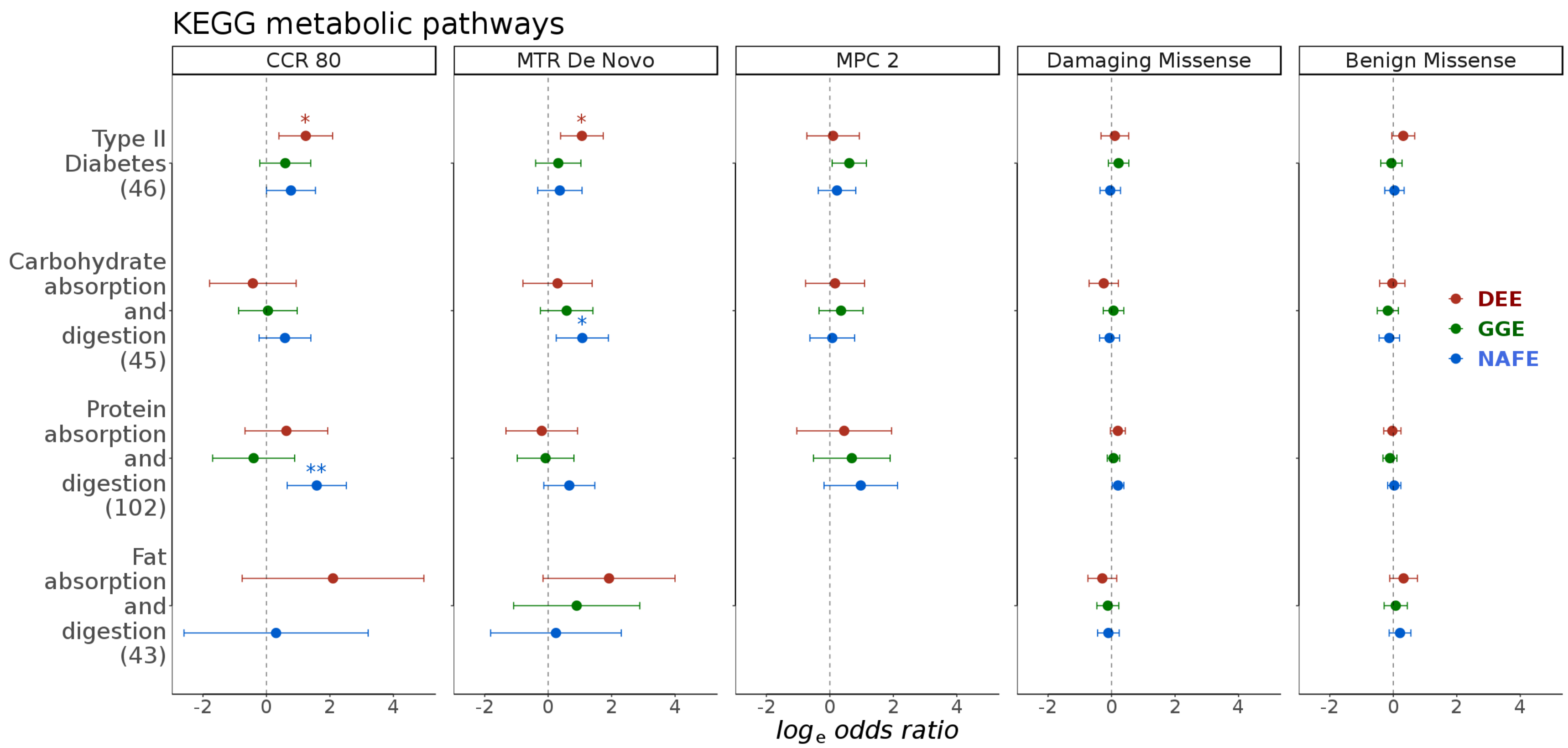

**Fig. S16: Burden in gene groups from KEGG metabolic pathways.** Panels: variant conditions. *y* axis: gene-sets (genes count between parenthesis). *x* axis: log odds ratio from regression analysis of individual burden of qualifying variants. Stars indicate FDR-adjusted *p* values: * < 0.05, ** < 0.005, *** < 0.0005, **** < 0.00005. Error bars indicate 95% confidence intervals of odds. DEE: developmental and epileptic encephalopathies. GGE: genetic generalized epilepsies. NAFE: non-acquired focal epilepsies. Missing odds and error bars indicate the lack of sufficient variant counts for the logistic fits to converge or that the intervals are too wide to plot on the current scale.

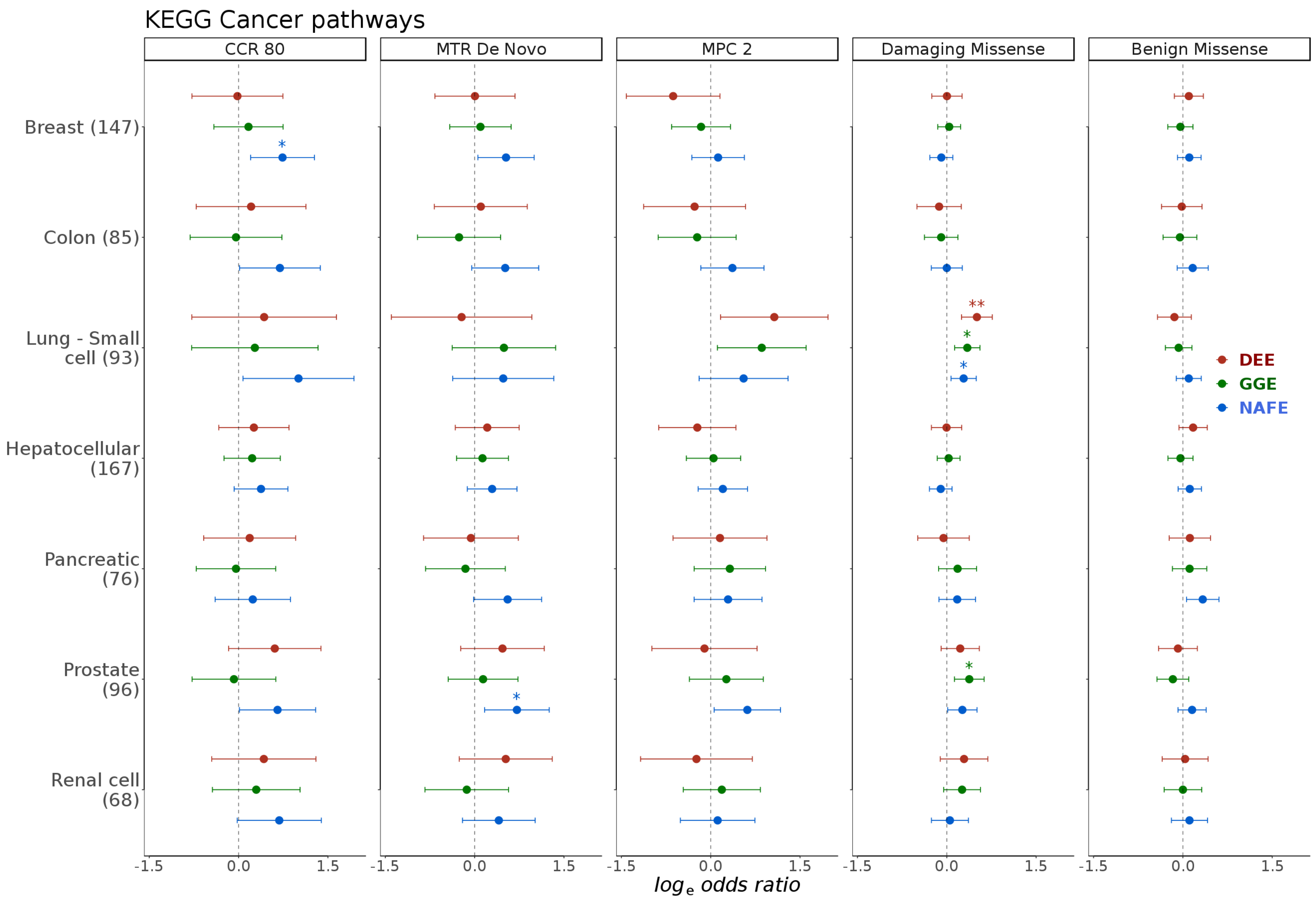

**Fig. S17: Burden in gene groups from KEGG cancer pathways.** Panels: variant conditions. *y* axis: gene-sets (genes count between parenthesis). *x* axis: log odds ratio from regression analysis of individual burden of qualifying variants. Stars indicate FDR-adjusted *p* values: * < 0.05, ** < 0.005, *** < 0.0005, **** < 0.00005. Error bars indicate 95% confidence intervals of odds. DEE: developmental and epileptic encephalopathies. GGE: genetic generalized epilepsies. NAFE: non-acquired focal epilepsies.

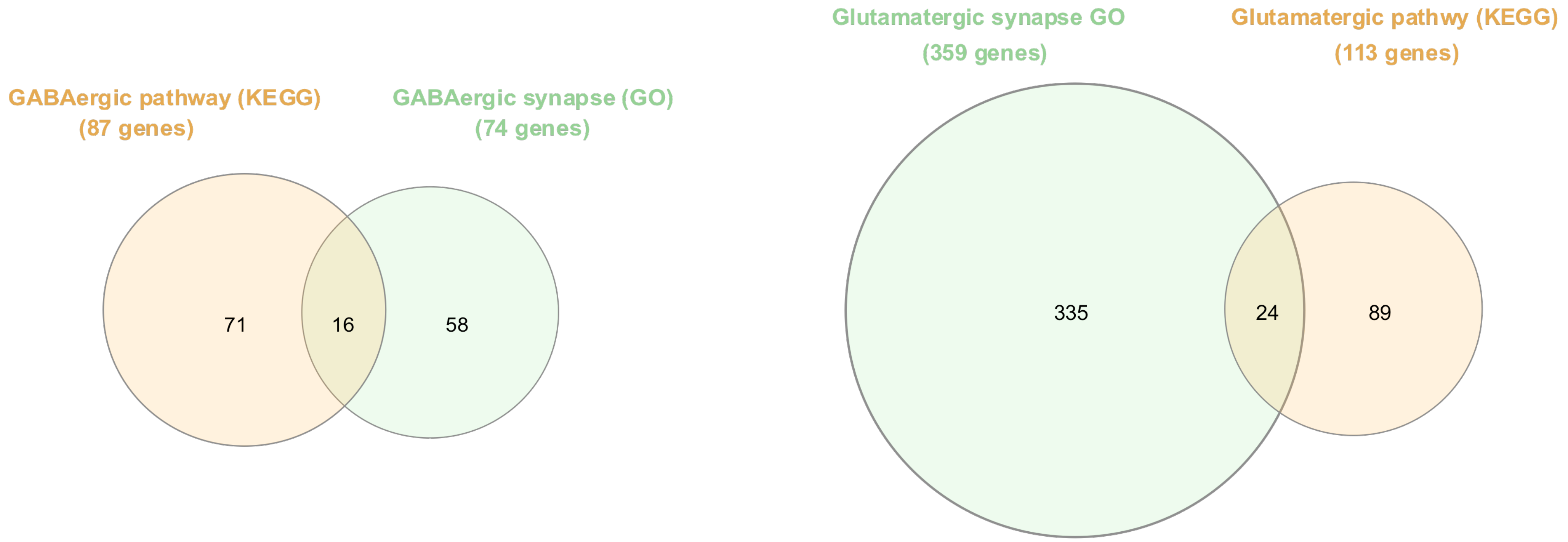

**Fig. S18: Overlap between gene groups representing the GABAergic and glutamatergic pathways (KEGG) and synapses (Gene Ontology).** GO: Gene Ontology.

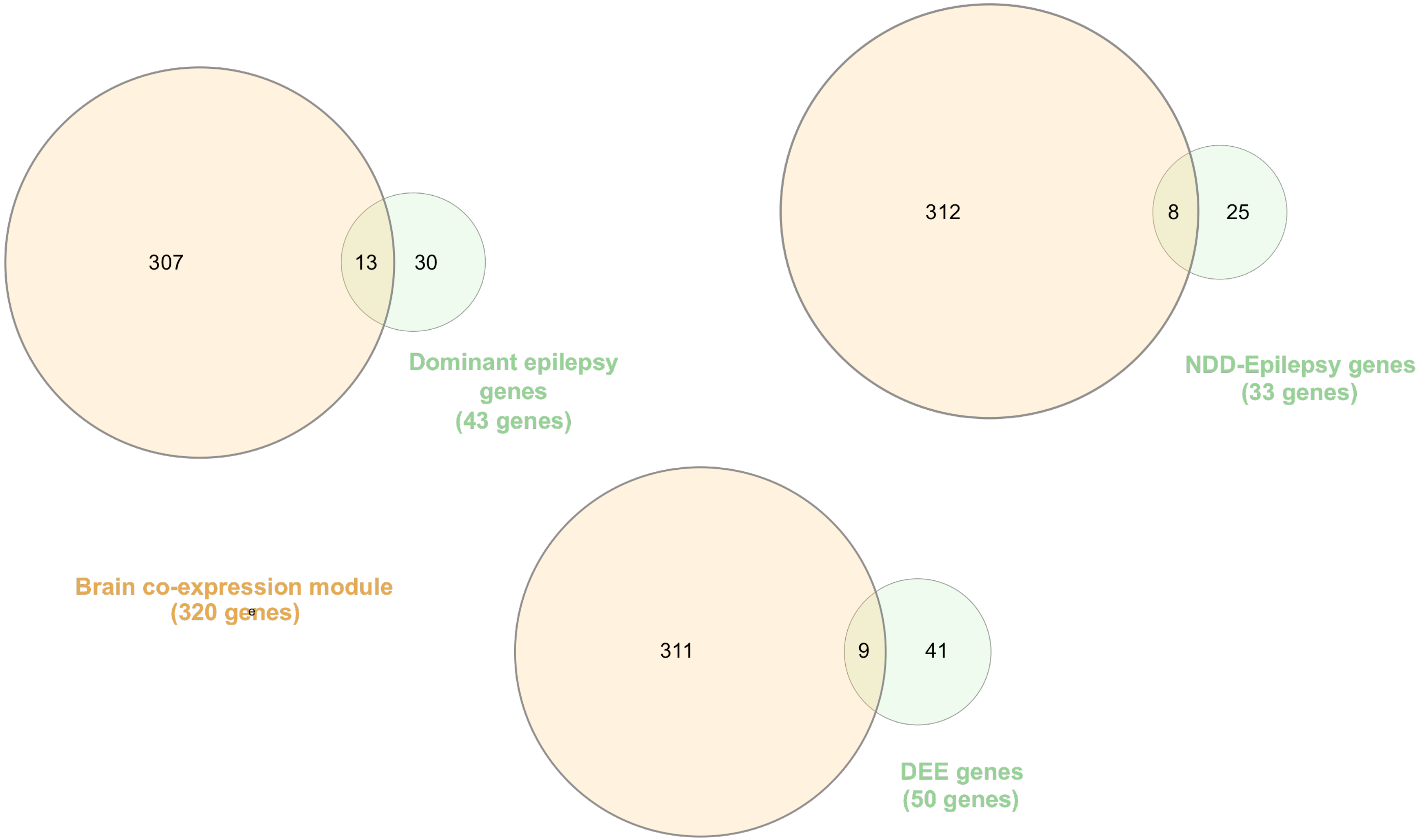

**Fig. S19: Overlap between an epilepsy-related co-expression module and groups representative of known disease genes.** The overlap is shown with three groups: Dominant epilepsy, developmental and epileptic encephalopathy (DEE) and neurodevelopmental disorders (NDD) with epilepsy disease genes.

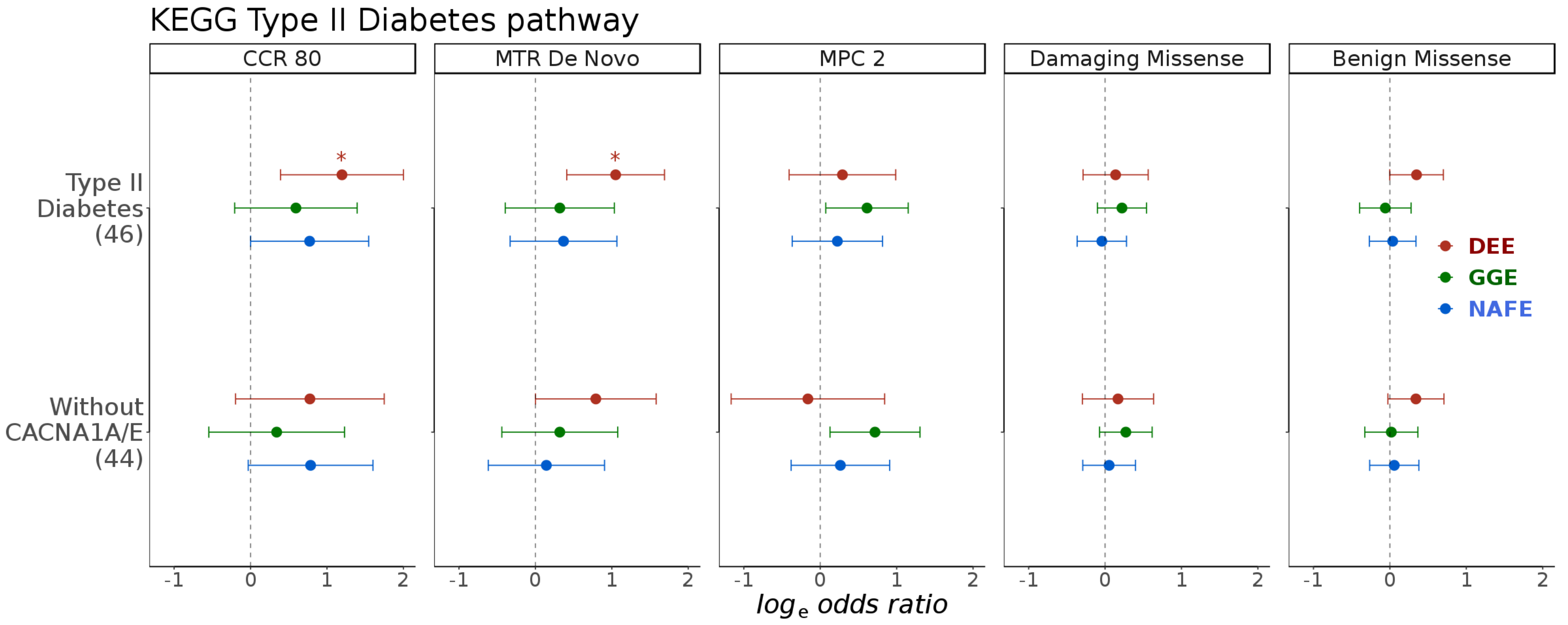

**Fig. S20: Burden in KEGG Type II Diabetes pathway genes with and without *CACNA1A/E*.** Panels: variant conditions. *y* axis: gene-sets (genes count between parenthesis). *x* axis: log odds ratio from regression analysis of individual burden of qualifying variants. Stars indicate FDR-adjusted *p* values: * < 0.05, ** < 0.005, *** < 0.0005, **** < 0.00005. Error bars indicate 95% confidence intervals of odds. DEE: developmental and epileptic encephalopathies. GGE: genetic generalized epilepsies. NAFE: non-acquired focal epilepsies.

**Epi25 Collaborative:**

**Epi25 sequencing, analysis, project management, and browser development at the Broad Institute:**

Yen-Chen Anne Feng[1-4], Daniel P. Howrigan[1,3,4], Liam E. Abbott[1,3,4], Katherine Tashman[1,3,4], Felecia Cerrato[3], Tarjinder Singh[1,3,4], Henrike Heyne[1,3,4], Andrea E. Byrnes[1,3,4], Claire Churchhouse[1,3,4], Nick Watts[1,3], Matthew Solomonson[1,3], Dennis Lal[4,5], Namrata Gupta[3], Stacey B. Gabriel[146], Mark J. Daly[1,3,4], Eric S. Lander[146,147,148], Benjamin M. Neale[1,3,4]

**Epi25 executive committee:**

Samuel F. Berkovic[9], Holger Lerche[8], David B. Goldstein[6], Daniel H. Lowenstein[7]

**Epi25 strategy, phenotyping, analysis, informatics, and project management committees:**

Samuel F. Berkovic[9], Holger Lerche[8], David B. Goldstein[6], Daniel H. Lowenstein[7], Gianpiero L. Cavalleri[67,70], Patrick Cossette[106], Chris Cotsapas[111], Peter De Jonghe[12-14], Tracy Dixon-Salazar[112], Renzo Guerrini[82], Hakon Hakonarson[101], Erin L. Heinzen[6], Ryan S. Dhindsa[6], Kate E. Stanley[6], Ingo Helbig[28,29,102], Patrick Kwan[10,11], Anthony G. Marson[58], Slavé Petrovski[11,116], Sitharthan Kamalakaran[6], Sanjay M. Sisodiya[57], Randy Stewart[113], Sarah Weckhuysen[12-14], Chantal Depondt[15], Dennis J. Dlugos[101], Ingrid E. Scheffer[9], Pasquale Striano[74], Catharine Freyer[7], Roland Krause[114], Patrick May[114], Kevin McKenna[7], Brigid M. Regan[9], Susannah T. Bellows[9], Costin Leu[4,5,57]

**Authors from individual Epi25 cohorts:**

**Australia: Melbourne (AUSAUS)**

Samuel F. Berkovic[9], Ingrid E. Scheffer[9], Brigid M. Regan[9], Caitlin A. Bennett[9], Susannah T. Bellows[9], Esther M.C. Johns[9], Alexandra Macdonald[9], Hannah Shilling[9], Rosemary Burgess[9], Dorien Weckhuysen[9], Melanie Bahlo[119,120]

**Australia: Royal Melbourne (AUSRMB)**

Terence J. O'Brien[10,11], Patrick Kwan[10,11], Slavé Petrovski[11,116], Marian Todaro[10,11]

**Belgium: Antwerp (BELATW)**

Sarah Weckhuysen[12-14], Hannah Stamberger[12-14], Peter De Jonghe[12-14]

**Belgium: Brussels (BELULB)**

Chantal Depondt[15]

**Canada: Andrade (CANUTN)**

Danielle M. Andrade[16,17], Tara R. Sadoway[17], Kelly Mo[17]

**Switzerland: Bern (CHEUBB)**

Heinz Kreste[118], Sabina Gallati[19]

**Cyprus (CYPCYP)**

Savvas S. Papacostas[20], Ioanna Kousiappa[20], George A. Tanteles[21]

**Czech Republic: Prague (CZEMTH)**

Katalin Štěrbová[22], Markéta Vlčková[23], Lucie Sedláčková[22], Petra Laššuthová[22]

**Germany: Frankfurt/Marburg (DEUPUM)**

Karl Martin Klein[24,25], Felix Rosenow[24,25], Philipp S. Reif[24,25], Susanne Knake[25]

**Germany: Bonn (DEUUKB)**

Wolfram S. Kunz[26,27], Gábor Zsurka[26,27], Christian E. Elger[27], Jürgen Bauer[27], Michael Rademacher[27]

**Germany: Kiel (DEUUKL)**

Ingo Helbig[28,29,102], Karl Martin Klein[24,25], Manuela Pendziwiat[29], Hiltrud Muhle[29], Annika Rademacher[29], Andreas van Baalen[29], Sarah von Spiczak[29], Ulrich Stephani[29], Zaid Afawi[30], Amos D. Korczyn[31], Moien Kanaan[32], Christina Canavati[32], Gerhard Kurlemann[33], Karen Müller-Schlüter[34], Gerhard Kluger[35,36], Martin Häusler[37], Ilan Blatt[31,115]

**Germany: Leipzig (DEUULG)**

Johannes R. Lemke[38], Ilona Krey[38]

**Germany: Tübingen (DEUUTB)**

Holger Lerche[8], Yvonne G. Weber[8,151], Stefan Wolking[8], Felicitas Becker[8,39], Christian Hengsbach[8], Sarah Rau[8], Ana F. Maisch[8], Bernhard J. Steinhoff[40], Andreas Schulze-Bonhage[41], Susanne Schubert-Bast[42], Herbert Schreiber[43], Ingo Borggräfe[44], Christoph J. Schankin[45], Thomas Mayer[46], Rudolf Korinthenberg[47], Knut Brockmann[48], Gerhard Kurlemann[33], Dieter Dennig[49], Rene Madeleyn[50]

**Finland: Kuopio (FINKPH)**

Reetta Kälviäinen[51], Pia Auvinen[51], Anni Saarela[51]

**Finland: Helsinki (FINUVH)**

Tarja Linnankivi[52], Anna-Elina Lehesjoki[53]

**Wales: Swansea (GBRSWU)**

Mark I. Rees[54,55], Seo-Kyung Chung[54,55], William O. Pickrell[54], Robert Powell[54,56]

**UK: UCL (GBRUCL)**

Sanjay M. Sisodiya[57,] Natascha Schneider[57], Simona Balestrini[57], Sara Zagaglia[57], Vera Braatz[57]

**UK: Imperial/Liverpool (GBRUNL)**

Anthony G. Marson[58], Michael R. Johnson[59], Pauls Auce[60], Graeme J. Sills[61]

**Hong Kong (HKGHKK)**

Patrick Kwan[10,11,62], Larry W. Baum[117,118,63], Pak C. Sham[117,118,63], Stacey S. Cherny[64], Colin H.T. Lui[65]

**Croatia (HRVUZG)**

Nina Barišić[66]

**Ireland: Dublin (IRLRCI)**

Gianpiero L. Cavalleri[67,70], Norman Delanty[67,70], Colin P. Doherty[68,70], Arif Shukralla[69], Mark McCormack[67], Hany El-Naggar[69,70]

**Italy: Milan (ITAICB)**

Laura Canafoglia[71], Silvana Franceschetti[71], Barbara Castellotti[72], Tiziana Granata[73]

**Italy: Genova (ITAIGI)**

Pasquale Striano[74], Federico Zara[75], Michele Iacomino[75], Francesca Madia[75], Maria Stella Vari[74], Maria Margherita Mancardi[75], Vincenzo Salpietro[74]

**Italy: Bologna (ITAUBG)**

Francesca Bisulli[76,77], Paolo Tinuper[76,77], Laura Licchetta[76,77], Tommaso Pippucci[78], Carlotta Stipa[79], Raffaella Minardi[76]

**Italy: Catanzaro (ITAUMC)**

Antonio Gambardella[80], Angelo Labate[80], Grazia Annesi[81], Lorella Manna[81], Monica Gagliardi[81]

**Italy: Florence (ITAUMR)**

Renzo Guerrini[82], Elena Parrini[82], Davide Mei[82], Annalisa Vetro[82], Claudia Bianchini[82], Martino Montomoli[82], Viola Doccini[82], Carla Marini[82]

**Japan: RIKEN Institute (JPNRKI)**

Toshimitsu Suzuki[83], Yushi Inoue[84], Kazuhiro Yamakawa[83]

**Lithuania (LTUUHK)**

Birute Tumiene[85,86]

**New Zealand: Otago (NZLUTO)**

Lynette G. Sadleir[87], Chontelle King[87], Emily Mountier[87]

**Turkey: Bogazici (TURBZU)**

S. Hande Caglayan[88], Mutluay Arslan[89], Zuhal Yapıcı[90], Uluc Yis[91], Pınar Topaloglu[90], Bulent Kara[92], Dilsad Turkdogan[93], Aslı Gundogdu-Eken[88]

**Turkey: Istanbul (TURIBU)**

Nerses Bebek[94,95], Sibel Uğur-İşeri[95], Betül Baykan[94], Barış Salman[95], Garen Haryanyan[94], Emrah Yücesan[149], Yeşim Kesim[94], Çiğdem Özkara[96]

**USA: BCH (USABCH)**

Annapurna Poduri[97,98], Beth R. Shiedley[97,98], Catherine Shain[97,98]

**USA: Philadelphia/CHOP (USACHP) and Philadelphia/Rowan (USACRW)**

Russell J. Buono[99,100,101], Thomas N. Ferraro[99,102], Michael R. Sperling[100], Dennis J. Dlugos[101,102], Warren Lo[103], Michael Privitera[104], Jacqueline A. French[105], Patrick Cossette[106], Steven Schachter[107], Hakon Hakonarson[101]

**USA: EPGP (USAEGP)**

Daniel H. Lowenstein[7], Ruben I. Kuzniecky[108], Dennis J. Dlugos[101,102], Orrin Devinsky[105]

**USA: NYU HEP (USAHEP)**

Daniel H. Lowenstein[7], Ruben I. Kuzniecky[108], Jacqueline A. French[105], Manu Hegde[7]

**USA: Penn/CHOP (USAUPN)**

Ingo Helbig[28,102], Pouya Khankhanian[109,110], Katherine L. Helbig[28], Colin A. Ellis[110]

**Italian controls**

Gianfranco Spalletta[121,122], Fabrizio Piras[121], Federica Piras[121], Tommaso Gili[123,121], Valentina Ciullo[121,124]

**German controls**

Andreas Reif[125,126]

**UK/IRL controls 1**

Andrew McQuillin[127], Nick Bass[127]

**UK/IRL controls 2**

Andrew McIntosh[128], Douglas Blackwood[128], Mandy Johnstone[128]

**FINRISK controls**

Aarno Palotie[1,2,4,129,130]

**Genomic Psychiatry Cohort (GPC) controls**

Michele T. Pato[131], Carlos N. Pato[131], Evelyn J. Bromet[132], Celia Barreto Carvalho[133], Eric D. Achtyes[134], Maria Helena Azevedo[135], Roman Kotov[132], Douglas S. Lehrer[136], Dolores Malaspina[137], Stephen R. Marder[138], Helena Medeiros[131], Christopher P. Morley[139], Diana O. Perkins[140], Janet L Sobell[141], Peter F. Buckley[142], Fabio Macciardi[143], Mark H. Rapaport[144], James A. Knowles[131], Genomic Psychiatry Cohort (GPC) Consortium, Ayman H. Fanous[131,145], Steven A. McCarroll[3,4,150]

**Affiliations:**

[1] Analytic and Translational Genetics Unit, Department of Medicine, Massachusetts General Hospital and Harvard Medical School, Boston, MA 02114, USA

[2] Psychiatric & Neurodevelopmental Genetics Unit, Department of Psychiatry, Massachusetts General Hospital and Harvard Medical School, Boston, MA 02114, USA

[3] Program in Medical and Population Genetics, Broad Institute of Harvard and MIT, 7 Cambridge Center, Cambridge, MA 02142, USA

[4] Stanley Center for Psychiatric Research, Broad Institute of Harvard and MIT, Cambridge, MA 02142, USA

[5] Genomic Medicine Institute, Cleveland Clinic, Cleveland, OH 44195, USA

[6] Institute for Genomic Medicine, Columbia University, New York, NY 10032, USA

[7] Department of Neurology, University of California, San Francisco, CA 94110, USA

[8] Department of Neurology and Epileptology, Hertie Institute for Clinical Brain Research, University of Tübingen, 72076 Tübingen, Germany

[9] Epilepsy Research Centre, Department of Medicine, University of Melbourne, Victoria, Australia
[10] Department of Neuroscience, Central Clinical School, Monash University, Alfred Hospital, Melbourne, Australia
[11] Departments of Medicine and Neurology, University of Melbourne, Royal Melbourne Hospital, Parkville, Australia
[12] Neurogenetics Group, Center for Molecular Neurology, VIB, Antwerp, Belgium
[13] Laboratory of Neurogenetics, Institute Born-Bunge, University of Antwerp, Belgium
[14] Division of Neurology, Antwerp University Hospital, Antwerp, Belgium
[15] Department of Neurology, Université Libre de Bruxelles, Brussels, Belgium
[16] Department of Neurology, Toronto Western Hospital, Toronto, ON M5T 2S8, Canada
[17] University Health Network, University of Toronto, Toronto, ON, Canada
[18] Departments of Neurology and BioMedical Research, Bern University Hospital and University of Bern, Bern, Switzerland
[19] Institute of Human Genetics, Bern University Hospital, Bern, Switzerland
[20] Neurology Clinic B, The Cyprus Institute of Neurology and Genetics, 2370 Nicosia, Cyprus
[21] Department of Clinical Genetics, The Cyprus Institute of Neurology and Genetics, 2370 Nicosia, Cyprus
[22] Department of Paediatric Neurology, 2nd Faculty of Medicine, Charles University and Motol Hospital, Prague, Czech Republic
[23] Department of Biology and Medical Genetics, 2nd Faculty of Medicine, Charles University and Motol
Hospital, Prague, Czech Republic
[24] Epilepsy Center Frankfurt Rhine-Main, Center of Neurology and Neurosurgery, Goethe University Frankfurt, Frankfurt, Germany
[25] Epilepsy Center Hessen-Marburg, Department of Neurology, Philipps University Marburg, Marburg, Germany
[26] Institute of Experimental Epileptology and Cognition Research, University Bonn, 53127 Bonn, Germany
[27] Department of Epileptology, University Bonn, 53127 Bonn, Germany
[28] Division of Neurology, Children's Hospital of Philadelphia, Philadelphia, PA 19104, USA
[29] Department of Neuropediatrics, Christian-Albrechts-University of Kiel, 24105 Kiel, Germany
[30] Sackler School of Medicine, Tel-Aviv University, Ramat Aviv, Israel
[31] Tel-Aviv University Sackler Faculty of Medicine, Ramat Aviv 69978, Israel
[32] Hereditary Research Lab, Bethlehem University, Bethlehem, Palestine
[33] Department of Neuropediatrics, Westfälische Wilhelms-University, Münster, Germany
[34] Epilepsy Center for Children, University Hospital Neuruppin, Brandenburg Medical School, Neuruppin, Germany

[35] Neuropediatric Clinic and Clinic for Neurorehabilitation, Epilepsy Center for Children and Adolescents, Vogtareuth, Germany
[36] Research Institute Rehabilitation / Transition / Palliation, PMU Salzburg, Austria
[37] Division of Neuropediatrics and Social Pediatrics, Department of Pediatrics, University Hospital, RWTH Aachen, Aachen, Germany
[38] Institute of Human Genetics, Leipzig, Germany
[39] RKU-University Neurology Clinic of Ulm, Ulm, Germany
[40] Kork Epilepsy Center, Kehl-Kork, Germany
[41] Epilepsy Center, University of Freiburg, Freiburg im Breisgau, Germany
[42] Section Neuropediatrics and Inborn Errors of Metabolism, University Children's Hospital, Heidelberg, Germany
[43] Neurological Practice Center & Neuropoint Patient Academy, Ulm, Germany
[44] Department of Pediatric Neurology and Developmental Medicine, LMU Munich, Munich, Germany
[45] Department of Neurology, University of Munich Hospital-Großhadern, Munich, Germany
[46] Saxonian Epilepsy Center Radeberg, Radeberg, Germany
[47] Division of Neuropediatrics and Muscular Disorders, University Hospital Freiburg, Freiburg, Germany
[48] University Children's Hospital, Göttingen, Germany
[49] Private Neurological Practice, Stuttgart, Germany
[50] Department of Pediatrics, Filderklinik, Filderstadt, Germany
[51] Neurocenter, Kuopio University Hospital, Kuopio Finland and Institute of Clinical Medicine, University of Eastern Finland, Finland
[52] Child Neurology, University of Helsinki and Helsinki University Hospital, Helsinki, Finland
[53] Medicum, University of Helsinki, Helsinki, Finland and Folkhälsan Research Center, Helsinki, Finland
[54] Neurology Research Group, Swansea University Medical School, Swansea University SA2 8PP, UK
[55] Faculty of Medicine and Health, University of Sydney, Sydney, Australia
[56] Department of Neurology, Morriston Hospital, Abertawe BroMorgannwg HealthBoard, Swansea, UK
[57] Department of Clinical and Experimental Epilepsy, UCL Queen Square Institute of Neurology, London, UK and Chalfont Centre for Epilepsy, Chalfont St Peter, UK
[58] Department of Molecular and Clinical Pharmacology, University of Liverpool, Liverpool, UK
[59] Division of Brain Sciences, Imperial College London, London, UK
[60] Department of Neurology, Walton Centre NHS Foundation Trust, Liverpool, UK
[61] School of Life Sciences, University of Glasgow, Glasgow, UK
[62] Department of Medicine and Therapeutics, Chinese University of Hong Kong, Hong Kong, China
[63] Department of Psychiatry, University of Hong Kong, Hong Kong, China

[64] Department of Epidemiology and Preventive Medicine and Department of Anatomy and Anthropology, Sackler Faculty of Medicine, Tel Aviv University, Israel
[65] Department of Medicine, Tseung Kwan O Hospital, Hong Kong, China
[66] Department of Pediatric University Hospital centre Zagreb, Croatia
[67] The Department of Molecular and Cellular Therapeutics, The Royal College of Surgeons in Ireland, Dublin, Ireland
[68] Neurology Department, St. James Hospital, Dublin, Ireland
[69] The Department of Neurology, Beaumont Hospital, Dublin, Ireland
[70] The FutureNeuro Research Centre, Ireland
[71] Neurophysiopathology, Fondazione IRCCS Istituto Neurologico Carlo Besta, Milan, Italy
[72] Unit of Genetics of Neurodegenerative and Metabolic Diseases, Fondazione IRCCS Istituto Neurologico Carlo Besta, Milan, Italy
[73] Department of Pediatric Neuroscience, Fondazione IRCCS Istituto Neurologico Carlo Besta, Milan, Italy
[74] Pediatric Neurology and Muscular Diseases Unit, Department of Neurosciences, Rehabilitation, Ophthalmology, Genetics, Maternal and Child Health, University of Genoa, "G. Gaslini" Institute, Genova, Italy
[75] Laboratory of Neurogenetics, "G. Gaslini" Institute, Genova, Italy
[76] IRCCS, Institute of Neurological Sciences of Bologna, Bologna, Italy
[77] Department of Biomedical and Neuromotor Sciences, University of Bologna, Bologna, Italy
[78] Medical Genetics Unit, Polyclinic Sant'Orsola-Malpighi University Hospital, Bologna, Italy
[79] Department of Biomedical and Neuromotor Sciences, University of Bologna, Bologna, Italy
[80] Institute of Neurology, Department of Medical and Surgical Sciences, University “Magna Graecia”, Catanzaro, Italy
[81] Institute of Molecular Bioimaging and Physiology, CNR, Section of Germaneto, Catanzaro, Italy
[82] Pediatric Neurology, Neurogenetics and Neurobiology Unit and Laboratories, Children's Hospital A. Meyer, University of Florence, Italy
[83] Laboratory for Neurogenetics, RIKEN Center for Brain Science, Saitama, Japan
[84] National Epilepsy Center, Shizuoka Institute of Epilepsy and Neurological Disorder, Shizuoka, Japan
[85] Institute of Biomedical Sciences, Faculty of Medicine, Vilnius University, Vilnius, Lithuania
[86] Centre for Medical Genetics, Vilnius University Hospital Santaros Klinikos, Vilnius, Lithuania
[87] Department of Paediatrics and Child Health, University of Otago, Wellington
[88] Department of Molecular Biology and Genetics, Bogaziçi University, Istanbul, Turkey
[89] Department of Child Neurology, Gulhane Education and Research Hospital, Health Sciences University, Ankara, Turkey
[90] Department of Child Neurology, Istanbul Faculty of Medicine, Istanbul University, Istanbul, Turkey
[91] Department of Child Neurology, Medical School, Dokuz Eylul University, Izmir, Turkey

[92] Department of Child Neurology, Medical School, Kocaeli University, Kocaeli, Turkey
[93] Department of Child Neurology, Medical School, Marmara University, Istanbul, Turkey
[94] Department of Neurology, Istanbul Faculty of Medicine, Istanbul University, Istanbul, Turkey
[95] Department of Genetics, Aziz Sancar Institute of Experimental Medicine, Istanbul University, Istanbul, Turkey
[96] Department of Neurology, Faculty of Medicine, Cerrahpaşa University Istanbul, Istanbul, Turkey
[97] Epilepsy Genetics Program, Department of Neurology, Boston Children's Hospital, Boston, MA 02115, USA
[98] Department of Neurology, Harvard Medical School, Boston, MA 02115, USA
[99] Cooper Medical School of Rowan University, Camden, NJ 08103, USA
[100] Thomas Jefferson University, Philadelphia, PA 19107, USA
[101] The Children's Hospital of Philadelphia, Philadelphia, PA 19104, USA
[102] Perelman School of Medicine, University of Pennsylvania, PA 19104, USA
[103] Nationwide Children's Hospital, Columbus, OH 43205, USA
[104] University of Cincinnati, Cincinnati, OH 45220, USA
[105] Department of Neurology, New York University/Langone Health, New York, NY 10016, USA
[106] University of Montreal, Montreal, QC H3T 1J4, Canada
[107] Beth Israel Deaconess/Harvard, Boston, MA 02115, USA
[108] Department of Neurology, Hofstra-Northwell Medical School, New York, NY 11549, USA
[109] Center for Neuro-engineering and Therapeutics, University of Pennsylvania, Philadelphia, PA 19104, USA
[110] Department of Neurology, Hospital of University of Pennsylvania, Philadelphia, PA 19104, USA
[111] School of Medicine, Yale University, New Haven, CT 06510, USA
[112] LGS Foundation, NY 11716, USA
[113] National Institute of Neurological Disorders and Stroke, MD 20852, USA
[114] Luxembourg Centre for Systems Biomedicine, University Luxembourg, Esch-sur-Alzette, Luxembourg
[115] Department of Neurology, Sheba Medical Center, Ramat Gan, Israel
[116] Centre for Genomics Research, Precision Medicine and Genomics, IMED Biotech Unit, AstraZeneca, Cambridge, UK
[117] The State Key Laboratory of Brain and Cognitive Sciences, University of Hong Kong, Hong Kong, China
[118] Centre for Genomic Sciences, University of Hong Kong, Hong Kong, China
[119] Population Health and Immunity Division, the Walter and Eliza Hall Institute of Medical Research, Parkville 3052, VIC, Australia
[120] Department of Medical Biology, The University of Melbourne, Melbourne 3010, VIC, Australia
[121] Neuropsychiatry Laboratory, IRCCS Santa Lucia Foundation, Rome, Italy

[122] Division of Neuropsychiatry, Menninger Department of Psychiatry and Behavioral Sciences, Baylor College of Medicine, Houston, TX, USA
[123] IMT School for Advanced Studies Lucca, Lucca, Italy
[124] Department of Neurosciences, Psychology, Drug Research and Child Health, University of Florence, Florence, Italy
[125] Department of Psychiatry, Psychosomatic Medicine and Psychotherapy, University Hospital Frankfurt
[126] Department of Psychiatry, Psychotherapy and Psychosomatics, University Hospital Würzburg
[127] Division of Psychiatry, University College London, London, UK
[128] Division of Psychiatry, Centre for Clinical Brain Sciences, University of Edinburgh, Edinburgh, UK
[129] Institute for Molecular Medicine Finland, University of Helsinki, 00014, Finland
[130] Department of Neurology, Massachusetts General Hospital, Boston, MA, USA
[131] Department of Psychiatry and Behavioral Sciences, SUNY Downstate Medical Center, Brooklyn, NY, USA
[132] Department of Psychiatry, Stony Brook University, Stony Brook, NY, USA
[133] Faculty of Social and Human Sciences, University of Azores, PT
[134] Cherry Health and Michigan State University College of Human Medicine, Grand Rapids, MI, USA
[135] Institute of Medical Psychology, Faculty of Medicine, University of Coimbra, Coimbra, PT
[136] Department of Psychiatry, Wright State University, Dayton, OH, USA
[137] Departments of Psychiatry, Genetics & Genomics, Icahn School of Medicine at Mount Sinai, NY, USA
[138] Semel Institute for Neuroscience at UCLA, Los Angeles, CA, USA
[139] Departments of Public Health and Preventive Medicine, Family Medicine, and Psychiatry and Behavioral Sciences, State University of New York, Upstate Medical University, Syracuse, NY, USA
[140] Department of Psychiatry, University of North Carolina, Chapel Hill, NC, USA
[141] Department of Psychiatry & Behavioral Sciences, University of Southern California, Los Angeles, CA, USA
[142] Department of Psychiatry, Virginia Commonwealth University School of Medicine, Richmond, VA, USA
[143] Department of Psychiatry and Human Behavior, University of California, Irvine, CA, USA
[144] Department of Psychiatry and Behavioral Sciences, Emory University, Atlanta, GA, USA
[145] Department of Psychiatry, Veterans Administration New York Harbor Healthcare System, Brooklyn, NY, USA
[146] Broad Institute of MIT and Harvard, Cambridge, MA, USA
[147] Department of Biology, Massachusetts Institute of Technology, Cambridge, MA, USA
[148] Department of Systems Biology, Harvard Medical School, Boston, MA, USA
[149] Bezmialem Vakif University, Institute of Life Sciences and Biotechnology, Istanbul, Turkey

[150] Department of Genetics, Harvard Medical School, Boston, MA, USA

[151] Department of Neurosurgery, University of Tübingen, Tübingen, Germany
